# Rising together: Exploring Sourdough Fermentation Diversity through Co-design in the HealthFerm Citizen Science Initiative

**DOI:** 10.1101/2025.05.23.655785

**Authors:** Annina Meyer, Thomas Gettemans, Jan Patrick Tan, Fabio Tuccillo, Chiara Viretto, Iulia-Roxana Angelescu, Yamina De Bondt, Michelle Neugebauer, Ali Zein Alabiden Tlais, Fabio Cavelti, Luc De Vuyst, Marco Gobbetti, Christophe M. Courtin, Medana Zamfir, Rossana Coda, Laura Nyström, Stefan Weckx, Nicholas A. Bokulich

**Affiliations:** Laboratory of Food Systems Biotechnology, Department of Health Sciences and Technology, ETH Zurich, Schmelzbergstrasse 7, 8092 Zurich, Switzerland; Research Group of Industrial Microbiology and Food Biotechnology (IMDO), Faculty of Sciences and Bioengineering Sciences, Vrije Universiteit Brussel (VUB), Pleinlaan 2, B-1050 Brussels, Belgium; Laboratory of Food Biochemistry, Department of Health Sciences and Technology, ETH Zurich, Schmelzbergstrasse 9, 8092 Zurich, Switzerland; Department of Food and Nutrition, University of Helsinki, Agnes Sjöbergin katu 2, 00014 Helsinki, Finland; Faculty of Agricultural, Environmental and Food Sciences, Free University of Bolzano-Bozen, 39100 Bolzano, Italy; Institute of Biology Bucharest of the Romanian Academy, 296 Splaiul Independentei, Bucharest, Romania; Laboratory of Food Chemistry and Biochemistry and Leuven Food Science and Nutrition Research Centre (LFoRCe), KU Leuven, Kasteelpark Arenberg 20, B-3001 Leuven, Belgium

## Abstract

Fermented foods are culturally significant and increasingly recognized for their potential health benefits, yet scientific data on household fermentation practices remain limited. We launched a co-designed citizen science (CS) initiative within the HealthFerm project to collect sourdough fermentation data across Europe. Over 1000 participants from 33 countries registered, with 671 samples submitted, enabling large-scale analysis of fermentation practices, motivations, and sourdough characteristics. Participants also completed standardized at-home experiments and sensory evaluations, generating a dataset linking baking habits with physicochemical and sensory profiles. Distinct patterns emerged: professional bakers used older, more frequently refreshed starters and fermented at higher temperatures. Ingredient choices and motivations varied by country, shaped by perceived health benefits. Beyond data collection, this initiative established a microbial biobank and harmonized metadata resource, while offering practical insights into co-design, logistics, and public engagement. The resulting framework provides a transferable model for participatory research in microbiology and food systems science.

## Introduction

Fermented foods, produced through desired microbial growth and enzymatic activities (Marco et al., 2021), are the primary natural source of microorganisms in the human diet, and hold significant cultural, nutritional, and ecological values (Hernández-Velázquez et al., 2024). Fermented foods are popularly perceived as beneficial for health, possibly due to the nature of fermentation as a traditional, low-input food processing technique, and source of dietary microorganisms. However, despite their roles in shaping fermented food products, supporting regional biodiversity, influencing gut health (Valentino et al., 2024), and even contributing to planetary well-being (Cavicchioli et al., 2019; Gupta et al., 2016), the functional significance of microorganisms in food production and health remains largely overlooked by the general public.

Studying food microbiomes presents significant hurdles, particularly in obtaining samples that reflect diverse geographical, cultural, and procedural origins, a limitation compounded by the constrained reach of traditional research approaches. Citizen science (CS) offers a transformative solution to overcome these challenges while also strengthening the connection between scientists and the public. Through public involvement in microbiome research — ranging from simple contributory efforts to fully co-designed initiatives — scientists can gather more representative datasets that reflect the ecological, cultural, culinary, or health-related diversity of microbiomes. Beyond expanding the scope of research, CS fosters community participation in scientific discovery, mutual learning and creates awareness of the critical roles microorganisms play in daily life (Garcia et al., 2023). It also acts as a platform for innovation, public engagement, and evidence-based policy advocacy, ensuring the microbial world’s significance is recognized and valued across disciplines and societies (Fraisl et al., 2022).

Integrating CS into system-level microbiome studies unlocks insights essential for addressing global challenges such as antimicrobial resistance, climate change, and food security (Lewandowski et al., 2017; Pateman et al., 2020). In fermented food research, it also highlights overlooked microbial ecosystems and deepens public and academic understanding of microbial diversity, bridges scientific research with traditional knowledge, and underscores its role in sustainable food systems, cultural heritage, and human and planetary health. For example, Kefir4All and the Global Sourdough Project highlighted the roles of microbial biodiversity in shaping the production and quality of home-made fermented foods (Landis et al., 2021; Walsh et al., 2024). Through extensive public participation, diverse outreach, and transparent dissemination of findings, these projects showcase the power of CS to generate robust datasets, challenge prevailing assumptions, and strengthen connections between scientists and the public.

Sourdough, a traditional fermented mixture of water and cereal flour(s) with roots traceable to ancient times, has played a key role in the evolution of diverse food cultures by enhancing the sensory attributes, texture, nutritional value, and microbiological safety of cereal-based foods (Cappelle et al., 2013). Fermentation allows microbial growth and metabolism in the dough, producing sourdough that can serve as a leavening agent and can be propagated through backslopping—adding a portion of fermented dough to fresh flour and water (Arora et al., 2021; Calvert et al., 2021; De Vuyst et al., 2017, 2023). Despite its simple ingredients, traditional sourdoughs exhibit remarkable microbial variability, influenced by differences in household and bakery practices, flour type, dough yield, backslopping ratio, and fermentation/storage conditions (Comasio et al., 2020; De Vuyst et al., 2017; Landis et al., 2021). As most studies to date focus on laboratory-prepared or small-scale bakery sourdoughs, comprehensive large-scale investigations of spontaneously fermented (Type 1) sourdoughs in real-world settings are still needed (Bondt et al., 2024).

Here, we describe the design and execution of a large-scale sourdough fermentation study, utilizing a CS co-design approach to facilitate greater collaboration with participants. The study was conducted within the framework of the HealthFerm project (https://healthferm.eu/), a European initiative exploring innovative pulse- and cereal-based food fermentations, their potential health effects, and consumer perceptions. Through its citizen science component, HealthFerm aims to advance understanding of the sourdough ecosystem and develop novel microbial resources to support the targeted redesign of fermented foods, promoting healthier and more sustainable diets. Here, we highlight important study design considerations as well as the technical challenges and potential biases inherent to this approach and present sourdough citizen science insights. Key outcomes include broad participant engagement, the creation of geographically diverse datasets, and valuable insights for future CS initiatives. Altogether, the project explored sourdough maintenance practices and consumer preferences and perceptions across Europe, establishing a framework for microbiologists designing collaborative, volunteer-driven microbiome research.

## Results

### Collaborative, iterative co-design guides methodological citizen science implementation and refinement

Collaborative methodological planning was carried out with HealthFerm partner institutions and through direct engagement with citizens. Five universities served as local citizen science centers (hubs) to promote the study, recruit participants, and collect sourdough samples (Fig. S1a). Three hubs organized co-design workshops located in Herent (Belgium), Helsinki (Finland), and Zurich (Switzerland), involving 70 participants (27, 17, and 26, respectively). These workshops played a central role in shaping and refining methodologies throughout the project (Fig. 1, Fig. S1b).

**Figure 1.**
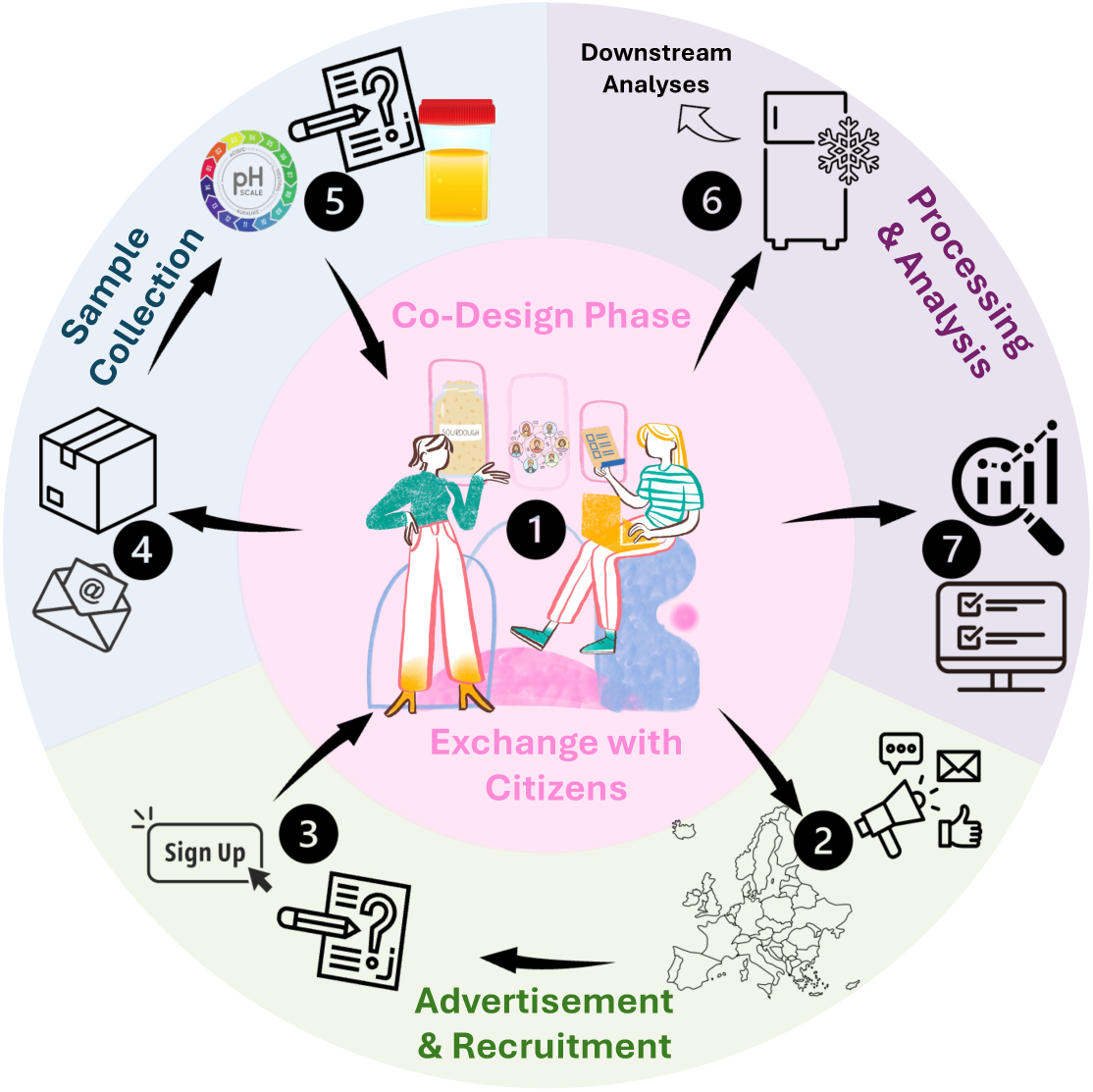
Co-design-based study setup for conducting sourdough fermentation related surveys and sampling sourdoughs across Europe. Study workflow: Co-design workshops and protocol harmonization (1) were followed by project advertising (2) and online recruitment (Survey 1) (3), local screening, and distribution of sampling kits (4). Participants conducted home experiments (sourdough/bread pH, bread density, sensory evaluation), reported results via Survey 2 (5), and mailed sourdough samples to hub labs. Samples were processed, bio banked (6), and metadata (resources 1, 2, and scientist-recorded data) curated and translated for integration and analysis (7). Icons used in this figure were sourced from Canva (www.canva.com) and edited in Microsoft PowerPoint.

Participants showed strong engagement and willingness to conduct at-home experiments and complete follow-up questionnaires. Their feedback informed improvements to the sampling kits, such as simplifying terminology, adding a project overview, and clarifying return instructions. Participant questions and interests were gathered to guide dissemination across project stages (Supplementary list 1).

Participants from different countries reported varying practices for sourdough refreshment, particularly in recipe precision. In Helsinki, only 66% used scales and exact recipes, compared to 85% in Zurich, reflecting a preference for volumetric units in Finland. Therefore, to standardize at-home measurements, a volume-to-mass conversion table was included in the instruction booklet. Feedback also guided decisions on sample submission and communication preferences. Local postal services were selected as the main delivery method, with optional drop-off for those near hub labs; sampling kits were distributed exclusively by mail. In response to requests for centralized updates, a multilingual blog was added to the HealthFerm website (Fig. S1b).

### Engagement patterns and sample contributions in a trans-European citizen science study

Extensive outreach across Europe led to strong engagement primarily from European as well as some non-European participants (Fig. 2a). By September 2024, the project had registered 1091 sourdoughs and collected 671 samples from 591 participants across 33 countries, including 27 in Europe (Fig. 2b). Most sign-ups (68.6%) and samples (65.7%) originated from hub countries (Switzerland, Finland, Belgium, Italy, Romania) and Germany (Fig. 2c), resulting in a slightly skewed language distribution (Fig. 2d). 61 individuals submitted more than one sourdough (n = 2–5).

**Figure 2.**
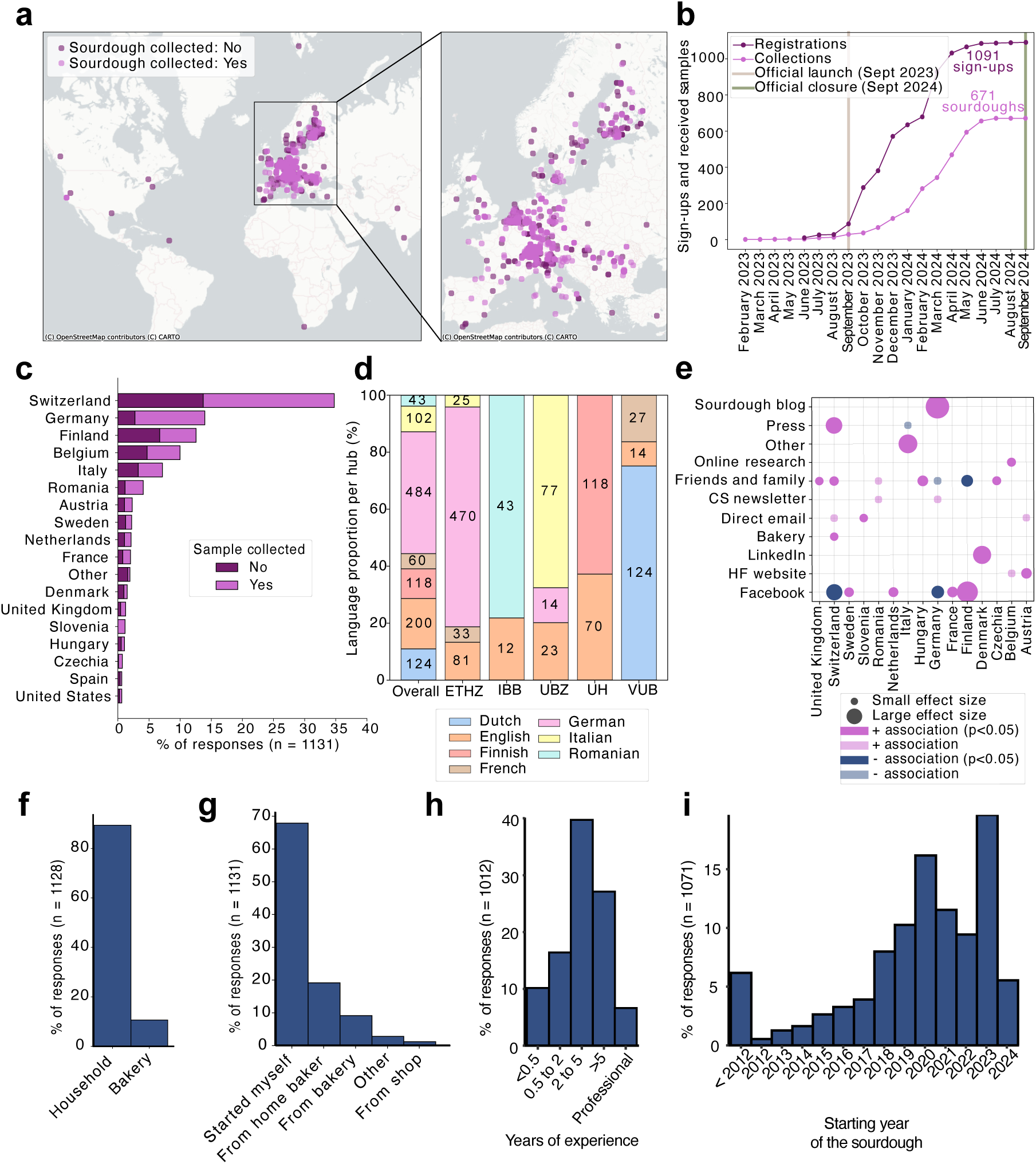
Sourdough citizen science participation and participant profiles. **a,** Global distribution of registrations and collected sourdough samples, with highest coverage in Europe. Overlapping dots indicate densely sampled regions. **b,** Registrations and sample collections over time, from project launch to campaign closure. **c,** Distribution of sign-ups and collected samples across top-contributing countries. Countries with fewer than five registered sourdoughs are grouped under “Other.” Hub countries show the highest coverage. **d,** Language distribution of all received registrations (n = 1131), including language splits per CS hub. Segment labels indicate absolute counts. **e,** Most effective advertising channels by country. Associations assessed via Chi-squared tests; only significant associations shown. **f,** Distribution of sourdough sources. “Bakery” includes all non-household origins (e.g., artisanal bakeries, industry, restaurants). Samples with unknown origins excluded (n = 3). **g,** Reported origins of registered sourdoughs. **h,** Participant experience with sourdough, ranging from less than half to more than 5 years, including professional bakers. **i,** Years when sourdoughs started.

Recruitment success varied by country and channel (Fig. 2e, Fig. S2a): sourdough blogs were particularly effective in Germany, press in Switzerland, LinkedIn in Denmark, and Facebook in Finland (Table S2). In contrast, word-of-mouth in Finland and Facebook in Switzerland and Germany were less impactful.

Merging sign-up forms and survey responses showed that 87.2% of sourdoughs came from household bakers (Fig. 2f), with 67.7% self-initiated and 8.0% passed on by others (Fig. 2g). Regarding experience, 26.6% had baked for under two years, while over a third had five or more years of experience or were professionals (Fig. 2h). Most sourdoughs were less than a year old at sampling (Fig. 2i), with a notable peak of newly started sourdoughs in 2020, reflecting the surge of popularity in sourdough baking during the COVID-19 pandemic; the oldest registered sourdough dated back to 1865.

### Fermentation routines and storage conditions differ between households and bakeries

Differences in environmental and fermentation practices were observed between professional (n=109) and home bakers (n=1019) (Fig. 3). Sourdoughs from both groups were proportionally distributed across European countries, minimizing regional bias (Fig. S2b). Professional bakers more often maintained older starters (55% > 5 years vs. 28% in households), refreshed them daily (41.3% vs. 43.7% weekly in households), and fermented at higher temperatures (23 °C vs. 21 °C) with shorter fermentation times. Both groups typically stored sourdoughs at or below 10 °C when not in use. Plastic containers were more common in bakeries (61.5%), while home bakers preferred glass jars (only 9.4% use plastic containers). The presence of plants, pets, or children in the baking environment did not differ significantly, likely due to unclear survey wording. Ingredients were similar across groups (Fig. 3c): wheat was most common (52.3% bakeries; 53.4% households), followed by rye, with widespread use of organic flours (66.1% bakeries; 59.6% households). Most reported dough yields 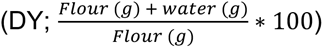 between 150 and 250, indicating firm to semi-liquid sourdoughs.

**Figure 3.**
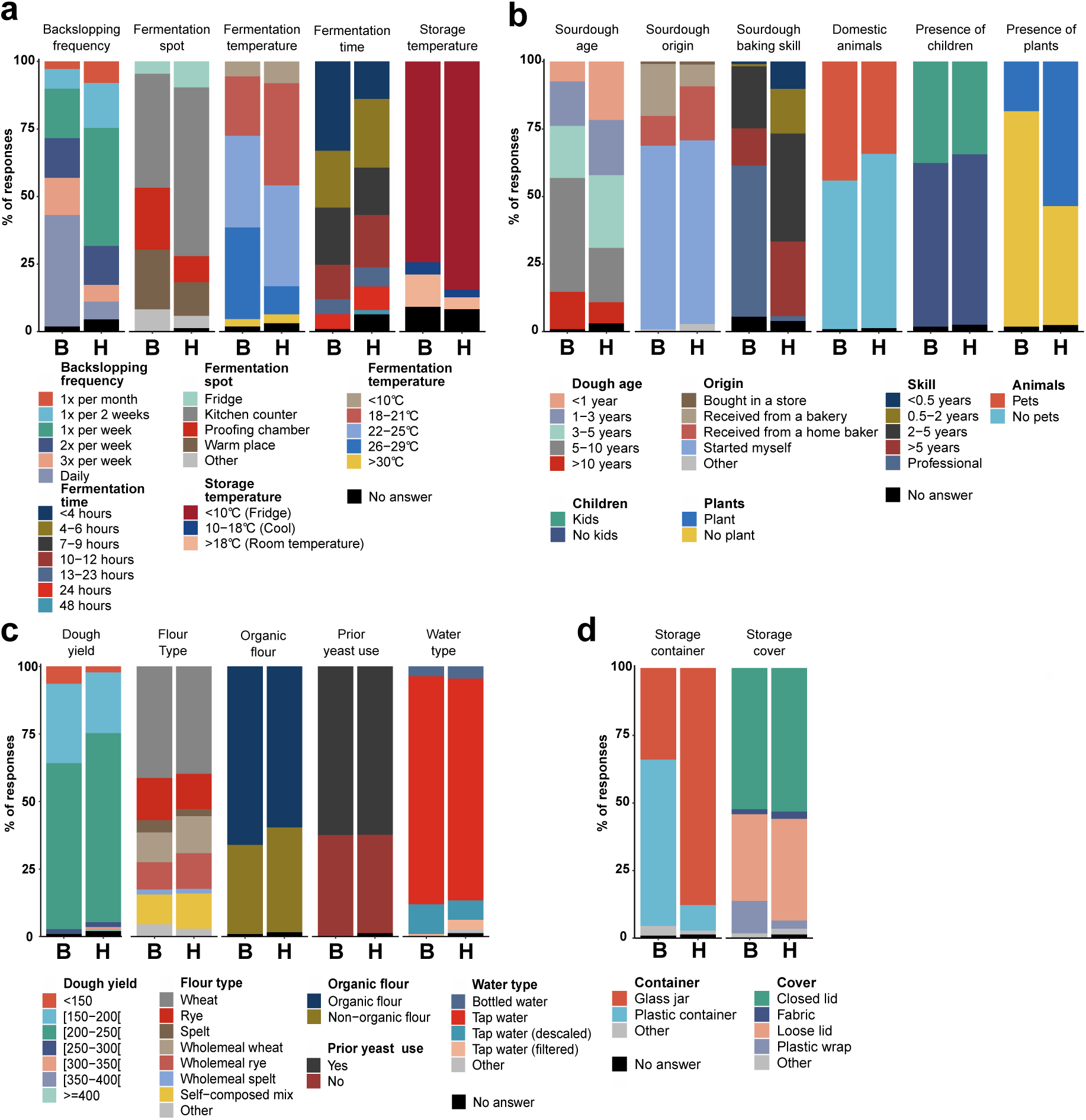
Variations in the registered process and environmental parameters, ingredients and fermentation vessels used in bakeries (B; 109 responses) and households (H; 1019 responses). **a,** Distribution of fermentation related parameters: the backslopping frequency, the location of the sourdough during fermentation, the temperature and time during fermentation and the temperature of storage of the sourdough between feeding moments. **b,** Distribution of various environmental parameters: sourdough age (categories defined with the lower limit included and the upper limit excluded), origin of the sourdough, the skill level of the sourdough baker, the ownership of pets and the presence of kids and plants. **c,** Distribution of sourdough ingredient related parameters: Dough yield (categories defined with the lower limit included and the upper limit excluded), flour type (*Triticum aestivum* is referred to as “wheat,” *Triticum spelta* as “spelt,” and *Secale cereale* as “rye”), the use of organic flour, the prior use of baker’s yeast in the environment where the sourdough is maintained, and the water type used when feeding the sourdough. **d,** Characteristics related to the vessel in which the sourdough is maintained.

### Baking motivations and perceived health benefits shape ingredient choices

Personal motivations and beliefs about sourdough varied between participants. Participants most frequently cited hedonistic motivations such as taste and a general interest in baking, and financial/tradition motivations less frequently (Fig. 4a). A majority (84.2%) perceived sourdough as healthier than yeast-leavened bread, with no significant difference across experience levels (Fig. S3a, *p* = 0.115). Gastrointestinal benefits were the most reported perceived health benefit, again consistent across levels of baking experience (Fig. 4b, Fig. S3b, *p* = 0.115). Health-motivated bakers significantly more frequently used rye and less frequently wheat as the grain base (Fig. 4c, Table S3). Tradition-driven bakers more often used rye and spelt, while rye was also overrepresented among those citing health-related reasons such as using “healthier ingredients” or experiencing “fewer intolerances” compared to baking with conventional yeast (Fig. 4c, Table S3). Bakers using wholemeal or flour mixes more often reported enjoyment, health motivation, and gastrointestinal benefits, compared to those using endosperm or unspecified flour types, who were less likely to cite enjoyment, taste, or digestibility (Fig. 4c, Table S3). Health-motivated bakers also used organic flour significantly more often and associated it with improved nutritional quality or bioavailability (Fig. 4c, Table S3).

**Figure 4.**
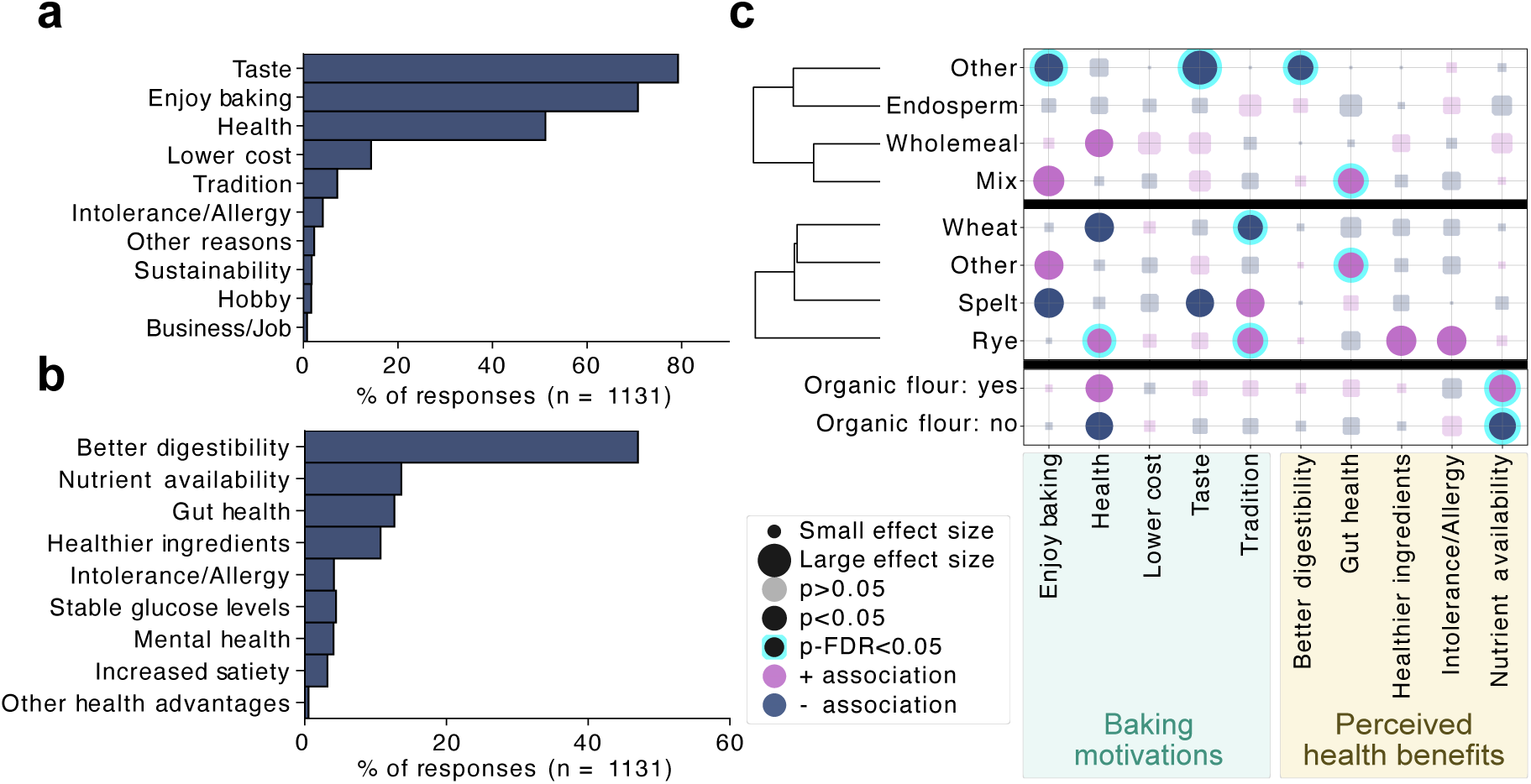
Personal motivations and perceptions related to sourdough baking. **a,** Participant- reported motivations for baking with sourdough, assessed using a “check-all-that-apply” format. **b,** Perceived health benefits of sourdough bread baking and consumption. Free-text responses were thematically categorized into nine main benefit categories. **c,** Chi-square analyses of associations between selected baking motivations and perceived health benefits and the categories flour milling grade, grain type, and use of organic versus non-organic flour. Positive and negative associations are color-coded; bubble size reflects effect size (Cramér’s V), and statistically significant associations are highlighted. Associations are hierarchically clustered per category (flour milling grade and grain type) based on directional Cramér’s V.

### Citizen science experiments reveal patterns and challenges in sourdough physicochemical and sensory characteristics

Among participants who submitted sourdough samples, 63.6% (427/671) completed five optional home experiments, while 15.6% did not perform any. In 13.8% of cases, no sourdough bread was baked, and 8.6% of those who baked did not complete the experiments. pH measurements performed at home revealed a positive correlation between sourdough and bread pH (Fig. 5a), indicating that sourdough acidification generally yielded more acidic bread, regardless of flour type. Bread density, however, was not affected by pH (Fig. 5a, Fig. S4). Laboratory-measured pH values were on average 0.73 units lower than home measurements (Fig. 5b, Fig. S5), likely due to further acidification during shipping and/or the use of accurate laboratory pH meters versus pH strips at home.

**Figure 5.**
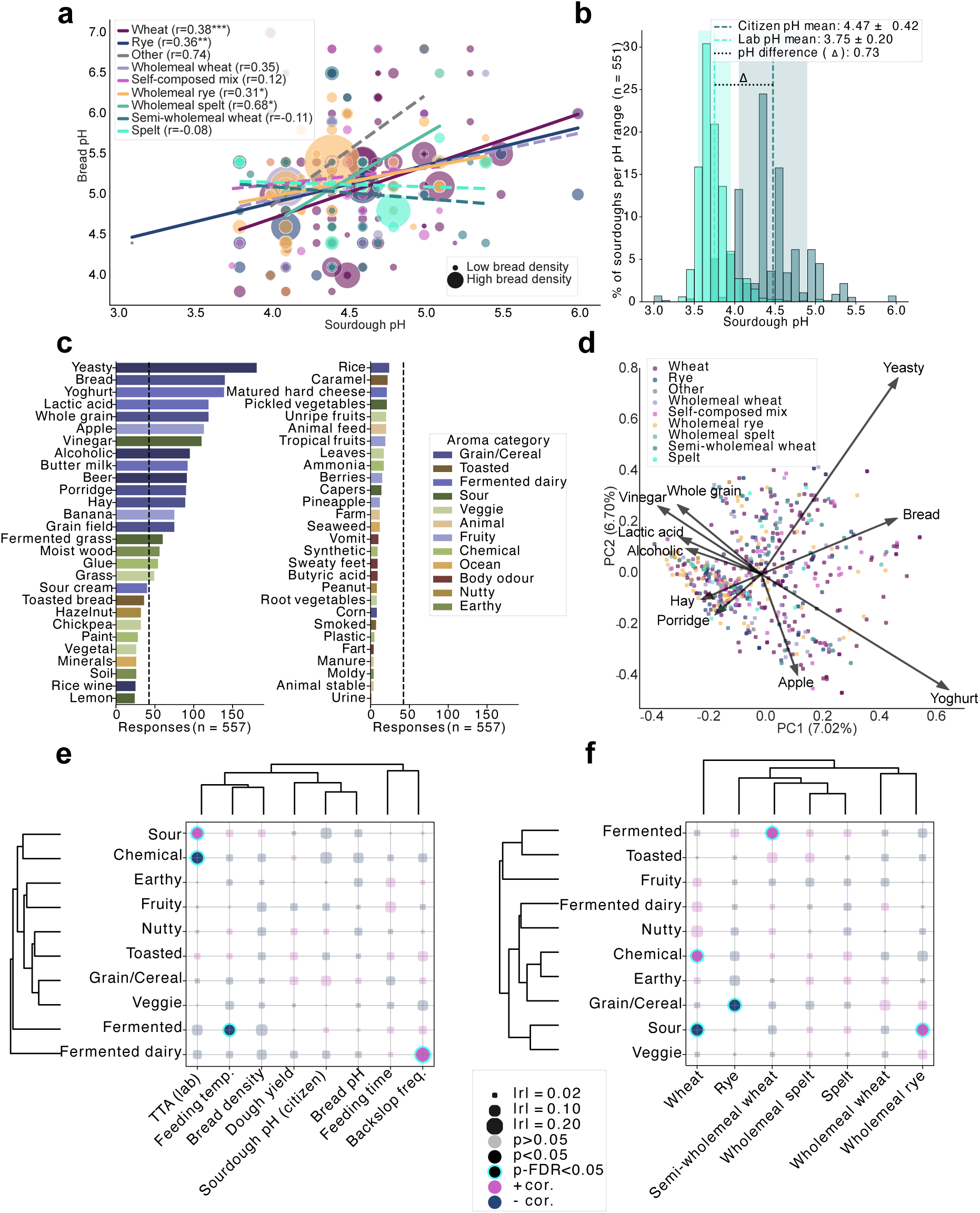
Citizen science contributions to sourdough characterization and sensory profiling. **a,** Correlation between sourdough and bread pH values, grouped by flour type used for feeding the sourdoughs; bubble size reflects bread density; significant Pearson correlations indicated (*p<0.05, **p<0.01, ***p<0.001). **b,** Comparison of pH measured by citizens vs. laboratory measurements post-shipping under non-temperature-controlled conditions. **c,** Reported sourdough odors ranked by frequency and colored by flavor category (n = 557); average occurrence indicates mean frequency of sensory notes. **d,** Principal Coordinates Analysis (PCoA) of Jaccard dissimilarity matrix for odors; top 10 aroma features contributing most to variance shown, colored by sourdough flour type. **(e, f)** Pearson correlations of aroma categories with feeding parameters and experimental measurements **(e),** and sourdough flour types **(f)**; features and predictors hierarchically clustered by correlation direction.

Participants characterized the aroma of their sourdoughs using the Check-All-That-Apply (CATA) methodology and selected from 56 predefined descriptors, organized into 13 categories (Fig. 5c). Fermented, grain/cereal, fermented dairy, fruity, and sour aromas were most prevalent; ocean, animal, and body odor were least common. High variability in aroma perception was evident in Jaccard dissimilarity-based PCoA (Fig. 5d), where the first two components explained only 13.7% of the total variance. Features contributing most to the variance included yeasty, bread, and yoghurt all positively correlated with PC1, whereas porridge, hay, lactic acid, alcoholic, vinegar and wholegrain aromas also ranked in the top ten features but negatively correlated with PC1 (Fig. 5d). PERMANOVA identified categorical variables such as plant presence, organic flour use, milling grade, and acidic beverage fermentation as significantly associated with aroma variation (BH-FDR p < 0.05), although effect sizes were low (R² < 0.005; Table S4). Among numerical variables, wheat proportion and the number of maintained sourdoughs (positive correlation) and backslopping frequency (negative correlation), were significant but weak predictors (Mantel Pearson R < 0.1; Table S5). Notably, not clear separation was observed when grouping samples by flour type in the full aroma feature space (Fig. 5d, Table S5).

Nonetheless, specific aroma descriptors in isolation correlated with experimental data and flour types. Sourdoughs described as “sour” had significantly higher TTAs (Fig. 5f, Table S6) and were more frequently composed of wholemeal rye flour (Fig. 5g). Wheat sourdoughs demonstrated distinct aroma characteristics (Fig. 5g), were less sour and more often associated with chemical odors, which negatively correlated with TTA (Fig. 5f, Table S6).

### Substrate-dependent variation in sourdough composition, maintenance, and physicochemical properties

The cereal used for sourdough influences its physicochemical properties and associated recipes. Wheat and rye, the most commonly used cereals for sourdough feeding (Fig. 3c), were selected for in-depth analysis. Reflecting rye’s higher water absorption, participants using rye maintained sourdoughs at higher dough yield than those using wheat (r = 0.30, p = 3.10×10⁻¹⁸; Fig. 6a). Rye sourdoughs also had slightly lower participant-measured pH (r = 0.12, p = 8.84×10⁻⁴) and higher TTA values (r = 0.37, p = 3.0×10⁻²⁶), despite similar lab pH (p = 0.15) (Fig. 6b,c). Participants baking with rye reported denser breads than those using wheat (r = 0.37, p = 3.48×10⁻¹¹; Fig. 6d). These differences were mirrored in sourdough clustering based on Pearson correlations between proportion of flour type used per sourdough and physicochemical or feeding parameters (Fig. 6e). Sourdoughs with high rye content (wholemeal or not) formed a distinct cluster, associated with higher TTA and dough yield, and lower pH and backslopping frequency. Sourdoughs with non- rye wholemeal flours or high spelt content showed intermediate profiles between rye- and wheat- based sourdoughs (Fig. 6e, Table S7).

**Figure 6.**
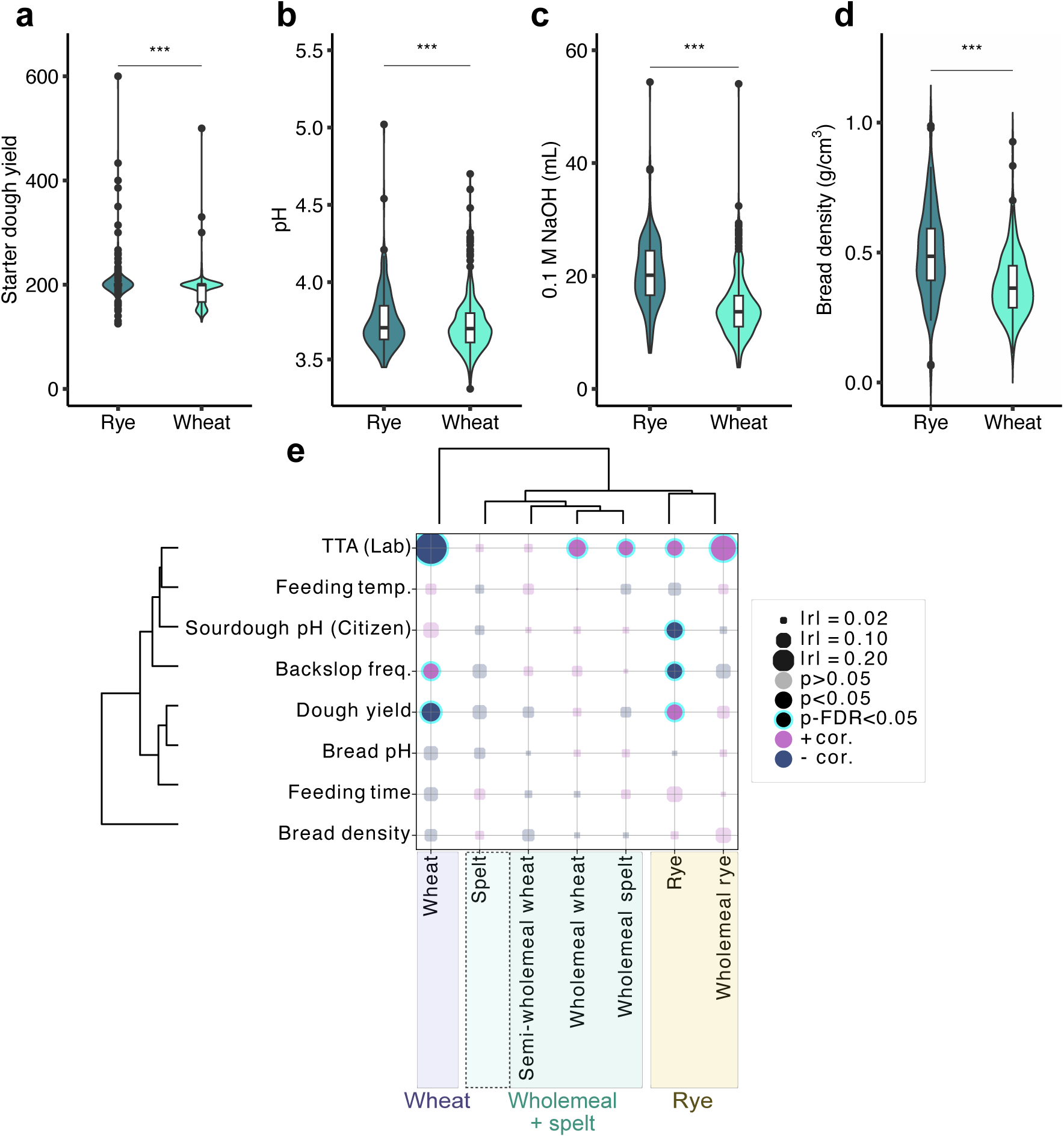
Substrate-dependent variation in sourdough physicochemical properties and maintenance practices. **a,** Comparison of dough yield between wheat- and rye-based sourdoughs. **b,** Participant-reported pH values using pH strips. **c,** Total titratable acidity (TTA) measured in the laboratory, defined as the volume (mL) of 0.1 M NaOH required to reach pH 8.5 in 10 g of sourdough diluted with 100 g deionized water. **d,** Bread density of breads made with wheat or rye sourdough, as reported by participants. Values >1 were considered erroneous and excluded from analysis. **(a,b,c,d)** Statistical comparisons were performed using the Mann– Whitney U test with BH-FDR correction (p ≥ 0.05 = ns; * < 0.05; ** < 0.01; *** < 0.001). **e,** Hierarchical clustering of sourdoughs by substrate, based on Pearson correlations between flour compositions (including mixed flours), feeding parameters, and sourdough characteristics. Bubble size reflects absolute correlation strength (|r|), and color indicates direction (positive/negative).

### Fermentation fingerprints: how region, substrate, and role define sourdough fermentation practices

Sourdough ingredient choices varied regionally across Europe, shaped by cultural traditions, climate, and lifestyle. Chi-square enrichment analysis revealed significant country-specific differences in flour types, milling grades, water sources, added ingredients, flour switching, organic use, and concurrent yeast usage (Fig. 7a, Table S8). Clustering countries by directional Cramér’s V effect sizes highlighted a distinct group of German-speaking countries (Germany, Austria, Switzerland), characterized by a preference for organic wholemeal flours — especially rye and spelt — whereas Romanian, Finnish, and Italian participants more often preferred non- organic, wheat-based sourdoughs. Northwestern European bakers (Belgium, Denmark, Sweden, UK) commonly used mixed flours and grain blends. Water type and commercial yeast practices also varied by country (Fig. 7a).

**Figure 7.**
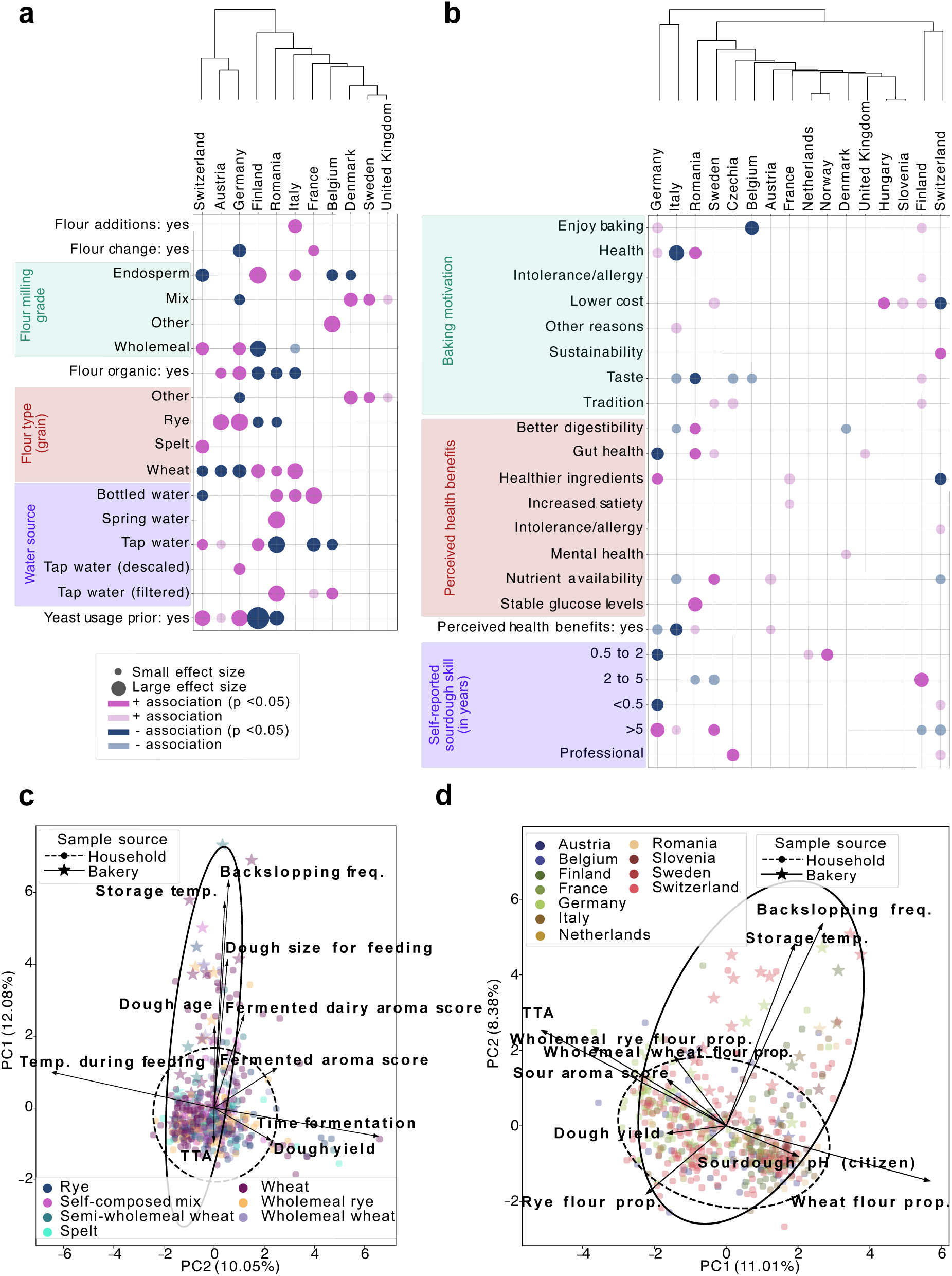
Regional variation in sourdough ingredients, maintenance practices, and physicochemical characteristics among European citizen scientists. **a,** Bubble plot of significant Chi-square associations showing regional differences in ingredients and maintenance practices (e.g., non-flour additions, flour type changes, milling grade, organic use, grain base, water source, baker’s yeast). **b,** Heatmap of baking motivations and perceived health benefits by country and self-reported skill level. **(a,b)** Only categories with at least one significant association are shown. Associations are color-coded by enrichment direction and significance (BH-FDR p < 0.05: dark; uncorrected p < 0.05: light); bubble size reflects effect size (Cramér’s V). Countries are clustered by Cramér’s V profiles. **c,** PCA of numeric metadata: feeding parameters, home experiment results, TTA, and latitude (excluding substrate). **d,** PCA including feeding parameters, home results, lab TTA, and flour proportions (excluding geography). **(c,d)** Samples with missing values were excluded. Points are colored by flour type or country; ellipses show 95% confidence intervals for household vs. bakery sources. Groups with <5 samples were excluded.

Regional differences also emerged in baking motivations and perceived health benefits (Fig. 7b, Table S9), sourdough storage practices, and household environments (Fig. S6, Table S10). Swiss participants emphasized sustainability, Romanians cited health benefits (digestibility, gut health, glucose control), and Italians rarely mentioned health as a motivation. Enjoyment and taste dominate across countries but are less reported in Belgium. Hierarchical clustering shows shared motivational patterns and sourdough baking experience levels between Swiss and Finnish bakers, while Germany and Italy cluster by a higher proportion of experienced bakers and fewer novices (Fig. 7b). In addition to sourdough, other food and beverage fermentations are commonly practiced in France, Austria, and Germany (especially dairy), but rare in Finland (Fig. S6).

Despite patterns related to country of origin and flour preferences, PCA of fermentation features did not yield clustering by country or flour type (Fig. 7c,d). Instead, PCA consistently separated household and bakery sourdoughs, pointing in opposite directions in both geographic and flour- related feature spaces (Fig. 7c,d). This household-bakery split was mainly driven by higher backslopping frequency, storage temperature, dough size, age, and fermented dairy aromas in bakery sourdoughs (Fig. 7c,d). As expected, feeding temperature was associated with shorter fermentation time and dough yield (Fig. 7c). Together, these results underline both country- and flour-related trends in how sourdoughs are fermented, while emphasizing individual variation and a strong divide between household and bakery sourdoughs.

## Discussion

The diversity of processing practices and parameters used in the production of traditional food fermentations plays a significant, yet often understudied, role in their microbiological and chemical properties, environmental sustainability, cultural heritage, and human health effects. Studying the diversity of home fermentation practices at the continental scale presents a non-trivial challenge for conventional scientific research that typically lacks the means to gather such comprehensive data. Citizen Science (CS) offers a solution to this challenge by enabling large-scale sample collection while simultaneously fostering public engagement and enthusiasm for science, strengthening the connection between scientists and the public, and raising awareness about the importance of scientific research related to food fermentation.

The present study demonstrates the potential of a co-design-driven CS approach to investigate the diversity of technological and cultural parameters of sourdough fermentation across Europe. Through the participation of over 1000 citizen scientists and the collection of more than 670 sourdough samples, a large and geographically diverse dataset was established that would have been difficult to obtain solely through conventional research methods. The co-design-based approach facilitated broad public participation and mutual learning between citizens and researchers, resulting in unique insights into sourdough maintenance motivations and practices, ingredient choices, and perceived health benefits. This approach enabled a comprehensive assessment of sourdough fermentation practices across Europe and permitted validation or reconsideration of prevailing assumptions surrounding traditional sourdough fermentation practices. Furthermore, the project established a sourdough biobank that will serve as a valuable resource for future fermented food research. A key distinction between this study and previous initiatives — including CS studies like the Global Sourdough Project and experimental studies like Reese *et al*. (Landis et al., 2021; Reese et al., 2020) — is the participatory nature of the HealthFerm citizen science model. Here, participants actively contributed to experiment selection and protocol refinement through feedback loops, making the process more adaptive and participant-centered. While earlier initiatives compiled metadata, the dataset presented in this study includes novel and uniquely detailed information - not only on sourdough fermentation itself, but also on behavioral and cultural dimensions, incorporating comprehensive sensory evaluations and perceived health benefits. Moreover, this co-designed citizen science framework is scalable and transferable to other traditional food systems. It can be tailored to address specific challenges in sample collection or metadata design, underscoring the value of iterative participant engagement in shaping robust research protocols.

Pre-launch workshops enhanced engagement by helping to refine sampling kits, instructions, and questionnaires, improving clarity and accessibility. Direct feedback and interactions between citizens and researchers revealed strong participant motivation to contribute to the study, carry out home experiments, and stay informed about project progress and outcomes. These exchanges suggested that hands-on involvement fostered a sense of being valued, sparked curiosity, and strengthened commitment. The collaborative, iterative process appeared to build trust and a sense of ownership, likely sustaining participant interest — especially in the study outcomes. Future citizen science projects may benefit from emphasizing co-design and transparent researcher–participant communication to maintain engagement and foster long-term scientific investment.

A significant finding of this study is understanding the public perceptions of the health benefits of sourdough bread. Most participants considered sourdough to be healthier than yeast-leavened bread, commonly citing improved digestibility and gut health as motivating factors for maintaining and baking with sourdough. Public perceptions of health benefits align well with recent scientific findings demonstrating increased mineral bioavailability in sourdough-based products *in vitro,* and evidence for improved digestibility or lowering the glycemic index *in vivo* (Arora et al., 2021; D’Amico et al., 2023; Ribet et al., 2023; Rizzello et al., 2019). Nevertheless, these perceptions reflect the intersection of traditional food fermentation, consumer beliefs, and scientific inquiry, emphasizing the relevance of sourdough in public discussions on health and nutrition.

The study also faced several challenges. Recruitment biases — particularly self-selection bias, where volunteers were likely more motivated or interested in fermentation than the general public — may have skewed participation. Such imbalances are inherent to citizen science, despite broad outreach, and were compounded by logistical challenges, variability in citizen-conducted experiments, and unpredictable sample arrivals, complicating scheduling of microbiological analyses (Berghen et al., 2024; Constant & Hughes, 2023; Waugh et al., 2023).

Geographical coverage also presented challenges. Advertisement strategies across a multilingual continent needed persistence and a multifaceted approach, working best when conducted in local languages. Project recruitment and materials were made available in seven languages, reflecting the national languages of the hub labs involved in this project as well as English. Nevertheless, regional language barriers still posed significant obstacles throughout the recruitment, data collection, and communication phases. This most likely explains the disproportionate geographic representation in the project. Countries sharing a language with the research hubs exhibited a high degree of participation (e.g., Switzerland, Germany, and Austria all share German as one of their official languages) whereas countries without language representation in the project yielded low or no participation (e.g., Spanish, Greek, Polish, *etc.*). Future citizen science initiatives may benefit from a strong, multilingual social media presence — potentially supported by digital marketing professionals and enhanced with interactive tools such as chatbots — to increase visibility, foster engagement, and encourage continued participant involvement.

Logistical barriers also impacted the study. Sampling kit and sourdough sample shipping was most efficient within the European Union (EU), but international shipments crossing non-EU borders encountered issues related to customs regulations and inconsistencies between postal services (availability of prepaid return labels and other services). Additionally, the absence of temperature-controlled shipping resulted in continued fermentation during transit, and prolonged transit times often led to gas buildup, causing some sourdoughs to leak from their containers. To mitigate this issue in future studies, it may be advisable to instruct participants explicitly to ship samples early in the week, thereby reducing the risk of extended exposure to non-cooled conditions during prolonged transit times (e.g., days when postal delivery services are not available). This continued fermentation possibly explains discrepancies between pH measurements taken at home and those recorded upon arrival at the laboratory. Alternatively, this discrepancy could have arisen from the use of colorimetric pH strips with low dynamic range (blue and green for pH 3 to 6) and less precision than the calibrated electronic pH meters used in the hub labs. Follow-up studies are necessary to evaluate potential microbiological and chemical changes that could have arisen from prolonged fermentation without temperature control. Addressing this uncertainty will help refine protocols for sample collection and shipping, ensuring higher data quality in future CS projects where samples are shipped through the postal system.

Despite participant-obtained data often being of similar quality than scientist-obtained data (Lewandowski et al., 2017), the large variability in participant-reported data posed significant challenges for curation and interpretation, particularly regarding home experiment results. This variability aligns with findings from citizen science projects, where differences in volunteers’ skills, motivations, and task familiarity can lead to inconsistencies in data collection (Weigelhofer & Pölz, 2016; Wiggins & Wilbanks, 2019). Sensory evaluations were conducted using the CATA methodology, a reproducible and consumer-appropriate approach to characterize complex aromas. Here, CATA enabled the collection of large volumes and diverse data. However, the observed variability was high - stemming from both sourdough heterogeneity and individual perceptual differences - which underscores the need for cautious interpretation of citizen- contributed sensory data. Extending workshop formats beyond co-design to include practical sessions prior to performing home experiments could enhance data accuracy and reliability in future CS studies (Fleming et al., 2015; Piochi et al., 2021).

Beyond methodological challenges and lessons learned for future CS initiatives, the study revealed valuable insights into sourdough maintenance practices across Europe. Distinct differences emerged between household and bakery sourdoughs, with bakeries typically maintaining older sourdoughs, backslopping more frequently, and fermenting at higher temperatures to meet the fast-paced demands of professional baking. In addition, the choice of cereal also emerged as a key factor shaping sourdough properties. Rye sourdoughs typically exhibited higher dough yields and total titratable acidity compared to wheat-based sourdoughs. These differences can likely be attributed to the higher arabinoxylan content and enzymatic activity of rye flour, which influences water absorption and fermentation kinetics (Courtin & Delcour, 2002; Deleu et al., 2020). Interestingly, the widespread use of organic flour among many participants reflects a strong consumer preference, suggesting that ingredient choices may influence microbial composition in ways that warrant further investigation in follow-up studies.

One of the most intriguing findings of the study was the emergence of strong regional signatures, where countries exhibited distinct ingredient preferences, fermentation habits, and environmental conditions for maintaining sourdough, as well as differences in baking motivations and perceived health benefits. For instance, German-speaking countries (Germany, Austria, Switzerland) showed a clear inclination toward organic wholemeal flours, particularly rye and spelt, reflecting a robust tradition of non-wheat-based bread making. In contrast, participants from southern and northern Europe primarily used wheat-based sourdoughs and non-organic flours, despite their rich sourdough bread cultures. Interestingly, Finnish participants preferred wheat sourdoughs, despite rye bread’s national prominence. This may suggest that rye breads are more commonly purchased than homemade, or it might reflect a latent recruitment bias, where traditional bakers might have been underrepresented or less likely to volunteer for the study. Instead, participants were more likely to be motivated by enjoyment, taste and perceived health benefits rather than tradition, as supported by the fact that fewer than 10% reported baking for traditional reasons, while over 80% cited taste and 70% the enjoyment of baking (Fig. 4a).

This study showcases the potential of co-designed citizen science to enable large-scale, multicultural research and generate comprehensive, population-level datasets that can be integrated with microbiome analyses. It also highlights the need for improved strategies to mitigate demographic biases, ensure data reliability, and address logistical challenges, particularly in multilingual and multinational contexts. The widespread belief in sourdough-related health benefits underscores the importance of continued scientific validation, especially regarding gut microbiota and metabolic health (Wiggins & Wilbanks, 2019). Community-generated insights from this study lay a foundation for future investigations into links between sourdough consumption and health outcomes. Moreover, the framework established here offers a scalable model for participatory research, applicable not only to sourdough but also to other traditionally fermented foods. By leveraging citizen participation, future efforts can bridge scientific inquiry with traditional knowledge, advancing our understanding of fermented food microbiomes and their implications for human and planetary health.

## Materials and methods

### Implementation and coordination of a multi-hub citizen science study

A co-design-driven CS project embedded in the European HealthFerm project lies at the heart of the current study. From the design phase onward, several key considerations were identified and addressed. These include compliance with key privacy and ethical regulations (such as the General Data Protection Regulation (GDPR), the Nagoya Protocol on Access and Benefit Sharing, and ethical approval), the need for communication in the native languages of the citizen scientists throughout the recruitment, experimental, and dissemination phases, logistical challenges such as sample shipping (*e.g.*, shipping time, administration, and costs), and unpredictable sample collection timelines influencing the following laboratory processing workload.

To manage these complexities, the CS effort was subdivided across five sampling hubs across Europe, each covering a specific geographical area (Fig.1). The different hubs were each responsible for the GDPR and Nagoya protocol compliance within their collection areas. This subdivision also simplified translation efforts, communication with the participants in seven languages, and the shipping arrangements with local couriers. However, it also introduced challenges in ensuring result comparability, requiring the standardization of protocols and documentation across all hubs. The sampling hubs were located at the Free University of Bozen-Bolzano (UBZ, Italy), Vrije Universiteit Brussel (VUB, Belgium), Institute of Biology Bucharest (IBB, Romania), University of Helsinki (UH, Finland), and ETH Zurich (ETHZ, Switzerland).

### Co-design workshops and questionnaire design

We chose to follow a co-design process to involve representative participants in the design of research objectives and procedures (Fig. 1, Fig S1b). These were further refined through multidisciplinary collaboration within the HealthFerm consortium, drawing on expertise in sourdough research, citizen science, microbiome science, food technology, food biochemistry, consumer science, and sensory analysis. To support the co-design process, we prepared draft materials essential for the campaign, including a sourdough metadata questionnaire, an instruction booklet for sampling and at-home experiments, and a sampling kit. Feedback on these prototypes was collected through three co-design workshops held in Belgium, Finland, and Switzerland, involving small groups of citizen scientists representative of the target audience - namely home and small-scale commercial sourdough bakers. All workshops followed a similar structure, beginning with an introductory presentation on the HealthFerm project and the CS initiative, followed by an interactive, layman-friendly presentation on sourdough and the current state of sourdough research, designed to both assess participants’ knowledge levels and to provide them with a foundational understanding of the science behind sourdough. This was followed by an interactive session where participants were guided through the draft questionnaire and instructions, while also trying some proposals for at-home experiments, including a pH measurement and a sensory analysis of a provided sourdough starter. Feedback from the workshop participants was collected throughout the session. The responses obtained were compared across the different workshops and key insights were incorporated in the further development of the sampling kits and sourdough metadata questionnaire used during the subsequent sampling campaign.

### Participant registration and at-home experimentation

The citizen science (CS) sourdough campaign was launched via an online sign-up form (Fig. 1, Fig. S1b), accessed through a multilingual registration page on the HealthFerm website (available in English and the six official languages of the participating hub countries). Participants were automatically assigned to a local hub laboratory based on location. Each hub hosted its own language-specific, GDPR-compliant registration survey - including informed consent, pseudonymization, and a sample rights transfer agreement - using Microsoft Forms or Google Forms, depending on institutional preferences.

The survey included 90 questions, collecting only essential personal information (e.g., name, email, address) for kit delivery and metadata on the participant’s sourdough. Respondents could also indicate willingness to contribute sourdough or other fermented food samples.

Recruitment was promoted via social media, press coverage, print and digital ads, university networks, public lectures, and conferences. Outreach also targeted sourdough communities, fermentation networks, professional bakers, and influencers. A dedicated HealthFerm CS Blog (https://healthferm.eu/news-and-events/blog) and FAQ (https://healthferm.eu/healthferm-community/faq) were maintained on the project website, with regular updates via LinkedIn and newsletters.

To ensure diverse geographic and substrate representation, participants were sub-selected for sample submission. Selected participants received a sampling kit by mail containing instructions, materials for sourdough sampling and home experiments (Table S1), and a unique five-character code for pseudonymization. A QR code linked to a follow-up survey (using the same secure tools) to upload experiment results and updated metadata.

Participants followed a standardized protocol (archived on Zenodo; see Data availability) to analyze their own sourdoughs and breads. Sourdough pH was measured by mixing 10 g of dough with 10 g of water and using pH strips (MilliPoreSigma, Darmstadt, Germany). Bread pH was assessed by homogenizing 10 g of crumb (crust removed) in 100 g of water. Bread density was determined by measuring a ∼3 cm crustless slice’s weight and dimensions. Aroma descriptions of sourdough and bread were recorded using sensory wheels developed by the University of Copenhagen for the Wild Sourdough and Smag for Livet projects (*Lav Din Egen Surdej – Mål Og Vurdér | Smagforlivet.Dk*, n.d.; Nichols, n.d.)

### Sourdough sample collection and processing

Sourdough samples were sent in ambient conditions by mail for lab analyses. Upon reception, samples were processed within 24 hours at the local collection hubs using common protocols and chemicals. The pH was determined by diluting 10 g of sourdough in 100 mL of deionized water. The same suspension was used to assess the total titratable acidity (TTA) by titrating with 0.1 M NaOH until a pH of 8.5 was reached, with the volume of NaOH added representing the TTA value. For future analyses and biobanking, 5 g of sample was diluted in either 5 g of sterile deionized water or 5 g of sterile 50% glycerol, aliquoted into cryovials, and stored at -80 °C.

### Survey processing, statistics, and data visualization

Each hub maintained three multilingual datasets: registration surveys, result surveys, and laboratory measurements. After sample collection, pseudo-anonymized datasets were centrally compiled, translated into English (semi-automated, using Google Sheets), and merged into unified registration, result, and laboratory tables. Manual curation was used to standardize vocabulary and harmonize free-text entries. Datasets were linked via unique 5-character sample codes and person IDs (archived on Zenodo; see Data availability); missing result variables were filled from registration data when available.

Some metadata - such as sensory attributes, pH, and bread density - were only collected during the sample phase and are thus available for a subset of samples. Flour types were classified from participant-reported grain and milling grade. In this study, *Triticum aestivum* is referred to as “wheat,” *Triticum spelta* as “spelt,” and *Secale cereale* as “rye“. For chi-square analyses, semi- wholemeal and wholemeal flours were grouped (as semi-wholemeal was only available for wheat and clustered closely with wholemeal wheat in both lab and feeding-parameter data). For selected enrichment analyses, composite flour types (e.g., wholemeal wheat) were decomposed into grain base and milling grade to distinguish associations with participant region, motivation, or health perception. In contrast, numerical and feature-space analyses retained these as separate categories to preserve resolution. Descriptive and correlative analyses excluded missing data on a per-variable basis unless stated otherwise. Data integration, curation, and analysis were conducted using Python 3.12.7 and R 4.4.1 (R Core Team, 2021).

#### Descriptive analyses

Descriptive analysis based on a bakery-household distinction was performed using the dplyr, tidyr, and ggplot2 packages in R. Data were filtered to exclude the samples with an unknown sample source, along with one single sample of industrial origin. Categorization was introduced for several of the used metadata (dough age, fermentation temperature, fermentation time, storage temperature, and dough yield) by which values present at ≤ 1% prevalence in each category were binned as ‘Other’.

#### Pairwise analysis

Both t-tests and the Mann-Whitney U tests were used depending on the data’s normality and homoscedasticity, while chi-square tests were applied for binary yes/no responses and categorical variables. Effect sizes were calculated using the formula for Cohen’s d, rank serial correlation, and Cramér’s V for t-tests, Mann-Whitney U test, and chi-square tests, respectively.

#### Maps

To visualize the geographic distribution of sourdough samples, spatial data processing and mapping were conducted using Python (v3.10.14), incorporating geopandas (v1.0.1) for geospatial data handling, shapely (v2.0.6) for geometric operations, matplotlib (v3.8.4) for data visualization, and contextility (v1.6.2) for basemap integration.

#### Collinearity analysis

Collinearity among variables was assessed prior to multivariate and enrichment analysis. Categorical associations were tested using Chi-square and Cramér’s V; numerical variables using Pearson, Spearman, and cosine similarity; and mixed types with Kruskal–Wallis tests. False discovery rate was controlled using the Benjamini–Hochberg method. Hierarchical clustering of association matrices guided the selection of representative variables to avoid redundancy and inclusion of collinear variables.

#### Enrichment analyses by category

Due to higher sampling density and geographic representation in Europe, geolocation-based enrichment analyses excluded non-European samples (longitude −50 to 70, latitude 0 to 70). To assess categorical enrichment across geographic clusters (e.g., countries), Chi-square tests were performed for each category–cluster pair. For significant associations, Cramér’s V quantified effect size, and enrichment scores indicated over- or underrepresentation. P-values were adjusted using Benjamini–Hochberg false discovery rate (BH-FDR). For numerical features, Mann–Whitney U tests were applied to each feature–cluster pair, comparing within-cluster values to all others. Enrichment scores were derived from mean differences, with a small constant (ɛ = 1×10⁻⁴) added to avoid division by zero. Scores were log₂-normalized for scale-independent comparisons. BH-FDR-adjusted p-values were used to classify features as uniquely or multiply enriched. Visualization followed the same clustering strategy, with only significant log₂-normalized enrichments displayed. For both categorical and numerical enrichments, positive enrichments (PE) and negative enrichments (NE) and uniquely (U) or multiply (M) associated features with clusters were flagged in output tables (Table S2; S3; S8-10) and visualized accordingly.

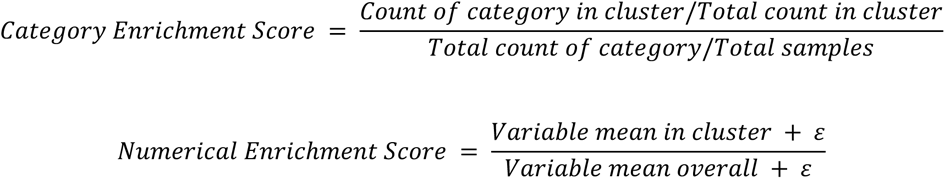

#### Analysis of sourdough aromas

Sensory diversity was assessed using Jaccard distances (full aroma spectrum). PERMANOVA was applied to the distance matrix to test categorical metadata effects, including only variables with ≥3 samples and applying BH-FDR correction. Relationships between sensory dissimilarities and numerical metadata were evaluated using Mantel tests (Pearson and Spearman) on scaled variables, with dynamic exclusion of missing data. Principal Coordinate Analysis (PCoA) was performed for visualization, retaining the first two axes (PC1, PC2). The top 10 aroma features explaining variance were identified via Pearson correlations with principal coordinates. Significant metadata were integrated into the PCoA plot, with scaled feature loadings shown as arrows.

#### Analysis of numerical metadata feature space

Principal Component Analysis (PCA) was performed using scikit-learn. Only non-collinear variables were considered; samples with missing values for the selected variables were excluded. PCA biplots were generated in matplotlib with confidence ellipses added at 95%.

## Data availability

The dataset supporting this study — including questionnaire files (Survey 1 and Survey 2), the home experiment instruction booklet, and the fully integrated metadata — is currently under restricted access to preserve the integrity of peer review. It will be made publicly available under Creative Commons Attribution 4.0 International (CC BY 4.0) license upon article publication. In the meantime, it is available upon request via Zenodo: https://doi.org/10.5281/zenodo.15366921.

## Code availability

Not applicable.

## Acknowledgements

The authors acknowledge financial support from the project HealthFerm, which is funded by the European Union under the Horizon Europe grant agreement No. 101060247 and by the Swiss State Secretariat for Education, Research and Innovation (SERI) under contract No. 22.00210. Views and opinions expressed are however those of the author(s) only and do not necessarily reflect those of the European Union nor European Research Executive Agency (REA). Neither the European Union nor REA can be held responsible for them.

The authors thank Alessandro Rearte, Rosy Mondardini, Wannes De Man, Michael Bom Frøst, Vimac Nolla Ardèvol, Jean-Paul Garin, Elena Martucci, and the rest of the HealthFerm consortium for their advice and support in setting up, promoting and advertising the project. We also thank Luisa Ferreira, Doriela Grabocka, Celine Verdonck, Silvia-Simona Grosu-Tudor, and Emanuela- Catalina Ionetic for their help in data curation and/or technical support. We offer our gratitude to all the participants who attended the co-design workshops, completed the questionnaires, submitted their sourdoughs, performed home experiments, and patiently waited for their results. None of this would have been possible without your contributions.

## Author information

### Contributions

Conceptualization: N.A.B., S.W., C.M.C., and R.C. conceived and designed the study with input from other authors. Methodology: A.M., T.G., J.P.T., F.C., Y.D.B., and N.A.B. designed the survey and experimental methodology with input from other authors. Investigation: A.M., T.G., J.P.T., C.V., I.-R.A., M.N., and A.Z.A.T. carried out data collection. Curation: A.M. and J.P.T. curated and managed the data. Formal analysis: A.M., J.P.T., and T.G. Visualization: A.M., T.G., and J.P.T. Supervision: N.A.B., L.D., M.G., M.Z., R.C., L.N., S.W., and C.M.C. Writing — original draft: A.M., T.G., J.P.T., and N.A.B. All authors contributed to the review and editing of the manuscript and approved the final version.

### Ethics declarations

This study obtained ethical approval for the activities of each respective hub lab. Approval was granted by the ethics committees of the respective universities under the reference numbers HealthFerm_Cod2023_14 (UniBZ), ECHW_429 (VUB), 183/2022 (IBB), 52/2023 (UH), and EK 2023-N-118 (ETHZ).

