## Supplementary information for "Rising together: Exploring Sourdough Fermentation Diversity through Co-design in the HealthFerm Citizen Science Initiative"

##### Subtitle: Sourdough Starter Characteristics, Baker Demographics, and Fermentation Practices across Europe

Annina Meyer*,[1], Thomas Gettemans*,[2], Jan Patrick Tan*,[3], Fabio Tuccillo[4], Chiara Viretto[5], Iulia-Roxana Angelescu[6], Yamina De Bondt[7], Michelle Neugebauer[1], Ali Zein Alabiden Tlais[5], Fabio Cavelti[1], Luc De Vuyst[2], Marco Gobbetti[5], Christophe M. Courtin[7], Medana Zamfir[6], Rossana Coda[4], Laura Nyström[3], Stefan Weckx[2], Nicholas A. Bokulich[1#]

*These authors contributed equally

Affiliations

1. Laboratory of Food Systems Biotechnology, Department of Health Sciences and Technology, ETH Zurich, Schmelzbergstrasse 7, 8092 Zurich, Switzerland
2. Research Group of Industrial Microbiology and Food Biotechnology (IMDO), Faculty of Sciences and Bioengineering Sciences, Vrije Universiteit Brussel (VUB), Pleinlaan 2, B-1050 Brussels, Belgium
3. Laboratory of Food Biochemistry, Department of Health Sciences and Technology, ETH Zurich, Schmelzbergstrasse 9, 8092 Zurich, Switzerland
4. Department of Food and Nutrition, University of Helsinki, Agnes Sjöbergin katu 2, 00014 Helsinki, Finland
5. Faculty of Agricultural, Environmental and Food Sciences, Free University of Bolzano-Bozen, 39100 Bolzano, Italy
6. Institute of Biology Bucharest of the Romanian Academy, 296 Splaiul Independentei, Bucharest, Romania
7. Laboratory of Food Chemistry and Biochemistry and Leuven Food Science and Nutrition Research Centre (LFoRCe), KU Leuven, Kasteelpark Arenberg 20, B-3001 Leuven, Belgium

#Correspondence


### Supplementary figures

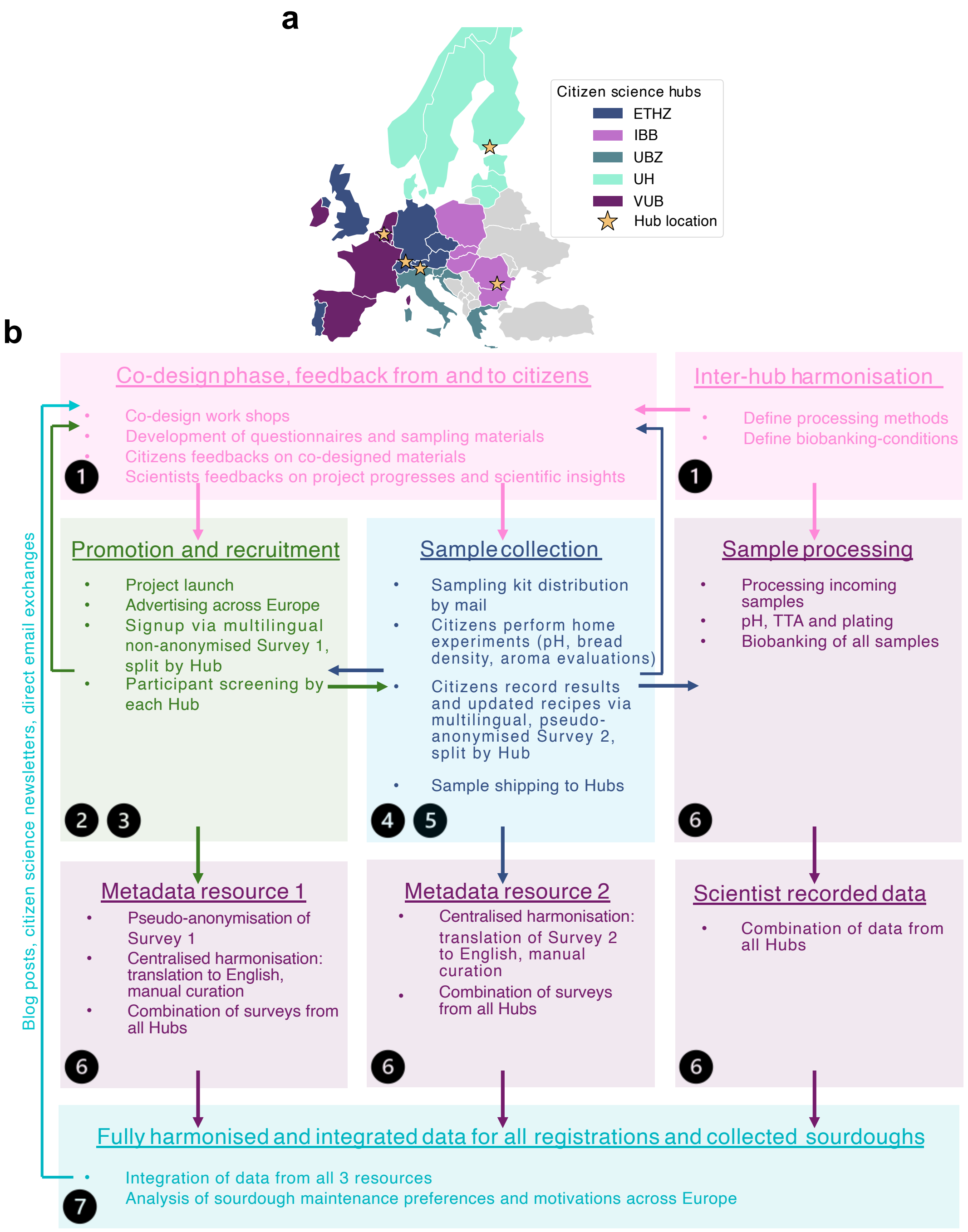

**Supplementary Figure 1. Methodological framework and project stages of the co-design-driven HealthFerm sourdough citizen science initiative. a**, Participating countries and laboratory hubs: ETHZ: ETH Zurich, Switzerland; IBB: Institute of Biology Bucharest, Romania; UBZ: Free University of Bozen-Bolzano, Italy; UH: University of Helsinki, Finland; VUB: Vrije Universiteit Brussel, Belgium. **b,** Schematic outlining the stepwise approach underlying the initiative, including: (1) the co-design phase, involving bidirectional feedback between citizens and scientists; (2–3) advertisement and participant recruitment via Survey 1; (4–5) sample collection and home-based experimental activities, home experiment result reporting via Survey 2; (6) unified sample processing across hubs and centralized harmonization of survey data; and (7) integrated survey data analysis. Communication and engagement channels (e.g., blog posts on the HealthFerm webpage, newsletters, and direct exchanges) supported participant involvement throughout the process.

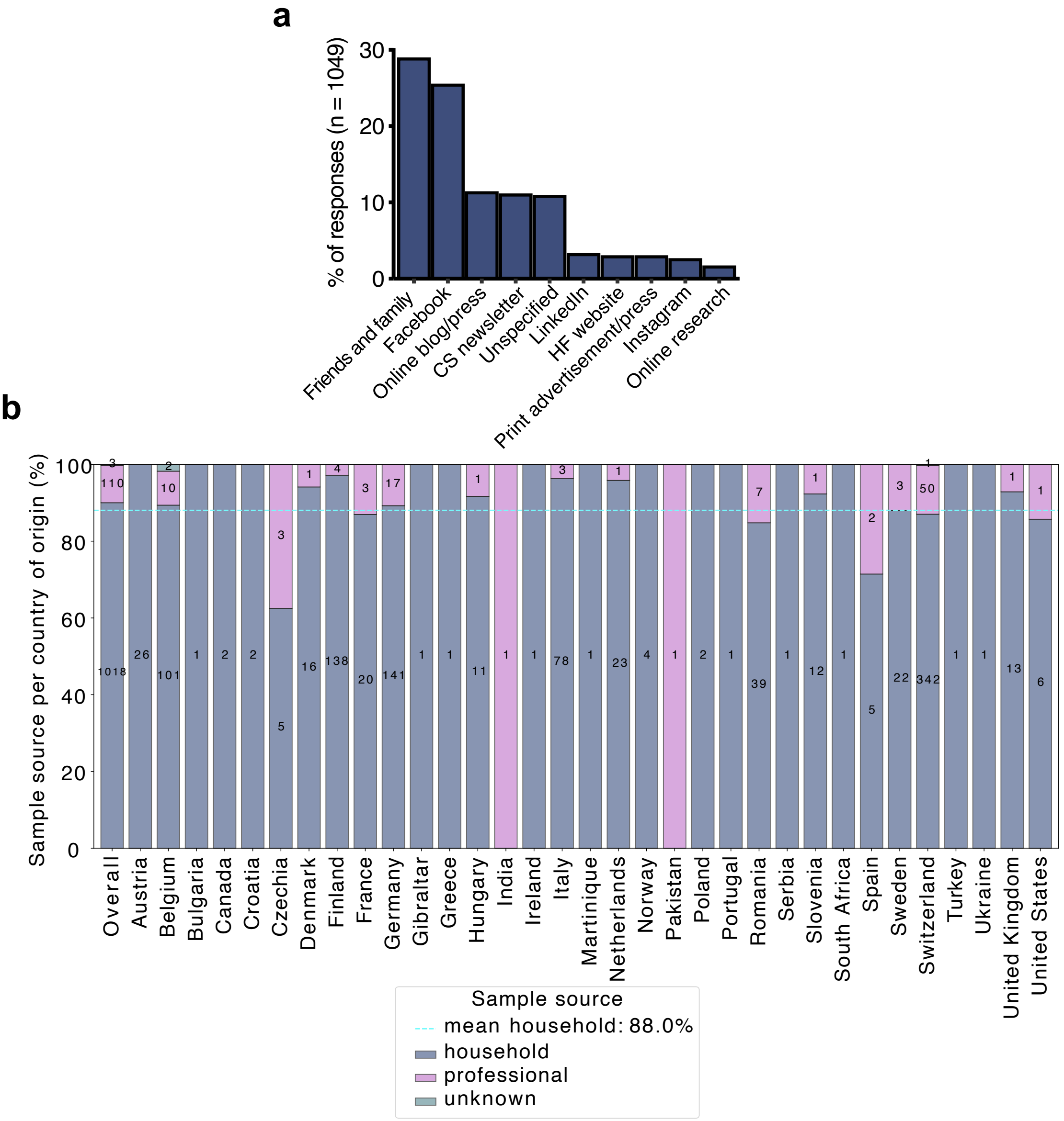

**Supplementary Figure 2. Advertisement platforms and signup distributions across countries. a**, Channels through which participants got in contact with the project team. **b**, Distribution of household and bakery samples across countries. The mean percentage of household samples across all countries is highlighted. Numbers within bars indicate absolute response counts per category.

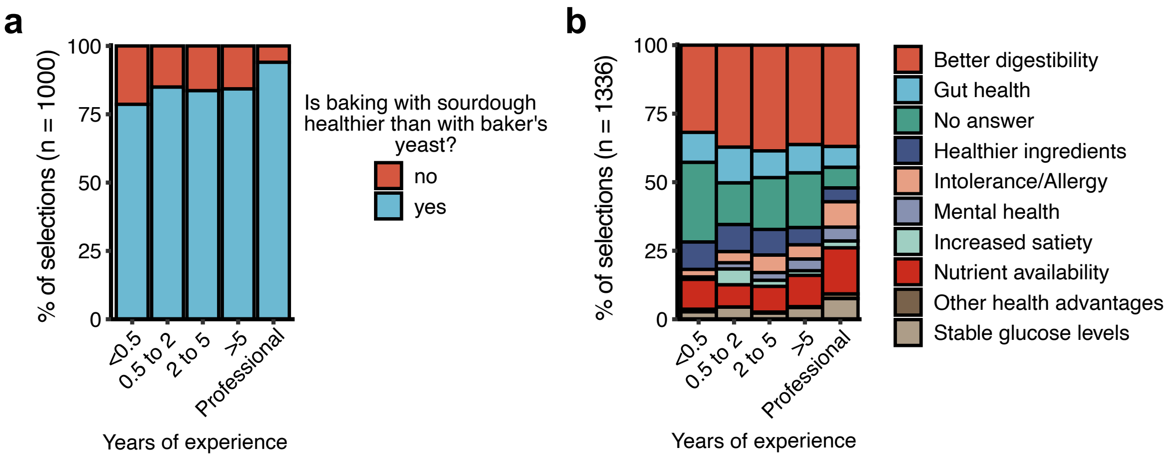

**Supplementary Figure 3. Perceived health benefits across skill levels**. **a**, Perception of sourdough bread being healthier than yeast-leavened bread, stratified by self-rated sourdough baking skill levels of all respondents. **b**, Perceived health benefits as specified by participants, stratified by self-rated sourdough baking skill levels of all respondents.

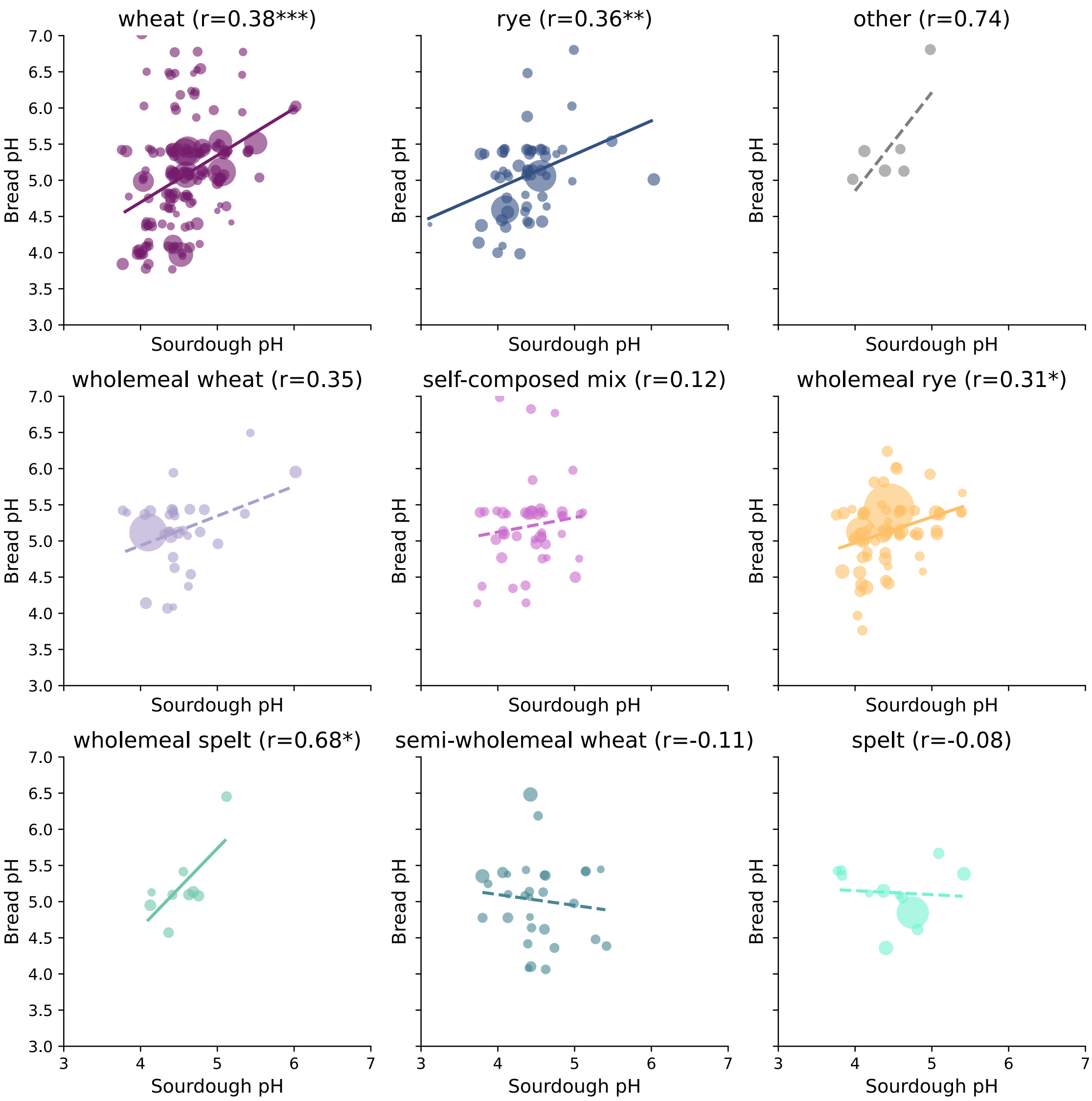

**Supplementary Figure 4. Sourdough and bread pH correlations split by flour type**. Pearson correlations between sourdough pH and sourdough bread pH as determined in home experiments, split by flour type. Significance of the correlation is marked according to p-values: *<0.05; **<0.01, ***<0.001. Bubble sizes are proportional to the bread density assessed by citizens ( bigg bubble size: high bread density; small bubble size: low bread density).

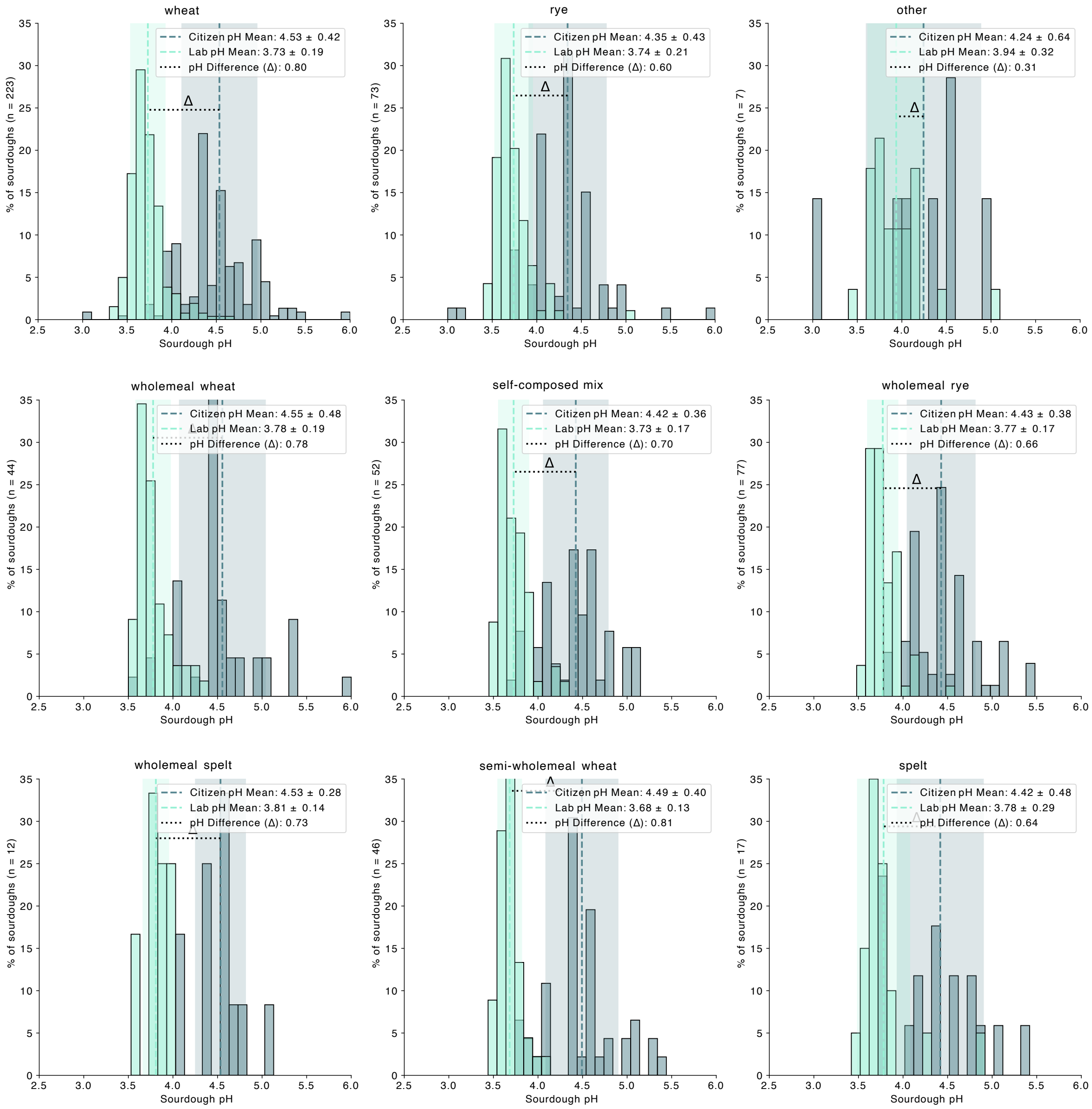

**Supplementary Figure 5. Distributions of at-home measured and in the laboratory measured sourdough pH**. pH shifts by sourdough flour type, comparing participant-measured pH at home with laboratory pH measurements after sample shipping.

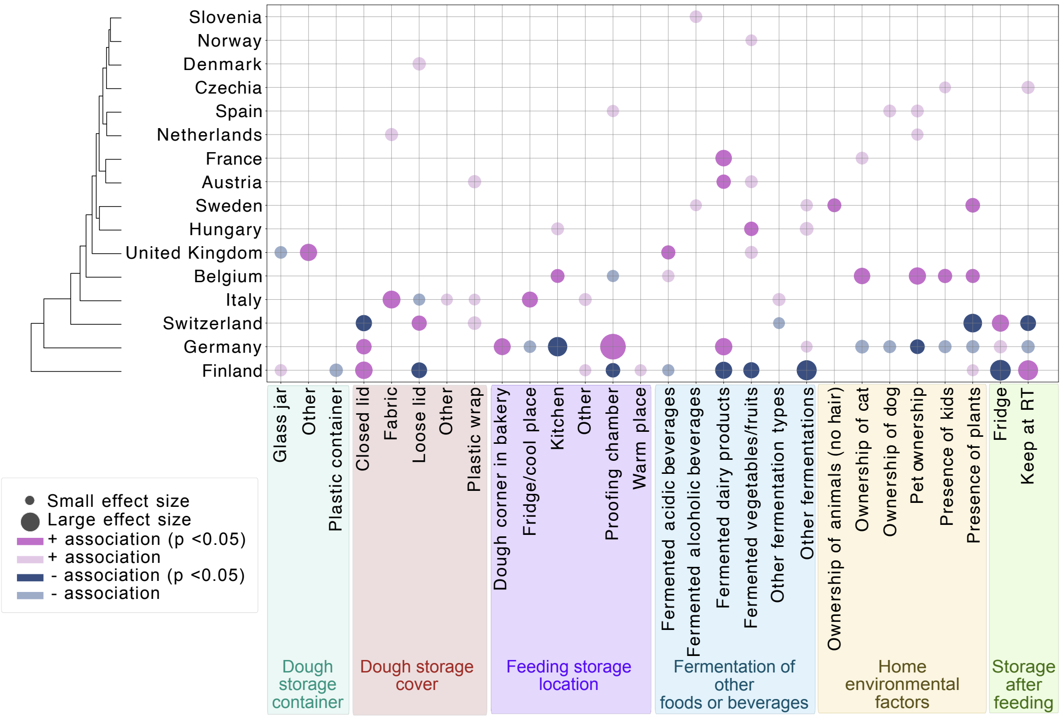

**Supplementary Figure 6. Sourdough maintenance practices and home environments among participants across European countries.** Bubble plot clustering countries based on similarities in sourdough maintenance practices and home environments. This includes dough storage devices and covers, feeding locations, whether participants ferment other foods or beverages, environmental factors, and where they store their sourdough after feeding. Only countries and subcategories with at least one significant Chi-square association across tested variables are displayed. Associations are color-coded based on their statistical significance (darker shade: BH-FDR corrected p-value < 0.05; lighter shade: uncorrected p-value < 0.05) and whether they show positive or negative enrichment within a given country, non-significant associations are not displayed. Countries are hierarchically clustered based on effect size (Cramér's V) across all features, including both significant and non-significant enrichments, and bubble size corresponds to effect size (Cramér's V).

### Supplementary Tables

**Supplementary Table 1.** Contents of the sampling kit and the purpose of each item.

| Items | Purpose |
|---|---|
| Instruction booklet | Provides an introduction to the project, graphical instructions for the sampling procedure and the at-home experiments along with a QR-code directing to the result questionnaire (deposited on Zenodo; https://doi.org/10.5281/zenodo.15366921) |
| Sampling container | A sterile container to avoid microbial contamination from participant-provided containers. |
| Plastic freezer bag | Provides a second barrier in case of sample leakage during shipment. |
| pH strips and flavour wheels | Required items for performing the at-home experiments. |
| Prepaid return shipment label | Facilitates sample shipment and covers any costs from the participant. |
| Project-branded pen and sticker | Souvenir for participation. |

**Supplementary Table 2.** Results of Chi-square enrichment tests for advertisement channels by country (corresponding to Figure 2E). p-values were corrected using the Benjamini–Hochberg false discovery rate (BH-FDR). Indicator codes: PE = positive enrichment, NE = negative enrichment, M = multiple indicators, U = unique indicator.

| *Category* | *Cluster* | *Chi2 Statistic* | *Cramér's V* | *Category count* | *Cluster Total* | *Category Total* | *Raw p-value* | *Corrected p-value* | *Enrichment Score* | *Interpretation* | *Indicator* |
|---|---|---|---|---|---|---|---|---|---|---|---|
| *bakery* | Belgium | 3.2207 | 0.0551 | 6 | 90 | 32 | 0.0727 | 0.1309 | 2.2104 | PE | M |
| *bakery* | Switzerland | 5.8903 | 0.0745 | 18 | 367 | 32 | 0.0152 | 0.0433 | 1.6262 | PE | M |
| *citizen science or uni newsletter* | Germany | 0.0473 | 0.0067 | 6 | 149 | 50 | 0.8278 | 0.9511 | 0.8545 | NE | M |
| *citizen science or uni newsletter* | Belgium | 1.3794 | 0.0361 | 7 | 90 | 50 | 0.2402 | 0.3706 | 1.6504 | PE | M |
| *citizen science or uni newsletter* | Austria | 4.5433 | 0.0654 | 4 | 26 | 50 | 0.0330 | 0.0744 | 3.2646 | PE | M |
| *citizen science or uni newsletter* | Switzerland | 4.8158 | 0.0674 | 25 | 367 | 50 | 0.0282 | 0.0662 | 1.4455 | PE | M |
| *citizen science or uni newsletter* | Slovenia | 6.1778 | 0.0763 | 3 | 13 | 50 | 0.0129 | 0.0411 | 4.8969 | PE | M |
| *direct email* | Germany | 4.9603 | 0.0684 | 14 | 149 | 56 | 0.0259 | 0.0637 | 1.7802 | PE | M |
| *direct email* | Belgium | 0.1365 | 0.0113 | 6 | 90 | 56 | 0.7118 | 0.8356 | 1.2631 | PE | M |
| *direct email* | Romania | 4.2898 | 0.0636 | 6 | 46 | 56 | 0.0383 | 0.0828 | 2.4713 | PE | M |
| *direct email* | Switzerland | 0.8161 | 0.0277 | 23 | 367 | 56 | 0.3663 | 0.5346 | 1.1874 | PE | M |
| *Facebook* | Finland | 360.2175 | 0.5827 | 126 | 139 | 267 | 0.0000 | 0.0000 | 3.6021 | PE | M |
| *Facebook* | Sweden | 11.3041 | 0.1032 | 14 | 25 | 267 | 0.0008 | 0.0032 | 2.2253 | PE | M |
| *Facebook* | Germany | 39.8325 | 0.1938 | 6 | 149 | 267 | 0.0000 | 0.0000 | 0.1600 | NE | M |
| *Facebook* | Netherlands | 12.5994 | 0.1090 | 14 | 24 | 267 | 0.0004 | 0.0017 | 2.3180 | PE | M |
| *Facebook* | Belgium | 0.6391 | 0.0245 | 19 | 90 | 267 | 0.4240 | 0.5871 | 0.8389 | NE | M |
| *Facebook* | France | 17.9115 | 0.1299 | 15 | 23 | 267 | 0.0000 | 0.0001 | 2.5916 | PE | M |
| *Facebook* | Austria | 0.2277 | 0.0146 | 5 | 26 | 267 | 0.6332 | 0.7787 | 0.7642 | NE | M |
| *Facebook* | Romania | 2.9258 | 0.0525 | 17 | 46 | 267 | 0.0872 | 0.1518 | 1.4686 | PE | M |
| *Facebook* | Switzerland | 117.4117 | 0.3327 | 19 | 367 | 267 | 0.0000 | 0.0000 | 0.2057 | NE | M |
| *Facebook* | Italy | 3.7341 | 0.0593 | 12 | 78 | 267 | 0.0533 | 0.0993 | 0.6114 | NE | M |
| *Facebook* | Slovenia | 0.0216 | 0.0045 | 4 | 13 | 267 | 0.8831 | 0.9873 | 1.2227 | PE | M |

| | | | | | | | | | | | |
|---|---|---|---|---|---|---|---|---|---|---|---|
| *Facebook* | Spain | 0.4164 | 0.0198 | 3 | 7 | 267 | 0.5187 | 0.7003 | 1.7030 | PE | M |
| *friends and family* | Finland | 28.9857 | 0.1653 | 9 | 139 | 269 | 0.0000 | 0.0000 | 0.2554 | NE | M |
| *friends and family* | Sweden | 0.7315 | 0.0263 | 4 | 25 | 269 | 0.3924 | 0.5576 | 0.6311 | NE | M |
| *friends and family* | Denmark | 0.2073 | 0.0140 | 3 | 17 | 269 | 0.6489 | 0.7787 | 0.6960 | NE | M |
| *friends and family* | United Kingdom | 5.9690 | 0.0750 | 8 | 14 | 269 | 0.0146 | 0.0433 | 2.2539 | PE | M |
| *friends and family* | Germany | 5.2462 | 0.0703 | 26 | 149 | 269 | 0.0220 | 0.0594 | 0.6883 | NE | M |
| *friends and family* | Belgium | 0.3447 | 0.0180 | 20 | 90 | 269 | 0.5571 | 0.7338 | 0.8765 | NE | M |
| *friends and family* | Czechia | 7.8586 | 0.0861 | 5 | 6 | 269 | 0.0051 | 0.0171 | 3.2869 | PE | M |
| *friends and family* | Romania | 4.0915 | 0.0621 | 18 | 46 | 269 | 0.0431 | 0.0875 | 1.5434 | PE | M |
| *friends and family* | Switzerland | 10.1301 | 0.0977 | 115 | 367 | 269 | 0.0015 | 0.0056 | 1.2359 | PE | M |
| *friends and family* | Hungary | 18.5717 | 0.1323 | 10 | 12 | 269 | 0.0000 | 0.0001 | 3.2869 | PE | M |
| *friends and family* | Italy | 2.3959 | 0.0475 | 26 | 78 | 269 | 0.1217 | 0.1991 | 1.3147 | PE | M |
| *friends and family* | Slovenia | 0.0171 | 0.0040 | 4 | 13 | 269 | 0.8959 | 0.9873 | 1.2136 | PE | M |
| *HealthFerm website* | Belgium | 5.1177 | 0.0695 | 7 | 90 | 34 | 0.0237 | 0.0609 | 2.4271 | PE | M |
| *HealthFerm website* | Austria | 17.0908 | 0.1269 | 5 | 26 | 34 | 0.0000 | 0.0002 | 6.0011 | PE | M |
| *HealthFerm website* | Switzerland | 1.8779 | 0.0421 | 16 | 367 | 34 | 0.1706 | 0.2709 | 1.3605 | PE | M |
| *Instagram* | Switzerland | 3.7585 | 0.0595 | 15 | 367 | 28 | 0.0525 | 0.0993 | 1.5488 | PE | U |
| *LinkedIn* | Denmark | 191.0073 | 0.4243 | 11 | 17 | 34 | 0.0000 | 0.0000 | 20.1920 | PE | M |
| *LinkedIn* | Switzerland | 0.2134 | 0.0142 | 10 | 367 | 34 | 0.6441 | 0.7787 | 0.8503 | NE | M |
| *online research* | Belgium | 8.0738 | 0.0872 | 5 | 90 | 16 | 0.0045 | 0.0162 | 3.6840 | PE | M |
| *online research* | Switzerland | 1.0836 | 0.0320 | 8 | 367 | 16 | 0.2979 | 0.4468 | 1.4455 | PE | M |
| *other* | Switzerland | 0.2888 | 0.0165 | 18 | 367 | 59 | 0.5910 | 0.7598 | 0.8820 | NE | M |
| *other* | Italy | 224.0940 | 0.4596 | 34 | 78 | 59 | 0.0000 | 0.0000 | 7.8388 | PE | M |
| *press* | Sweden | 2.7746 | 0.0511 | 6 | 25 | 122 | 0.0958 | 0.1616 | 2.0872 | PE | M |
| *press* | Belgium | 0.0027 | 0.0016 | 11 | 90 | 122 | 0.9583 | 1.0000 | 1.0629 | PE | M |
| *press* | Switzerland | 129.7522 | 0.3497 | 99 | 367 | 122 | 0.0000 | 0.0000 | 2.3460 | PE | M |
| *press* | Italy | 4.0670 | 0.0619 | 3 | 78 | 122 | 0.0437 | 0.0875 | 0.3345 | NE | M |
| *sourdough blog* | Germany | 577.7783 | 0.7379 | 91 | 149 | 94 | 0.0000 | 0.0000 | 6.8935 | PE | U |

**Supplementary Table 3.** Chi-square enrichment test results for baking motivations across flour types used (related to Figure 4C). p-values were corrected using the Benjamini–Hochberg false discovery rate (BH-FDR). Indicator codes: PE = positive enrichment, NE = negative enrichment, M = multiple indicators, U = unique indicator.

| ***Feature*** | ***Category*** | ***Cluster*** | ***Chi2 Statistic*** | ***Cramér's V*** | ***Category count*** | ***Cluster Total*** | ***Category Total*** | ***Raw p-value*** | ***Corrected p-value*** | ***Enrichment Score*** | ***Interpretation*** | ***Indicator*** |
|---|---|---|---|---|---|---|---|---|---|---|---|---|
| *Baking Motivation: Enjoy Baking* | yes | endosperm | 0.5807 | 0.0227 | 445 | 638 | 800 | 0.4460 | 0.6918 | 0.9861 | NE | M |
| *Baking Motivation: Enjoy Baking* | yes | mix | 6.3947 | 0.0752 | 116 | 145 | 800 | 0.0114 | 0.0870 | 1.1310 | PE | M |
| *Baking Motivation: Enjoy Baking* | yes | organic: no | 0.0313 | 0.0053 | 309 | 432 | 800 | 0.8596 | 0.8596 | 0.9942 | NE | M |
| *Baking Motivation: Enjoy Baking* | yes | other | 4.8249 | 0.0664 | 116 | 145 | 788 | 0.0281 | 0.1247 | 1.1107 | PE | M |
| *Baking Motivation: Enjoy Baking* | yes | other | 17.1397 | 0.1231 | 10 | 29 | 800 | 0.0000 | 0.0007 | 0.4875 | NE | M |
| *Baking Motivation: Enjoy Baking* | yes | rye | 0.0115 | 0.0032 | 212 | 296 | 788 | 0.9147 | 0.9628 | 0.9943 | NE | M |
| *Baking Motivation: Enjoy Baking* | yes | spelt | 5.3434 | 0.0699 | 29 | 51 | 788 | 0.0208 | 0.1234 | 0.7894 | NE | M |
| *Baking Motivation: Enjoy Baking* | yes | wheat | 0.0821 | 0.0087 | 431 | 602 | 788 | 0.7745 | 0.9628 | 0.9940 | NE | M |
| *Baking Motivation: Enjoy Baking* | yes | wholemeal | 0.1724 | 0.0123 | 229 | 319 | 800 | 0.6780 | 0.8437 | 1.0149 | PE | M |
| *Baking Motivation: Enjoy Baking* | yes | organic: yes | 0.0313 | 0.0053 | 491 | 680 | 800 | 0.8596 | 0.8596 | 1.0037 | PE | M |
| *Baking Motivation: Health* | yes | endosperm | 1.1958 | 0.0325 | 317 | 638 | 579 | 0.2742 | 0.5893 | 0.9706 | NE | M |
| *Baking Motivation: Health* | yes | mix | 0.0948 | 0.0092 | 72 | 145 | 579 | 0.7581 | 0.8730 | 0.9699 | NE | M |
| *Baking Motivation: Health* | yes | organic: no | 4.0970 | 0.0607 | 208 | 432 | 579 | 0.0430 | 0.2148 | 0.9247 | NE | M |
| *Baking Motivation: Health* | yes | other | 0.1812 | 0.0129 | 72 | 145 | 565 | 0.6703 | 0.8938 | 0.9615 | NE | M |
| *Baking Motivation: Health* | yes | other | 1.5859 | 0.0374 | 11 | 29 | 579 | 0.2079 | 0.5554 | 0.7409 | NE | M |
| *Baking Motivation: Health* | yes | rye | 10.3556 | 0.0973 | 177 | 296 | 565 | 0.0013 | 0.0129 | 1.1578 | PE | M |
| *Baking Motivation: Health* | yes | spelt | 0.2786 | 0.0160 | 24 | 51 | 565 | 0.5976 | 0.8333 | 0.9112 | NE | M |
| *Baking Motivation: Health* | yes | wheat | 5.0102 | 0.0677 | 292 | 602 | 565 | 0.0252 | 0.1247 | 0.9392 | NE | M |
| *Baking Motivation: Health* | yes | wholemeal | 4.0334 | 0.0597 | 179 | 319 | 579 | 0.0446 | 0.2825 | 1.0961 | PE | M |

| | | | | | | | | | | | | |
|---|---|---|---|---|---|---|---|---|---|---|---|---|
| *Baking Motivation: Health* | yes | organic: yes | 4.0970 | 0.0607 | 371 | 680 | 579 | 0.0430 | 0.2148 | 1.0478 | PE | M |
| *Baking Motivation: Lower Cost* | yes | endosperm | 0.4422 | 0.0198 | 87 | 638 | 162 | 0.5061 | 0.7439 | 0.9520 | NE | M |
| *Baking Motivation: Lower Cost* | yes | mix | 0.6890 | 0.0247 | 17 | 145 | 162 | 0.4065 | 0.6866 | 0.8185 | NE | M |
| *Baking Motivation: Lower Cost* | yes | organic: no | 0.2001 | 0.0134 | 66 | 432 | 162 | 0.6547 | 0.8510 | 1.0487 | PE | M |
| *Baking Motivation: Lower Cost* | yes | other | 0.8176 | 0.0273 | 17 | 145 | 159 | 0.3659 | 0.6418 | 0.8067 | NE | M |
| *Baking Motivation: Lower Cost* | yes | rye | 0.4515 | 0.0203 | 47 | 296 | 159 | 0.5016 | 0.7717 | 1.0925 | PE | M |
| *Baking Motivation: Lower Cost* | yes | spelt | 1.4043 | 0.0358 | 4 | 51 | 159 | 0.2360 | 0.6005 | 0.5396 | NE | M |
| *Baking Motivation: Lower Cost* | yes | wheat | 0.2688 | 0.0157 | 91 | 602 | 159 | 0.6042 | 0.8333 | 1.0401 | PE | M |
| *Baking Motivation: Lower Cost* | yes | wholemeal | 3.4224 | 0.0550 | 56 | 319 | 162 | 0.0643 | 0.2931 | 1.2256 | PE | M |
| *Baking Motivation: Lower Cost* | yes | organic: yes | 0.2001 | 0.0134 | 96 | 680 | 162 | 0.6547 | 0.8510 | 0.9691 | NE | M |
| *Baking Motivation: Taste* | yes | endosperm | 0.7697 | 0.0261 | 499 | 638 | 896 | 0.3803 | 0.6722 | 0.9873 | NE | M |
| *Baking Motivation: Taste* | yes | mix | 2.7965 | 0.0497 | 123 | 145 | 896 | 0.0945 | 0.3264 | 1.0708 | PE | M |
| *Baking Motivation: Taste* | yes | organic: no | 0.7548 | 0.0261 | 342 | 432 | 896 | 0.3850 | 0.6416 | 0.9825 | NE | M |
| *Baking Motivation: Taste* | yes | other | 1.5951 | 0.0382 | 123 | 145 | 882 | 0.2066 | 0.5903 | 1.0522 | PE | M |
| *Baking Motivation: Taste* | yes | other | 33.4556 | 0.1720 | 10 | 29 | 896 | 0.0000 | 0.0000 | 0.4353 | NE | M |
| *Baking Motivation: Taste* | yes | rye | 1.0181 | 0.0305 | 245 | 296 | 882 | 0.3130 | 0.6260 | 1.0267 | PE | M |
| *Baking Motivation: Taste* | yes | spelt | 4.1534 | 0.0616 | 35 | 51 | 882 | 0.0416 | 0.1511 | 0.8512 | NE | M |
| *Baking Motivation: Taste* | yes | wheat | 0.8068 | 0.0272 | 479 | 602 | 882 | 0.3691 | 0.6418 | 0.9869 | NE | M |
| *Baking Motivation: Taste* | yes | wholemeal | 3.0837 | 0.0522 | 264 | 319 | 896 | 0.0791 | 0.3005 | 1.0446 | PE | M |
| *Baking Motivation: Taste* | yes | organic: yes | 0.7548 | 0.0261 | 554 | 680 | 896 | 0.3850 | 0.6416 | 1.0111 | PE | M |
| *Baking Motivation: Tradition* | yes | endosperm | 3.2968 | 0.0540 | 54 | 638 | 81 | 0.0694 | 0.2931 | 1.1818 | PE | M |
| *Baking Motivation: Tradition* | yes | mix | 0.9900 | 0.0296 | 7 | 145 | 81 | 0.3197 | 0.6231 | 0.6741 | NE | M |
| *Baking Motivation: Tradition* | yes | organic: no | 0.8823 | 0.0282 | 27 | 432 | 81 | 0.3476 | 0.6416 | 0.8580 | NE | M |
| *Baking Motivation: Tradition* | yes | other | 1.1296 | 0.0321 | 7 | 145 | 80 | 0.2879 | 0.6260 | 0.6602 | NE | M |
| *Baking Motivation: Tradition* | yes | rye | 13.1166 | 0.1095 | 36 | 296 | 80 | 0.0003 | 0.0093 | 1.6632 | PE | M |
| *Baking Motivation: Tradition* | yes | spelt | 4.3140 | 0.0628 | 8 | 51 | 80 | 0.0378 | 0.1511 | 2.1451 | PE | M |

| | | | | | | | | | | | | |
|---|---|---|---|---|---|---|---|---|---|---|---|---|
| *Baking Motivation: Tradition* | yes | wheat | 11.4924 | 0.1025 | 29 | 602 | 80 | 0.0007 | 0.0093 | 0.6588 | NE | M |
| *Baking Motivation: Tradition* | yes | wholemeal | 0.3615 | 0.0179 | 20 | 319 | 81 | 0.5477 | 0.7633 | 0.8754 | NE | M |
| *Baking Motivation: Tradition* | yes | organic: yes | 0.8823 | 0.0282 | 54 | 680 | 81 | 0.3476 | 0.6416 | 1.0902 | PE | M |
| *Perceived Benefit: Better Digestibility* | yes | endosperm | 0.5907 | 0.0229 | 307 | 638 | 532 | 0.4421 | 0.6918 | 1.0230 | PE | M |
| *Perceived Benefit: Better Digestibility* | yes | mix | 0.1672 | 0.0122 | 71 | 145 | 532 | 0.6826 | 0.8437 | 1.0410 | PE | M |
| *Perceived Benefit: Better Digestibility* | yes | organic: no | 0.1530 | 0.0117 | 203 | 432 | 532 | 0.6956 | 0.8510 | 0.9822 | NE | M |
| *Perceived Benefit: Better Digestibility* | yes | other | 0.0135 | 0.0035 | 71 | 145 | 527 | 0.9075 | 0.9628 | 1.0165 | PE | M |
| *Perceived Benefit: Better Digestibility* | yes | other | 11.8702 | 0.1024 | 4 | 29 | 532 | 0.0006 | 0.0054 | 0.2932 | NE | M |
| *Perceived Benefit: Better Digestibility* | yes | rye | 0.0154 | 0.0038 | 144 | 296 | 527 | 0.9012 | 0.9628 | 1.0099 | PE | M |
| *Perceived Benefit: Better Digestibility* | yes | spelt | 0.0004 | 0.0006 | 24 | 51 | 527 | 0.9845 | 1.0000 | 0.9769 | NE | M |
| *Perceived Benefit: Better Digestibility* | yes | wheat | 0.0330 | 0.0055 | 288 | 602 | 527 | 0.8558 | 0.9628 | 0.9931 | NE | M |
| *Perceived Benefit: Better Digestibility* | yes | organic: yes | 0.1530 | 0.0117 | 329 | 680 | 532 | 0.6956 | 0.8510 | 1.0113 | PE | M |
| *Perceived Benefit: Gut Health* | yes | endosperm | 3.3650 | 0.0545 | 70 | 638 | 143 | 0.0666 | 0.2931 | 0.8678 | NE | M |
| *Perceived Benefit: Gut Health* | yes | mix | 12.4166 | 0.1048 | 32 | 145 | 143 | 0.0004 | 0.0054 | 1.7455 | PE | M |
| *Perceived Benefit: Gut Health* | yes | organic: no | 1.2381 | 0.0334 | 49 | 432 | 143 | 0.2658 | 0.6416 | 0.8820 | NE | M |
| *Perceived Benefit: Gut Health* | yes | other | 11.9370 | 0.1045 | 32 | 145 | 140 | 0.0006 | 0.0093 | 1.7245 | PE | M |
| *Perceived Benefit: Gut Health* | yes | rye | 1.6890 | 0.0393 | 31 | 296 | 140 | 0.1937 | 0.5903 | 0.8184 | NE | M |
| *Perceived Benefit: Gut Health* | yes | spelt | 0.7178 | 0.0256 | 9 | 51 | 140 | 0.3969 | 0.6615 | 1.3790 | PE | M |
| *Perceived Benefit: Gut Health* | yes | wheat | 2.4130 | 0.0470 | 68 | 602 | 140 | 0.1203 | 0.4011 | 0.8827 | NE | M |
| *Perceived Benefit: Gut Health* | yes | wholemeal | 0.0275 | 0.0049 | 39 | 319 | 143 | 0.8684 | 0.9072 | 0.9669 | NE | M |
| *Perceived Benefit: Gut Health* | yes | organic: yes | 1.2381 | 0.0334 | 94 | 680 | 143 | 0.2658 | 0.6416 | 1.0749 | PE | M |
| *Perceived Benefit: Healthier Ingredients* | yes | endosperm | 0.0215 | 0.0044 | 67 | 638 | 121 | 0.8833 | 0.9072 | 0.9816 | NE | M |
| *Perceived Benefit: Healthier Ingredients* | yes | mix | 0.3355 | 0.0172 | 13 | 145 | 121 | 0.5625 | 0.7633 | 0.8380 | NE | M |
| *Perceived Benefit: Healthier Ingredients* | yes | organic: no | 0.0887 | 0.0089 | 45 | 432 | 121 | 0.7659 | 0.8510 | 0.9573 | NE | M |
| *Perceived Benefit: Healthier Ingredients* | yes | other | 0.4709 | 0.0207 | 13 | 145 | 120 | 0.4926 | 0.7717 | 0.8174 | NE | M |

| | | | | | | | | | | | | |
|---|---|---|---|---|---|---|---|---|---|---|---|---|
| *Perceived Benefit: Healthier Ingredients* | yes | rye | 5.7720 | 0.0726 | 44 | 296 | 120 | 0.0163 | 0.1234 | 1.3552 | PE | M |
| *Perceived Benefit: Healthier Ingredients* | yes | spelt | 0.9236 | 0.0291 | 3 | 51 | 120 | 0.3365 | 0.6410 | 0.5363 | NE | M |
| *Perceived Benefit: Healthier Ingredients* | yes | wheat | 1.1579 | 0.0325 | 60 | 602 | 120 | 0.2819 | 0.6260 | 0.9086 | NE | M |
| *Perceived Benefit: Healthier Ingredients* | yes | wholeme al | 1.3188 | 0.0341 | 40 | 319 | 121 | 0.2508 | 0.5776 | 1.1721 | PE | M |
| *Perceived Benefit: Healthier Ingredients* | yes | organic: yes | 0.0887 | 0.0089 | 76 | 680 | 121 | 0.7659 | 0.8510 | 1.0271 | PE | M |
| *Perceived Benefit: Intolerance/Allergy* | yes | endosper m | 1.1340 | 0.0317 | 49 | 638 | 78 | 0.2869 | 0.5893 | 1.1136 | PE | M |
| *Perceived Benefit: Intolerance/Allergy* | yes | mix | 1.5092 | 0.0365 | 6 | 145 | 78 | 0.2193 | 0.5554 | 0.6000 | NE | M |
| *Perceived Benefit: Intolerance/Allergy* | yes | organic: no | 2.2294 | 0.0448 | 37 | 432 | 78 | 0.1354 | 0.4514 | 1.2210 | PE | M |
| *Perceived Benefit: Intolerance/Allergy* | yes | other | 1.3795 | 0.0355 | 6 | 145 | 74 | 0.2402 | 0.6005 | 0.6117 | NE | M |
| *Perceived Benefit: Intolerance/Allergy* | yes | other | 0.1378 | 0.0110 | 3 | 29 | 78 | 0.7105 | 0.8437 | 1.5000 | PE | M |
| *Perceived Benefit: Intolerance/Allergy* | yes | rye | 5.2786 | 0.0695 | 29 | 296 | 74 | 0.0216 | 0.1234 | 1.4484 | PE | M |
| *Perceived Benefit: Intolerance/Allergy* | yes | wheat | 1.0431 | 0.0309 | 36 | 602 | 74 | 0.3071 | 0.6260 | 0.8841 | NE | M |
| *Perceived Benefit: Intolerance/Allergy* | yes | wholeme al | 0.1530 | 0.0116 | 20 | 319 | 78 | 0.6957 | 0.8437 | 0.9091 | NE | M |
| *Perceived Benefit: Intolerance/Allergy* | yes | organic: yes | 2.2294 | 0.0448 | 41 | 680 | 78 | 0.1354 | 0.4514 | 0.8596 | NE | M |
| *Perceived Benefit: Nutrient Availability* | yes | endosper m | 2.1426 | 0.0435 | 78 | 638 | 154 | 0.1433 | 0.4188 | 0.8979 | NE | M |
| *Perceived Benefit: Nutrient Availability* | yes | mix | 0.0385 | 0.0058 | 21 | 145 | 154 | 0.8445 | 0.9072 | 1.0636 | PE | M |
| *Perceived Benefit: Nutrient Availability* | yes | organic: no | 15.8165 | 0.1193 | 37 | 432 | 154 | 0.0001 | 0.0007 | 0.6184 | NE | M |
| *Perceived Benefit: Nutrient Availability* | yes | other | 0.0257 | 0.0048 | 21 | 145 | 150 | 0.8726 | 0.9628 | 1.0563 | PE | M |
| *Perceived Benefit: Nutrient Availability* | yes | other | 0.0606 | 0.0073 | 3 | 29 | 154 | 0.8056 | 0.9004 | 0.7597 | NE | M |
| *Perceived Benefit: Nutrient Availability* | yes | rye | 0.1436 | 0.0115 | 43 | 296 | 150 | 0.7048 | 0.9094 | 1.0595 | PE | M |
| *Perceived Benefit: Nutrient Availability* | yes | spelt | 0.3873 | 0.0188 | 5 | 51 | 150 | 0.5337 | 0.7907 | 0.7150 | NE | M |
| *Perceived Benefit: Nutrient Availability* | yes | wheat | 0.0338 | 0.0056 | 81 | 602 | 150 | 0.8540 | 0.9628 | 0.9813 | NE | M |
| *Perceived Benefit: Nutrient Availability* | yes | wholeme al | 2.4140 | 0.0462 | 52 | 319 | 154 | 0.1203 | 0.3808 | 1.1972 | PE | M |
| *Perceived Benefit: Nutrient Availability* | yes | organic: yes | 15.8165 | 0.1193 | 117 | 680 | 154 | 0.0001 | 0.0007 | 1.2424 | PE | M |

**Supplementary Table 4.** PERMANOVA results based on the Jaccard distance matrix of aroma profiles, with variables ranked by explained variance ($R^2$) (related to Figure 5D).

| *Variable* | *Pseudo-F* | *R²* | *Sample Size* | *Number of Groups* | *Uncorrected p-value* | *Corrected p-value (BH-FDR)* |
|---|---|---|---|---|---|---|
| *plants* | 2.5745 | 0.0046 | 557 | 2 | 0.0020 | 0.0116 |
| *flour_organic* | 2.0455 | 0.0037 | 553 | 2 | 0.0020 | 0.0116 |
| *flour_quality* | 1.9707 | 0.0035 | 557 | 4 | 0.0010 | 0.0079 |
| *flour_type_red_red* | 1.8579 | 0.0034 | 550 | 4 | 0.0010 | 0.0079 |
| *acid_beverage* | 1.8652 | 0.0033 | 557 | 2 | 0.0070 | 0.0338 |
| *lower_cost* | 1.8248 | 0.0033 | 557 | 2 | 0.0090 | 0.0412 |
| *samp_source* | 1.8068 | 0.0032 | 557 | 3 | 0.0010 | 0.0079 |
| *tradition* | 1.7394 | 0.0031 | 557 | 2 | 0.0200 | 0.0757 |
| *sustainanbility* | 1.7141 | 0.0031 | 557 | 2 | 0.0130 | 0.0566 |
| *sample_received_season* | 1.6674 | 0.0030 | 557 | 4 | 0.0020 | 0.0116 |
| *flour_change_origin* | 0.9861 | 0.0030 | 331 | 131 | 0.7450 | 0.8102 |
| *flour_type_red* | 1.6309 | 0.0029 | 557 | 9 | 0.0010 | 0.0079 |
| *stor_feeding* | 1.5949 | 0.0029 | 557 | 6 | 0.0010 | 0.0079 |
| *kids* | 1.5851 | 0.0028 | 555 | 2 | 0.0450 | 0.1554 |
| *europe_region* | 1.5390 | 0.0028 | 553 | 4 | 0.0030 | 0.0154 |
| *no_pets* | 1.5412 | 0.0028 | 557 | 2 | 0.0500 | 0.1554 |
| *cheese* | 1.5134 | 0.0027 | 557 | 2 | 0.0470 | 0.1554 |
| *dairy_products* | 1.4868 | 0.0027 | 557 | 2 | 0.0520 | 0.1560 |
| *bread_refresh_loc* | 1.1126 | 0.0027 | 417 | 44 | 0.0020 | 0.0116 |
| *bread_baked* | 1.4641 | 0.0026 | 556 | 2 | 0.0680 | 0.1849 |
| *cs_info* | 1.3232 | 0.0026 | 517 | 13 | 0.0030 | 0.0154 |
| *fermented_cereals* | 1.4224 | 0.0025 | 557 | 2 | 0.0640 | 0.1796 |
| *samp_month* | 1.1011 | 0.0025 | 432 | 12 | 0.1400 | 0.3085 |
| *flour_type* | 1.3964 | 0.0025 | 557 | 15 | 0.0010 | 0.0079 |

| | | | | | | |
|---|---|---|---|---|---|---|
| *health_benefits_no_answer* | 1.3880 | 0.0025 | 557 | 2 | 0.0980 | 0.2508 |
| *alcoholic_beverage* | 1.3445 | 0.0024 | 557 | 2 | 0.1010 | 0.2511 |
| *dough_source* | 1.3317 | 0.0024 | 557 | 5 | 0.0200 | 0.0757 |
| *after_feeding* | 1.2735 | 0.0024 | 533 | 3 | 0.0940 | 0.2478 |
| *like_taste* | 1.3268 | 0.0024 | 557 | 2 | 0.1410 | 0.3085 |
| *dog* | 1.3182 | 0.0024 | 557 | 2 | 0.1560 | 0.3085 |
| *skill* | 1.2411 | 0.0023 | 528 | 5 | 0.0620 | 0.1796 |
| *hub* | 1.3026 | 0.0023 | 557 | 5 | 0.0230 | 0.0834 |
| *yeast_usage_prior* | 1.3009 | 0.0023 | 557 | 2 | 0.1550 | 0.3085 |
| *stable_glucose_level* | 1.2920 | 0.0023 | 557 | 2 | 0.1490 | 0.3085 |
| *samp_date* | 1.0825 | 0.0023 | 469 | 176 | 0.0010 | 0.0079 |
| *samp_country* | 1.2846 | 0.0023 | 557 | 19 | 0.0010 | 0.0079 |
| *beer* | 1.2845 | 0.0023 | 557 | 2 | 0.1530 | 0.3085 |
| *wholemeal_spelt* | 1.2839 | 0.0023 | 557 | 2 | 0.1290 | 0.3033 |
| *other_fermentations_yes_no* | 1.2171 | 0.0023 | 533 | 2 | 0.2160 | 0.3871 |
| *aroma_dough_difficulty* | 1.2398 | 0.0023 | 543 | 6 | 0.0200 | 0.0757 |
| *ph_measured* | 1.2268 | 0.0022 | 556 | 2 | 0.1900 | 0.3673 |
| *cow* | 1.2172 | 0.0022 | 557 | 2 | 0.1560 | 0.3085 |
| *lamas* | 1.2166 | 0.0022 | 557 | 2 | 0.0490 | 0.1554 |
| *fermented_vegetables* | 1.2118 | 0.0022 | 557 | 2 | 0.2020 | 0.3739 |
| *fermented_legumes* | 1.1959 | 0.0021 | 557 | 2 | 0.2320 | 0.4037 |
| *water_type* | 1.1823 | 0.0021 | 557 | 5 | 0.1260 | 0.3033 |
| *person_id* | 1.1782 | 0.0021 | 557 | 455 | 0.0010 | 0.0079 |
| *flour_change* | 1.1766 | 0.0021 | 557 | 2 | 0.2380 | 0.4060 |
| *hamster* | 1.1556 | 0.0021 | 557 | 2 | 0.2180 | 0.3871 |
| *intoleranaces_allergy* | 1.1519 | 0.0021 | 557 | 2 | 0.2430 | 0.4066 |
| *email_signup* | 1.0489 | 0.0020 | 521 | 2 | 0.3680 | 0.5426 |
| *fermented_roots* | 1.1122 | 0.0020 | 557 | 2 | 0.2010 | 0.3739 |

| | | | | | | |
|---|---|---|---|---|---|---|
| *cat* | 1.0993 | 0.0020 | 557 | 2 | 0.3230 | 0.5006 |
| *samp_city* | 1.0983 | 0.0020 | 557 | 357 | 0.0010 | 0.0079 |
| *covid* | 1.0971 | 0.0020 | 557 | 3 | 0.2970 | 0.4785 |
| *health* | 1.0903 | 0.0020 | 557 | 2 | 0.3280 | 0.5006 |
| *guinea_pig* | 1.0857 | 0.0019 | 557 | 2 | 0.3230 | 0.5006 |
| *baking_no_answer* | 1.0756 | 0.0019 | 557 | 2 | 0.2550 | 0.4186 |
| *fish* | 1.0500 | 0.0019 | 557 | 2 | 0.3390 | 0.5085 |
| *rabbit* | 1.0252 | 0.0018 | 557 | 2 | 0.4040 | 0.5858 |
| *mental_health* | 1.0246 | 0.0018 | 557 | 2 | 0.4220 | 0.6019 |
| *dough_stor_device* | 1.0116 | 0.0018 | 557 | 3 | 0.4580 | 0.6325 |
| *dough_cover* | 1.0052 | 0.0018 | 557 | 5 | 0.4500 | 0.6315 |
| *goat* | 0.9861 | 0.0018 | 557 | 2 | 0.5560 | 0.7329 |
| *nutrient_availability* | 0.9858 | 0.0018 | 557 | 2 | 0.4960 | 0.6743 |
| *flour_additions* | 0.9709 | 0.0017 | 557 | 3 | 0.5650 | 0.7337 |
| *bird* | 0.9608 | 0.0017 | 557 | 2 | 0.6340 | 0.7454 |
| *condiment* | 0.9455 | 0.0017 | 557 | 2 | 0.5490 | 0.7329 |
| *wine* | 0.9178 | 0.0016 | 557 | 2 | 0.5820 | 0.7446 |
| *chicken* | 0.9157 | 0.0016 | 557 | 2 | 0.5910 | 0.7452 |
| *hobby* | 0.9079 | 0.0016 | 557 | 2 | 0.6090 | 0.7454 |
| *gebril* | 0.9022 | 0.0016 | 557 | 2 | 0.7880 | 0.8464 |
| *reptiles* | 0.8919 | 0.0016 | 557 | 2 | 0.6300 | 0.7454 |
| *other_fermentations* | 0.8904 | 0.0016 | 557 | 2 | 0.6170 | 0.7454 |
| *fermented_fruits* | 0.8883 | 0.0016 | 557 | 2 | 0.6620 | 0.7578 |
| *business_job* | 0.8703 | 0.0016 | 557 | 2 | 0.6520 | 0.7563 |
| *like_baking* | 0.8575 | 0.0015 | 557 | 2 | 0.6290 | 0.7454 |
| *bees* | 0.8415 | 0.0015 | 557 | 2 | 0.7320 | 0.8102 |
| *others_health_adv* | 0.8271 | 0.0015 | 557 | 2 | 0.7400 | 0.8102 |
| *gut_health* | 0.7900 | 0.0014 | 557 | 2 | 0.7320 | 0.8102 |

| | | | | | | |
|---|---|---|---|---|---|---|
| *other_baking_reasons* | 0.7851 | 0.0014 | 557 | 2 | 0.7980 | 0.8467 |
| *more_satiety* | 0.7604 | 0.0014 | 557 | 2 | 0.8080 | 0.8469 |
| *horse* | 0.7432 | 0.0013 | 557 | 2 | 0.8910 | 0.9120 |
| *better_digestibility* | 0.7230 | 0.0013 | 557 | 2 | 0.8470 | 0.8773 |
| *healthier_ingredients* | 0.5923 | 0.0011 | 557 | 2 | 0.9520 | 0.9560 |
| *intolance_allergy* | 0.5907 | 0.0011 | 557 | 2 | 0.9560 | 0.9560 |

**Supplementary Table 5.** Mantel test results based on the Jaccard distance matrix of aroma profiles, with variables ranked by Pearson correlation coefficient ($R^2$) (related to Figure 5D). p-values were corrected using the Benjamini–Hochberg false discovery rate (BH-FDR).

| ***Numerical Variable*** | ***Number of samples*** | ***Mantel R (Pearson)*** | ***P-Value (Pearson)*** | ***Corrected P-Value (Pearson)*** | ***Mantel R (Spearman)*** | ***P-Value (Spearman)*** | ***Corrected P-Value (Spearman)*** |
|---|---|---|---|---|---|---|---|
| *number_sourdoughs* | 533 | 0.0666 | 0.0010 | 0.0165 | 0.0793 | 0.0010 | 0.0165 |
| *barley* | 557 | 0.0269 | 0.1050 | 0.3729 | 0.0229 | 0.2030 | 0.5913 |
| *rye* | 557 | 0.0211 | 0.0900 | 0.3729 | 0.0152 | 0.2950 | 0.6084 |
| *wheat* | 557 | 0.0201 | 0.0010 | 0.0165 | 0.0191 | 0.0010 | 0.0165 |
| *gluten_free_flour* | 557 | 0.0200 | 0.2260 | 0.5327 | -0.0091 | 0.6390 | 0.8168 |
| *tta_laboratory* | 541 | 0.0193 | 0.1320 | 0.3960 | 0.0168 | 0.1150 | 0.3960 |
| *time_feeding* | 553 | 0.0180 | 0.2700 | 0.5940 | 0.0076 | 0.5810 | 0.8168 |
| *longitude* | 557 | 0.0165 | 0.2900 | 0.5981 | 0.0046 | 0.7340 | 0.8352 |
| *temp_feeding* | 557 | 0.0130 | 0.3540 | 0.6872 | 0.0220 | 0.0830 | 0.3913 |
| *ph_value_citizen* | 544 | 0.0089 | 0.4720 | 0.8653 | 0.0056 | 0.6350 | 0.8168 |
| *spelt* | 557 | 0.0087 | 0.5710 | 0.9422 | -0.0032 | 0.8540 | 0.8960 |
| *bread_density* | 446 | 0.0073 | 0.7010 | 0.9529 | 0.0291 | 0.0430 | 0.3548 |
| *wholemeal_rye* | 557 | 0.0050 | 0.6650 | 0.9529 | -0.0031 | 0.8170 | 0.8960 |
| *dough_age* | 556 | 0.0030 | 0.8370 | 0.9529 | 0.0117 | 0.3560 | 0.6782 |
| *refreshment_amount* | 557 | 0.0027 | 0.8860 | 0.9529 | -0.0116 | 0.4110 | 0.6782 |
| *samp_year* | 556 | 0.0027 | 0.8630 | 0.9529 | 0.0097 | 0.4540 | 0.7134 |
| *refreshment_flour* | 557 | 0.0018 | 0.8980 | 0.9529 | -0.0248 | 0.0750 | 0.3913 |

| | | | | | | | |
|---|---|---|---|---|---|---|---|
| *refreshment_water* | 557 | 0.0017 | 0.9170 | 0.9529 | -0.0249 | 0.0690 | 0.3913 |
| *latitude* | 557 | 0.0012 | 0.9240 | 0.9529 | 0.0049 | 0.6870 | 0.8168 |
| *other_wheat_varieties* | 557 | -0.0007 | 0.9760 | 0.9760 | -0.0026 | 0.8960 | 0.8960 |
| *wholemeal_wheat* | 557 | -0.0028 | 0.8370 | 0.9529 | -0.0060 | 0.6930 | 0.8168 |
| *dough_yield* | 555 | -0.0039 | 0.8030 | 0.9529 | 0.0022 | 0.8850 | 0.8960 |
| *elevation* | 557 | -0.0041 | 0.6870 | 0.9529 | -0.0049 | 0.5720 | 0.8168 |
| *temp_storage* | 557 | -0.0070 | 0.6260 | 0.9529 | -0.0055 | 0.6850 | 0.8168 |
| *pH_laboratory* | 544 | -0.0177 | 0.1820 | 0.5005 | -0.0145 | 0.2670 | 0.6084 |
| *bread_making_frequency* | 520 | -0.0197 | 0.2050 | 0.5204 | -0.0173 | 0.2410 | 0.6084 |
| *semi-wholemeal_wheat* | 557 | -0.0241 | 0.1040 | 0.3729 | -0.0269 | 0.1140 | 0.3960 |
| *sample_transport_dur* | 526 | -0.0254 | 0.1130 | 0.3729 | 0.0147 | 0.2150 | 0.5913 |
| *bread_ph* | 444 | -0.0297 | 0.0580 | 0.3190 | -0.0124 | 0.3850 | 0.6782 |
| *dough_size_feeding* | 557 | -0.0338 | 0.0550 | 0.3190 | -0.0148 | 0.2770 | 0.6084 |
| *backslop_freq* | 554 | -0.0447 | 0.0020 | 0.0220 | -0.0398 | 0.0050 | 0.0550 |

**Supplementary Table 6.** Pearson and Spearman correlations, as well as cosine similarity, between perceived aromas and measurable sourdough characteristics, fermentation parameters, and substrate proportions per sourdough (related to Figure 5E–F). p-values were corrected using the Benjamini–Hochberg false discovery rate (BH-FDR).

| ***Variable 1*** | ***Variable 2*** | ***Pearson Correlation*** | ***Spearman Correlation*** | ***Cosine Similarity*** | ***Uncorrected Pearson P-Value*** | ***Corrected Pearson P-Value*** | ***Uncorrected Spearman P-Value*** | ***Corrected Spearman P-Value*** |
|---|---|---|---|---|---|---|---|---|
| *Chemical* | bread_density | -0.0529 | -0.0120 | 0.1779 | 0.2646 | 0.6083 | 0.8007 | 0.9132 |
| *Chemical* | dough_age | -0.0334 | -0.0200 | 0.1232 | 0.4316 | 0.7512 | 0.6374 | 0.8274 |
| *Chemical* | dough_yield | 0.0178 | -0.0333 | 0.4013 | 0.6752 | 0.8637 | 0.4330 | 0.6924 |
| *Chemical* | pH_laboratory | 0.0027 | -0.0073 | 0.4084 | 0.9507 | 0.9690 | 0.8648 | 0.9472 |
| *Chemical* | backslop_freq | -0.0608 | -0.0655 | 0.2290 | 0.1529 | 0.4941 | 0.1234 | 0.3561 |
| *Chemical* | barley | -0.0191 | 0.0139 | 0.0021 | 0.6534 | 0.8573 | 0.7426 | 0.8870 |
| *Chemical* | bread_ph | -0.0808 | -0.1042 | 0.4005 | 0.0890 | 0.3569 | 0.0281 | 0.1240 |
| *Chemical* | dough_size_feeding | 0.0416 | 0.0540 | 0.1356 | 0.3272 | 0.6586 | 0.2031 | 0.4653 |
| *Chemical* | ph_value_citizen | -0.1017 | -0.1070 | 0.3939 | 0.0177 | 0.1127 | 0.0125 | 0.0693 |

| | | | | | | | | |
|---|---|---|---|---|---|---|---|---|
| *Chemical* | rye | -0.0689 | -0.0599 | 0.1263 | 0.1041 | 0.4000 | 0.1581 | 0.3963 |
| *Chemical* | semi-wholemeal_wheat | 0.0218 | 0.0236 | 0.1391 | 0.6075 | 0.8397 | 0.5785 | 0.7897 |
| *Chemical* | spelt | 0.0574 | 0.0849 | 0.1522 | 0.1759 | 0.5269 | 0.0452 | 0.1791 |
| *Chemical* | temp_feeding | -0.0343 | -0.0372 | 0.3886 | 0.4190 | 0.7353 | 0.3813 | 0.6475 |
| *Chemical* | temp_storage | -0.0107 | -0.0016 | 0.3227 | 0.8008 | 0.9062 | 0.9696 | 0.9900 |
| *Chemical* | time_feeding | -0.0321 | 0.0108 | 0.2249 | 0.4514 | 0.7524 | 0.7996 | 0.9132 |
| *Chemical* | wheat | 0.1212 | 0.1237 | 0.3710 | 0.0042 | 0.0330 | 0.0035 | 0.0240 |
| *Chemical* | wholemeal_rye | -0.0816 | -0.0973 | 0.1269 | 0.0544 | 0.2477 | 0.0216 | 0.1065 |
| *Chemical* | wholemeal_spelt | 0.0217 | 0.0316 | 0.0767 | 0.6090 | 0.8397 | 0.4563 | 0.7081 |
| *Chemical* | wholemeal_wheat | -0.0528 | -0.0638 | 0.1163 | 0.2131 | 0.5581 | 0.1328 | 0.3719 |
| *Chemical* | tta_laboratory | -0.1353 | -0.1396 | 0.3404 | 0.0016 | 0.0144 | 0.0011 | 0.0089 |
| *Earthy* | bread_density | -0.0188 | 0.0020 | 0.1936 | 0.6924 | 0.8657 | 0.9671 | 0.9890 |
| *Earthy* | dough_age | -0.0314 | -0.0467 | 0.1148 | 0.4597 | 0.7600 | 0.2720 | 0.5439 |
| *Earthy* | dough_yield | 0.0030 | 0.0262 | 0.3752 | 0.9443 | 0.9690 | 0.5372 | 0.7640 |
| *Earthy* | pH_laboratory | 0.0330 | 0.0430 | 0.3807 | 0.4420 | 0.7524 | 0.3170 | 0.5925 |
| *Earthy* | backslop_freq | 0.0159 | -0.0191 | 0.2625 | 0.7094 | 0.8746 | 0.6529 | 0.8388 |
| *Earthy* | barley | -0.0181 | 0.0235 | 0.0016 | 0.6704 | 0.8637 | 0.5805 | 0.7897 |
| *Earthy* | bread_ph | -0.0578 | -0.0600 | 0.3788 | 0.2243 | 0.5721 | 0.2067 | 0.4702 |
| *Earthy* | dough_size_feeding | -0.0074 | -0.0266 | 0.0857 | 0.8615 | 0.9294 | 0.5316 | 0.7594 |
| *Earthy* | ph_value_citizen | -0.0055 | 0.0018 | 0.3775 | 0.8987 | 0.9523 | 0.9664 | 0.9890 |
| *Earthy* | rye | -0.0825 | -0.0383 | 0.1026 | 0.0518 | 0.2452 | 0.3667 | 0.6356 |
| *Earthy* | semi-wholemeal_wheat | -0.0161 | -0.0005 | 0.0980 | 0.7045 | 0.8720 | 0.9903 | 0.9976 |
| *Earthy* | spelt | 0.0449 | 0.0592 | 0.1351 | 0.2897 | 0.6360 | 0.1631 | 0.4062 |
| *Earthy* | temp_feeding | -0.0297 | -0.0118 | 0.3639 | 0.4844 | 0.7805 | 0.7804 | 0.9080 |
| *Earthy* | temp_storage | -0.0139 | -0.0608 | 0.2997 | 0.7439 | 0.8872 | 0.1519 | 0.3906 |
| *Earthy* | time_feeding | 0.0675 | 0.0310 | 0.2817 | 0.1127 | 0.4203 | 0.4670 | 0.7176 |
| *Earthy* | wheat | 0.0352 | 0.0321 | 0.2983 | 0.4071 | 0.7217 | 0.4496 | 0.7046 |
| *Earthy* | wholemeal_rye | -0.0179 | -0.0078 | 0.1655 | 0.6734 | 0.8637 | 0.8539 | 0.9422 |
| *Earthy* | wholemeal_spelt | 0.0447 | 0.0487 | 0.0943 | 0.2926 | 0.6400 | 0.2512 | 0.5382 |
| *Earthy* | wholemeal_wheat | 0.0439 | 0.0621 | 0.1876 | 0.3008 | 0.6404 | 0.1433 | 0.3794 |
| *Earthy* | tta_laboratory | -0.0030 | -0.0372 | 0.3554 | 0.9444 | 0.9690 | 0.3877 | 0.6498 |

| | | | | | | | | |
|---|---|---|---|---|---|---|---|---|
| *Fermented* | bread_density | -0.0975 | -0.0533 | 0.3188 | 0.0396 | 0.2093 | 0.2612 | 0.5410 |
| *Fermented* | dough_age | -0.0808 | -0.0584 | 0.2115 | 0.0570 | 0.2529 | 0.1687 | 0.4116 |
| *Fermented* | dough_yield | 0.0054 | -0.0387 | 0.6980 | 0.8994 | 0.9523 | 0.3624 | 0.6324 |
| *Fermented* | pH_laboratory | 0.0533 | 0.0489 | 0.7095 | 0.2148 | 0.5592 | 0.2553 | 0.5410 |
| *Fermented* | backslop_freq | 0.0531 | 0.0602 | 0.4982 | 0.2123 | 0.5581 | 0.1572 | 0.3963 |
| *Fermented* | barley | -0.0467 | -0.0574 | 0.0011 | 0.2707 | 0.6177 | 0.1762 | 0.4213 |
| *Fermented* | bread_ph | 0.0118 | 0.0298 | 0.7123 | 0.8045 | 0.9062 | 0.5308 | 0.7594 |
| *Fermented* | dough_size_feed ing | 0.1002 | 0.0589 | 0.2407 | 0.0181 | 0.1138 | 0.1654 | 0.4102 |
| *Fermented* | ph_value_citizen | 0.0233 | 0.0237 | 0.7119 | 0.5872 | 0.8393 | 0.5817 | 0.7897 |
| *Fermented* | rye | 0.0574 | 0.0547 | 0.3545 | 0.1762 | 0.5269 | 0.1973 | 0.4536 |
| *Fermented* | semi-wholemeal_whe at | 0.1274 | 0.1188 | 0.2952 | 0.0026 | 0.0218 | 0.0050 | 0.0331 |
| *Fermented* | spelt | 0.0452 | 0.0626 | 0.2077 | 0.2873 | 0.6348 | 0.1401 | 0.3771 |
| *Fermented* | temp_feeding | -0.1156 | -0.1246 | 0.6716 | 0.0063 | 0.0461 | 0.0032 | 0.0225 |
| *Fermented* | temp_storage | -0.0258 | -0.0304 | 0.5621 | 0.5440 | 0.8239 | 0.4734 | 0.7221 |
| *Fermented* | time_feeding | 0.0367 | 0.1025 | 0.4514 | 0.3887 | 0.7021 | 0.0159 | 0.0823 |
| *Fermented* | wheat | -0.0292 | -0.0263 | 0.5002 | 0.4921 | 0.7909 | 0.5358 | 0.7637 |
| *Fermented* | wholemeal_rye | -0.0645 | -0.0455 | 0.2956 | 0.1283 | 0.4489 | 0.2840 | 0.5590 |
| *Fermented* | wholemeal_spelt | 0.0415 | 0.0460 | 0.1286 | 0.3278 | 0.6586 | 0.2784 | 0.5522 |
| *Fermented* | wholemeal_whe at | -0.0321 | -0.0402 | 0.2594 | 0.4496 | 0.7524 | 0.3431 | 0.6106 |
| *Fermented* | tta_laboratory | -0.0846 | -0.0863 | 0.6430 | 0.0492 | 0.2404 | 0.0448 | 0.1786 |
| *Fermented dairy* | bread_density | -0.0747 | -0.1565 | 0.3280 | 0.1152 | 0.4269 | 0.0009 | 0.0077 |
| *Fermented dairy* | dough_age | 0.0521 | 0.0609 | 0.2991 | 0.2197 | 0.5628 | 0.1515 | 0.3906 |
| *Fermented dairy* | dough_yield | -0.0444 | -0.0407 | 0.6940 | 0.2966 | 0.6404 | 0.3381 | 0.6087 |
| *Fermented dairy* | pH_laboratory | 0.0112 | 0.0029 | 0.7093 | 0.7948 | 0.9062 | 0.9463 | 0.9838 |
| *Fermented dairy* | backslop_freq | 0.1793 | 0.1542 | 0.5666 | 0.0000 | 0.0004 | 0.0003 | 0.0029 |
| *Fermented dairy* | barley | -0.0462 | -0.0248 | 0.0016 | 0.2760 | 0.6210 | 0.5590 | 0.7823 |
| *Fermented dairy* | bread_ph | -0.0335 | -0.0525 | 0.7023 | 0.4818 | 0.7795 | 0.2692 | 0.5439 |
| *Fermented dairy* | dough_size_feed ing | 0.0973 | 0.0404 | 0.2390 | 0.0217 | 0.1301 | 0.3413 | 0.6095 |
| *Fermented dairy* | ph_value_citizen | -0.0495 | -0.0472 | 0.7044 | 0.2487 | 0.6002 | 0.2717 | 0.5439 |

| | | | | | | | | |
|---|---|---|---|---|---|---|---|---|
| *Fermented dairy* | rye | -0.0253 | -0.0060 | 0.3034 | 0.5514 | 0.8251 | 0.8873 | 0.9602 |
| *Fermented dairy* | semi-wholemeal_wheat | 0.0217 | 0.0160 | 0.2245 | 0.6090 | 0.8397 | 0.7066 | 0.8577 |
| *Fermented dairy* | spelt | -0.0465 | -0.0351 | 0.1457 | 0.2735 | 0.6198 | 0.4077 | 0.6637 |
| *Fermented dairy* | temp_feeding | -0.0377 | -0.0357 | 0.6861 | 0.3744 | 0.6917 | 0.4008 | 0.6575 |
| *Fermented dairy* | temp_storage | 0.0508 | 0.0049 | 0.5955 | 0.2316 | 0.5828 | 0.9084 | 0.9730 |
| *Fermented dairy* | time_feeding | 0.0197 | 0.0020 | 0.4431 | 0.6442 | 0.8526 | 0.9630 | 0.9881 |
| *Fermented dairy* | wheat | 0.0834 | 0.0737 | 0.5563 | 0.0490 | 0.2404 | 0.0822 | 0.2655 |
| *Fermented dairy* | wholemeal_rye | -0.0083 | 0.0231 | 0.3316 | 0.8444 | 0.9203 | 0.5858 | 0.7903 |
| *Fermented dairy* | wholemeal_spelt | -0.0197 | -0.0166 | 0.0861 | 0.6425 | 0.8526 | 0.6956 | 0.8554 |
| *Fermented dairy* | wholemeal_wheat | 0.0463 | 0.0773 | 0.3109 | 0.2751 | 0.6210 | 0.0683 | 0.2379 |
| *Fermented dairy* | tta_laboratory | -0.0582 | -0.0676 | 0.6512 | 0.1763 | 0.5269 | 0.1165 | 0.3473 |
| *Fruity* | bread_density | -0.0683 | -0.0529 | 0.2604 | 0.1501 | 0.4898 | 0.2649 | 0.5419 |
| *Fruity* | dough_age | 0.0026 | -0.0688 | 0.2208 | 0.9511 | 0.9690 | 0.1049 | 0.3240 |
| *Fruity* | dough_yield | -0.0437 | -0.0025 | 0.5705 | 0.3046 | 0.6419 | 0.9525 | 0.9870 |
| *Fruity* | pH_laboratory | 0.0249 | 0.0215 | 0.5830 | 0.5627 | 0.8342 | 0.6175 | 0.8156 |
| *Fruity* | backslop_freq | -0.0264 | -0.0171 | 0.3746 | 0.5354 | 0.8188 | 0.6886 | 0.8554 |
| *Fruity* | barley | 0.0419 | 0.0755 | 0.0620 | 0.3239 | 0.6586 | 0.0749 | 0.2523 |
| *Fruity* | bread_ph | -0.0157 | -0.0330 | 0.5876 | 0.7421 | 0.8872 | 0.4877 | 0.7333 |
| *Fruity* | dough_size_feeding | -0.0105 | 0.0741 | 0.1342 | 0.8050 | 0.9062 | 0.0806 | 0.2646 |
| *Fruity* | ph_value_citizen | -0.0412 | -0.0376 | 0.5836 | 0.3374 | 0.6658 | 0.3813 | 0.6475 |
| *Fruity* | rye | -0.0230 | -0.0163 | 0.2469 | 0.5875 | 0.8393 | 0.7005 | 0.8577 |
| *Fruity* | semi-wholemeal_wheat | -0.0385 | -0.0224 | 0.1435 | 0.3644 | 0.6783 | 0.5985 | 0.7972 |
| *Fruity* | spelt | -0.0423 | -0.0182 | 0.1132 | 0.3191 | 0.6586 | 0.6686 | 0.8479 |
| *Fruity* | temp_feeding | 0.0069 | 0.0290 | 0.5725 | 0.8706 | 0.9322 | 0.4943 | 0.7383 |
| *Fruity* | temp_storage | -0.0216 | -0.0567 | 0.4639 | 0.6116 | 0.8397 | 0.1818 | 0.4290 |
| *Fruity* | time_feeding | 0.0874 | -0.0049 | 0.4135 | 0.0399 | 0.2093 | 0.9078 | 0.9730 |
| *Fruity* | wheat | 0.0601 | 0.0560 | 0.4593 | 0.1564 | 0.5004 | 0.1867 | 0.4358 |

| | | | | | | | | |
|---|---|---|---|---|---|---|---|---|
| *Fruity* | wholemeal_rye | 0.0050 | -0.0113 | 0.2815 | 0.9055 | 0.9540 | 0.7908 | 0.9132 |
| *Fruity* | wholemeal_spelt | -0.0583 | -0.0583 | 0.0357 | 0.1697 | 0.5241 | 0.1692 | 0.4116 |
| *Fruity* | wholemeal_wheat | -0.0549 | -0.0325 | 0.1911 | 0.1957 | 0.5316 | 0.4446 | 0.7002 |
| *Fruity* | tta_laboratory | -0.0068 | -0.0410 | 0.5436 | 0.8748 | 0.9325 | 0.3415 | 0.6095 |
| *Grain/Cereal* | bread_density | -0.0262 | -0.0731 | 0.3869 | 0.5816 | 0.8366 | 0.1233 | 0.3561 |
| *Grain/Cereal* | dough_age | 0.0265 | 0.0742 | 0.2972 | 0.5335 | 0.8188 | 0.0806 | 0.2646 |
| *Grain/Cereal* | dough_yield | 0.0558 | 0.1133 | 0.7505 | 0.1895 | 0.5311 | 0.0075 | 0.0448 |
| *Grain/Cereal* | pH_laboratory | 0.0722 | 0.0844 | 0.7532 | 0.0924 | 0.3685 | 0.0491 | 0.1876 |
| *Grain/Cereal* | backslop_freq | 0.0331 | 0.0960 | 0.5169 | 0.4364 | 0.7512 | 0.0239 | 0.1131 |
| *Grain/Cereal* | barley | -0.0487 | -0.0233 | 0.0042 | 0.2510 | 0.6014 | 0.5837 | 0.7903 |
| *Grain/Cereal* | bread_ph | 0.0302 | -0.0086 | 0.7552 | 0.5253 | 0.8151 | 0.8561 | 0.9429 |
| *Grain/Cereal* | dough_size_feeding | -0.0172 | -0.0272 | 0.1721 | 0.6850 | 0.8657 | 0.5214 | 0.7569 |
| *Grain/Cereal* | ph_value_citizen | 0.0791 | 0.0629 | 0.7591 | 0.0651 | 0.2808 | 0.1427 | 0.3794 |
| *Grain/Cereal* | rye | -0.1300 | -0.1027 | 0.2624 | 0.0021 | 0.0185 | 0.0153 | 0.0805 |
| *Grain/Cereal* | semi-wholemeal_wheat | -0.0413 | -0.0419 | 0.1968 | 0.3305 | 0.6586 | 0.3231 | 0.5969 |
| *Grain/Cereal* | spelt | -0.0341 | 0.0124 | 0.1664 | 0.4214 | 0.7374 | 0.7698 | 0.9032 |
| *Grain/Cereal* | temp_feeding | -0.0384 | -0.1168 | 0.7285 | 0.3661 | 0.6783 | 0.0058 | 0.0367 |
| *Grain/Cereal* | temp_storage | -0.0179 | -0.0297 | 0.6025 | 0.6735 | 0.8637 | 0.4836 | 0.7333 |
| *Grain/Cereal* | time_feeding | -0.0603 | -0.0172 | 0.4271 | 0.1565 | 0.5004 | 0.6870 | 0.8554 |
| *Grain/Cereal* | wheat | -0.0277 | -0.0438 | 0.5348 | 0.5149 | 0.8057 | 0.3018 | 0.5761 |
| *Grain/Cereal* | wholemeal_rye | 0.0620 | 0.0847 | 0.3931 | 0.1442 | 0.4802 | 0.0457 | 0.1799 |
| *Grain/Cereal* | wholemeal_spelt | -0.0569 | -0.0560 | 0.0689 | 0.1800 | 0.5269 | 0.1868 | 0.4358 |
| *Grain/Cereal* | wholemeal_wheat | 0.0853 | 0.0871 | 0.3496 | 0.0442 | 0.2227 | 0.0398 | 0.1654 |
| *Grain/Cereal* | tta_laboratory | -0.0195 | 0.0233 | 0.7003 | 0.6517 | 0.8571 | 0.5886 | 0.7906 |
| *Nutty* | bread_density | -0.0564 | -0.0525 | 0.0988 | 0.2343 | 0.5843 | 0.2686 | 0.5439 |

| | | | | | | | | |
|---|---|---|---|---|---|---|---|---|
| *Nutty* | dough_age | -0.0304 | -0.0161 | 0.0728 | 0.4738 | 0.7736 | 0.7056 | 0.8577 |
| *Nutty* | dough_yield | 0.0376 | 0.0827 | 0.2673 | 0.3769 | 0.6926 | 0.0515 | 0.1943 |
| *Nutty* | pH_laboratory | -0.0588 | -0.0404 | 0.2647 | 0.1710 | 0.5249 | 0.3470 | 0.6123 |
| *Nutty* | backslop_freq | 0.0123 | 0.0233 | 0.1874 | 0.7721 | 0.8958 | 0.5849 | 0.7903 |
| *Nutty* | barley | -0.0093 | 0.0585 | 0.0039 | 0.8268 | 0.9186 | 0.1676 | 0.4116 |
| *Nutty* | bread_ph | -0.0479 | -0.0904 | 0.2684 | 0.3136 | 0.6519 | 0.0569 | 0.2073 |
| *Nutty* | dough_size_feeding | -0.0170 | 0.0252 | 0.0493 | 0.6896 | 0.8657 | 0.5527 | 0.7772 |
| *Nutty* | ph_value_citizen | 0.0242 | 0.0201 | 0.2722 | 0.5732 | 0.8365 | 0.6393 | 0.8274 |
| *Nutty* | rye | -0.0481 | -0.0245 | 0.0791 | 0.2573 | 0.6021 | 0.5636 | 0.7823 |
| *Nutty* | semi-wholemeal_wheat | 0.0351 | 0.0296 | 0.1115 | 0.4081 | 0.7217 | 0.4860 | 0.7333 |
| *Nutty* | spelt | -0.0486 | 0.0138 | 0.0216 | 0.2523 | 0.6021 | 0.7448 | 0.8870 |
| *Nutty* | temp_feeding | 0.0263 | 0.0072 | 0.2668 | 0.5350 | 0.8188 | 0.8657 | 0.9472 |
| *Nutty* | temp_storage | 0.0536 | 0.0485 | 0.2471 | 0.2062 | 0.5482 | 0.2531 | 0.5397 |
| *Nutty* | time_feeding | -0.0199 | 0.0315 | 0.1485 | 0.6399 | 0.8526 | 0.4604 | 0.7126 |
| *Nutty* | wheat | 0.1030 | 0.0972 | 0.2629 | 0.0150 | 0.0973 | 0.0218 | 0.1065 |
| *Nutty* | wholemeal_rye | -0.0325 | -0.0052 | 0.0995 | 0.4441 | 0.7524 | 0.9034 | 0.9712 |
| *Nutty* | wholemeal_spelt | 0.0105 | 0.0105 | 0.0477 | 0.8047 | 0.9062 | 0.8047 | 0.9132 |
| *Nutty* | wholemeal_wheat | -0.0050 | -0.0044 | 0.1015 | 0.9054 | 0.9540 | 0.9172 | 0.9730 |
| *Nutty* | tta_laboratory | -0.0087 | -0.0031 | 0.2484 | 0.8400 | 0.9203 | 0.9429 | 0.9837 |
| *Sour* | bread_density | 0.0446 | 0.1060 | 0.3371 | 0.3478 | 0.6661 | 0.0252 | 0.1150 |
| *Sour* | dough_age | -0.0228 | 0.0024 | 0.1941 | 0.5912 | 0.8397 | 0.9557 | 0.9871 |
| *Sour* | dough_yield | -0.0137 | 0.0149 | 0.5556 | 0.7467 | 0.8872 | 0.7259 | 0.8711 |
| *Sour* | pH_laboratory | -0.0045 | 0.0041 | 0.5697 | 0.9170 | 0.9578 | 0.9241 | 0.9752 |
| *Sour* | backslop_freq | 0.0086 | -0.0238 | 0.3809 | 0.8399 | 0.9203 | 0.5768 | 0.7897 |
| *Sour* | barley | 0.0474 | 0.1039 | 0.0662 | 0.2640 | 0.6083 | 0.0141 | 0.0759 |
| *Sour* | bread_ph | -0.0464 | -0.0377 | 0.5729 | 0.3290 | 0.6586 | 0.4277 | 0.6857 |
| *Sour* | dough_size_feeding | 0.0451 | 0.0584 | 0.1738 | 0.2882 | 0.6348 | 0.1686 | 0.4116 |
| *Sour* | ph_value_citizen | -0.0896 | -0.0805 | 0.5611 | 0.0367 | 0.2012 | 0.0606 | 0.2181 |
| *Sour* | rye | 0.0114 | 0.0063 | 0.2632 | 0.7882 | 0.9062 | 0.8827 | 0.9588 |
| *Sour* | semi-wholemeal_wheat | -0.0579 | -0.0471 | 0.1220 | 0.1725 | 0.5249 | 0.2668 | 0.5439 |
| *Sour* | spelt | 0.0404 | 0.0861 | 0.1738 | 0.3411 | 0.6661 | 0.0423 | 0.1707 |

| | | | | | | | | |
|---|---|---|---|---|---|---|---|---|
| *Sour* | temp_feeding | 0.0399 | -0.0075 | 0.5597 | 0.3475 | 0.6661 | 0.8594 | 0.9449 |
| *Sour* | temp_storage | -0.0251 | 0.0114 | 0.4463 | 0.5540 | 0.8261 | 0.7879 | 0.9125 |
| *Sour* | time_feeding | 0.0046 | 0.0415 | 0.3494 | 0.9148 | 0.9578 | 0.3301 | 0.6010 |
| *Sour* | wheat | -0.1338 | -0.1487 | 0.3359 | 0.0015 | 0.0143 | 0.0004 | 0.0044 |
| *Sour* | wholemeal_rye | 0.1294 | 0.1386 | 0.3626 | 0.0022 | 0.0190 | 0.0010 | 0.0084 |
| *Sour* | wholemeal_spelt | 0.0319 | 0.0312 | 0.1057 | 0.4523 | 0.7524 | 0.4625 | 0.7141 |
| *Sour* | wholemeal_wheat | -0.0523 | -0.0668 | 0.1846 | 0.2175 | 0.5628 | 0.1153 | 0.3458 |
| *Sour* | tta_laboratory | 0.1207 | 0.1331 | 0.5690 | 0.0049 | 0.0380 | 0.0019 | 0.0146 |
| *Toasted* | bread_density | -0.0471 | -0.0500 | 0.1293 | 0.3209 | 0.6586 | 0.2918 | 0.5639 |
| *Toasted* | dough_age | -0.0213 | 0.0290 | 0.1016 | 0.6162 | 0.8397 | 0.4957 | 0.7383 |
| *Toasted* | dough_yield | 0.0485 | 0.1271 | 0.3260 | 0.2541 | 0.6021 | 0.0027 | 0.0194 |
| *Toasted* | pH_laboratory | -0.0422 | -0.0152 | 0.3211 | 0.3262 | 0.6586 | 0.7236 | 0.8711 |
| *Toasted* | backslop_freq | 0.0769 | -0.0156 | 0.2691 | 0.0705 | 0.3002 | 0.7133 | 0.8625 |
| *Toasted* | barley | 0.1067 | 0.0381 | 0.1164 | 0.0117 | 0.0802 | 0.3692 | 0.6356 |
| *Toasted* | bread_ph | -0.0199 | -0.0310 | 0.3177 | 0.6757 | 0.8637 | 0.5148 | 0.7508 |
| *Toasted* | dough_size_feeding | -0.0272 | 0.0305 | 0.0534 | 0.5218 | 0.8117 | 0.4728 | 0.7221 |
| *Toasted* | ph_value_citizen | -0.0444 | -0.0538 | 0.3208 | 0.3009 | 0.6404 | 0.2100 | 0.4758 |
| *Toasted* | rye | -0.0149 | 0.0155 | 0.1323 | 0.7248 | 0.8867 | 0.7149 | 0.8628 |
| *Toasted* | semi-wholemeal_wheat | 0.0827 | 0.0838 | 0.1701 | 0.0511 | 0.2437 | 0.0482 | 0.1852 |
| *Toasted* | spelt | -0.0176 | -0.0005 | 0.0643 | 0.6784 | 0.8637 | 0.9915 | 0.9976 |
| *Toasted* | temp_feeding | 0.0284 | 0.0462 | 0.3201 | 0.5035 | 0.7969 | 0.2765 | 0.5512 |
| *Toasted* | temp_storage | 0.0484 | -0.0046 | 0.2876 | 0.2544 | 0.6021 | 0.9139 | 0.9730 |
| *Toasted* | time_feeding | 0.0462 | 0.0613 | 0.2317 | 0.2779 | 0.6215 | 0.1501 | 0.3891 |
| *Toasted* | wheat | -0.0058 | 0.0037 | 0.2303 | 0.8912 | 0.9472 | 0.9309 | 0.9791 |
| *Toasted* | wholemeal_rye | -0.0374 | -0.0170 | 0.1216 | 0.3783 | 0.6928 | 0.6888 | 0.8554 |
| *Toasted* | wholemeal_spelt | 0.0646 | 0.0368 | 0.1059 | 0.1277 | 0.4489 | 0.3863 | 0.6498 |
| *Toasted* | wholemeal_wheat | -0.0434 | -0.0367 | 0.0898 | 0.3070 | 0.6426 | 0.3878 | 0.6498 |
| *Toasted* | tta_laboratory | 0.0303 | -0.0342 | 0.3136 | 0.4824 | 0.7795 | 0.4270 | 0.6857 |
| *Veggie* | bread_density | -0.0280 | 0.0744 | 0.2107 | 0.5547 | 0.8261 | 0.1169 | 0.3473 |
| *Veggie* | dough_age | 0.0219 | -0.0287 | 0.1849 | 0.6057 | 0.8397 | 0.4988 | 0.7397 |
| *Veggie* | dough_yield | -0.0112 | 0.0300 | 0.4386 | 0.7916 | 0.9062 | 0.4803 | 0.7308 |
| *Veggie* | pH_laboratory | 0.0605 | 0.0739 | 0.4447 | 0.1591 | 0.5036 | 0.0850 | 0.2718 |

| | | | | | | | | |
|---|---|---|---|---|---|---|---|---|
| *Veggie* | backslop_freq | -0.0803 | -0.0921 | 0.2425 | 0.0588 | 0.2592 | 0.0303 | 0.1324 |
| *Veggie* | barley | -0.0240 | -0.0455 | 0.0000 | 0.5718 | 0.8365 | 0.2837 | 0.5590 |
| *Veggie* | bread_ph | -0.0162 | -0.0320 | 0.4308 | 0.7328 | 0.8872 | 0.5008 | 0.7397 |
| *Veggie* | dough_size_feeding | 0.0230 | 0.1038 | 0.1285 | 0.5873 | 0.8393 | 0.0142 | 0.0759 |
| *Veggie* | ph_value_citizen | 0.0028 | -0.0042 | 0.4463 | 0.9482 | 0.9690 | 0.9227 | 0.9752 |
| *Veggie* | rye | -0.0084 | -0.0094 | 0.1940 | 0.8435 | 0.9203 | 0.8252 | 0.9218 |
| *Veggie* | semi-wholemeal_wheat | -0.0196 | -0.0137 | 0.1152 | 0.6439 | 0.8526 | 0.7477 | 0.8888 |
| *Veggie* | spelt | -0.0147 | 0.0024 | 0.0988 | 0.7291 | 0.8872 | 0.9546 | 0.9871 |
| *Veggie* | temp_feeding | -0.0560 | -0.0331 | 0.4238 | 0.1866 | 0.5271 | 0.4352 | 0.6925 |
| *Veggie* | temp_storage | -0.0778 | -0.0451 | 0.3202 | 0.0667 | 0.2858 | 0.2881 | 0.5634 |
| *Veggie* | time_feeding | -0.0362 | -0.0039 | 0.2452 | 0.3953 | 0.7096 | 0.9279 | 0.9775 |
| *Veggie* | wheat | -0.0217 | -0.0072 | 0.3110 | 0.6089 | 0.8397 | 0.8660 | 0.9472 |
| *Veggie* | wholemeal_rye | 0.0664 | 0.0480 | 0.2640 | 0.1176 | 0.4313 | 0.2579 | 0.5410 |
| *Veggie* | wholemeal_spelt | 0.0236 | 0.0189 | 0.0836 | 0.5790 | 0.8366 | 0.6564 | 0.8388 |
| *Veggie* | wholemeal_wheat | -0.0321 | -0.0219 | 0.1502 | 0.4492 | 0.7524 | 0.6068 | 0.8031 |
| *Veggie* | tta_laboratory | 0.0038 | 0.0135 | 0.4167 | 0.9291 | 0.9643 | 0.7532 | 0.8920 |

**Supplementary Table 7.** Pearson and Spearman correlations, as well as cosine similarity, between measurable sourdough characteristics, home experiment results, fermentation parameters, and substrate proportions per sourdough (related to Figure 6E). p-values were corrected using the Benjamini–Hochberg false discovery rate (BH-FDR).

| ***Variable 1*** | ***Variable 2*** | ***Pearson Correlation*** | ***Spearman Correlation*** | ***Cosine Similarity*** | ***Uncorrected Pearson P-Value*** | ***Corrected Pearson P-Value*** | ***Uncorrected Spearman P-Value*** | ***Corrected Spearman P-Value*** |
|---|---|---|---|---|---|---|---|---|
| *bread_density* | dough_age | -0.0170 | 0.0071 | 0.1880 | 0.7213 | 0.8844 | 0.8815 | 0.9588 |
| *bread_density* | dough_yield | 0.0089 | 0.1312 | 0.5179 | 0.8524 | 0.9243 | 0.0056 | 0.0364 |
| *bread_density* | pH_laboratory | 0.0485 | 0.0432 | 0.5254 | 0.3126 | 0.6519 | 0.3692 | 0.6356 |
| *bread_density* | backslop_freq | -0.0691 | -0.2610 | 0.3081 | 0.1463 | 0.4802 | 0.0000 | 0.0000 |
| *bread_density* | barley | -0.0182 | -0.0730 | 0.0180 | 0.7020 | 0.8720 | 0.1238 | 0.3561 |
| *bread_density* | bread_ph | -0.0023 | 0.0266 | 0.5177 | 0.9614 | 0.9706 | 0.5787 | 0.7897 |
| *bread_density* | dough_size_feeding | -0.0194 | -0.0203 | 0.1040 | 0.6833 | 0.8657 | 0.6689 | 0.8479 |
| *bread_density* | ph_value_citizen | -0.0049 | -0.0888 | 0.5202 | 0.9181 | 0.9578 | 0.0621 | 0.2212 |

| | | | | | | | | |
|---|---|---|---|---|---|---|---|---|
| *bread_density* | rye | 0.0284 | 0.1110 | 0.2604 | 0.5495 | 0.8251 | 0.0190 | 0.0964 |
| *bread_density* | semi-wholemeal_wheat | -0.0659 | -0.1419 | 0.0883 | 0.1646 | 0.5159 | 0.0027 | 0.0194 |
| *bread_density* | spelt | 0.0349 | 0.1040 | 0.1612 | 0.4626 | 0.7609 | 0.0281 | 0.1240 |
| *bread_density* | temp_feeding | 0.0117 | -0.0266 | 0.5155 | 0.8054 | 0.9062 | 0.5754 | 0.7897 |
| *bread_density* | temp_storage | 0.0654 | 0.0941 | 0.4589 | 0.1679 | 0.5237 | 0.0470 | 0.1828 |
| *bread_density* | time_feeding | -0.0336 | 0.0861 | 0.2895 | 0.4804 | 0.7795 | 0.0703 | 0.2425 |
| *bread_density* | wheat | -0.0694 | -0.3064 | 0.3443 | 0.1435 | 0.4802 | 0.0000 | 0.0000 |
| *bread_density* | wholemeal_rye | 0.1042 | 0.2153 | 0.3289 | 0.0279 | 0.1567 | 0.0000 | 0.0001 |
| *bread_density* | wholemeal_spelt | -0.0154 | 0.0653 | 0.0616 | 0.7453 | 0.8872 | 0.1684 | 0.4116 |
| *bread_density* | wholemeal_wheat | -0.0156 | -0.0067 | 0.1915 | 0.7427 | 0.8872 | 0.8881 | 0.9602 |
| *bread_density* | tta_laboratory | 0.0166 | 0.2313 | 0.5007 | 0.7305 | 0.8872 | 0.0000 | 0.0000 |
| *dough_age* | bread_density | -0.0170 | 0.0071 | 0.1880 | 0.7213 | 0.8844 | 0.8815 | 0.9588 |
| *dough_age* | dough_yield | 0.0184 | 0.0476 | 0.3702 | 0.6651 | 0.8633 | 0.2636 | 0.5410 |
| *dough_age* | pH_laboratory | 0.0439 | -0.0795 | 0.4767 | 0.3068 | 0.6426 | 0.0642 | 0.2257 |
| *dough_age* | backslop_freq | 0.0759 | 0.0383 | 0.2983 | 0.0745 | 0.3116 | 0.3688 | 0.6356 |
| *dough_age* | barley | -0.0188 | -0.0867 | 0.0005 | 0.6583 | 0.8587 | 0.0410 | 0.1668 |
| *dough_age* | bread_ph | 0.0097 | -0.0192 | 0.3755 | 0.8389 | 0.9203 | 0.6875 | 0.8554 |
| *dough_age* | dough_size_feeding | 0.0364 | -0.0128 | 0.1234 | 0.3915 | 0.7047 | 0.7631 | 0.8992 |
| *dough_age* | ph_value_citizen | -0.0419 | 0.0137 | 0.3621 | 0.3294 | 0.6586 | 0.7503 | 0.8902 |
| *dough_age* | rye | -0.0164 | 0.0045 | 0.1541 | 0.6989 | 0.8720 | 0.9160 | 0.9730 |
| *dough_age* | semi-wholemeal_wheat | 0.0136 | -0.0630 | 0.1223 | 0.7492 | 0.8872 | 0.1377 | 0.3771 |
| *dough_age* | spelt | -0.0544 | -0.1531 | 0.0443 | 0.2001 | 0.5386 | 0.0003 | 0.0031 |
| *dough_age* | temp_feeding | 0.0198 | 0.0162 | 0.3674 | 0.6418 | 0.8526 | 0.7023 | 0.8577 |
| *dough_age* | temp_storage | 0.0701 | 0.0368 | 0.3398 | 0.0989 | 0.3869 | 0.3871 | 0.6498 |
| *dough_age* | time_feeding | -0.0002 | 0.0030 | 0.2267 | 0.9965 | 0.9965 | 0.9447 | 0.9837 |
| *dough_age* | wheat | 0.0022 | -0.0106 | 0.2720 | 0.9579 | 0.9706 | 0.8032 | 0.9132 |

| | | | | | | | | |
|---|---|---|---|---|---|---|---|---|
| *dough_age* | wholemeal_rye | 0.0214 | 0.0943 | 0.1944 | 0.6151 | 0.8397 | 0.0262 | 0.1188 |
| *dough_age* | wholemeal_spelt | -0.0132 | -0.0128 | 0.0403 | 0.7564 | 0.8883 | 0.7632 | 0.8992 |
| *dough_age* | wholemeal_wheat | 0.0835 | 0.0247 | 0.2188 | 0.0492 | 0.2404 | 0.5613 | 0.7823 |
| *dough_age* | tta_laboratory | 0.0448 | 0.1080 | 0.4586 | 0.2984 | 0.6404 | 0.0120 | 0.0671 |
| *dough_yield* | bread_density | 0.0089 | 0.1312 | 0.5179 | 0.8524 | 0.9243 | 0.0056 | 0.0364 |
| *dough_yield* | dough_age | 0.0184 | 0.0476 | 0.3702 | 0.6651 | 0.8633 | 0.2636 | 0.5410 |
| *dough_yield* | pH_laboratory | 0.0395 | 0.0192 | 0.9833 | 0.3582 | 0.6739 | 0.6550 | 0.8388 |
| *dough_yield* | backslop_freq | -0.0498 | -0.0745 | 0.6483 | 0.2432 | 0.5916 | 0.0804 | 0.2646 |
| *dough_yield* | barley | 0.0372 | 0.0615 | 0.0529 | 0.3817 | 0.6970 | 0.1481 | 0.3872 |
| *dough_yield* | bread_ph | -0.0983 | -0.1069 | 0.9729 | 0.0388 | 0.2069 | 0.0246 | 0.1141 |
| *dough_yield* | dough_size_feeding | -0.0100 | -0.0656 | 0.2370 | 0.8146 | 0.9099 | 0.1225 | 0.3561 |
| *dough_yield* | ph_value_citizen | -0.0323 | -0.1101 | 0.9793 | 0.4526 | 0.7524 | 0.0103 | 0.0591 |
| *dough_yield* | rye | 0.1453 | 0.1883 | 0.4659 | 0.0006 | 0.0064 | 0.0000 | 0.0001 |
| *dough_yield* | semi-wholemeal_wheat | -0.0567 | -0.0444 | 0.2820 | 0.1823 | 0.5269 | 0.2965 | 0.5695 |
| *dough_yield* | spelt | -0.0879 | -0.0912 | 0.2313 | 0.0384 | 0.2069 | 0.0316 | 0.1356 |
| *dough_yield* | temp_feeding | -0.1016 | -0.0629 | 0.9551 | 0.0167 | 0.1070 | 0.1389 | 0.3771 |
| *dough_yield* | temp_storage | -0.0133 | -0.0767 | 0.7949 | 0.7549 | 0.8883 | 0.0708 | 0.2425 |
| *dough_yield* | time_feeding | 0.0960 | 0.2029 | 0.6113 | 0.0242 | 0.1427 | 0.0000 | 0.0000 |
| *dough_yield* | wheat | -0.1986 | -0.2476 | 0.6911 | 0.0000 | 0.0001 | 0.0000 | 0.0000 |
| *dough_yield* | wholemeal_rye | 0.0713 | 0.1441 | 0.4767 | 0.0933 | 0.3697 | 0.0007 | 0.0065 |
| *dough_yield* | wholemeal_spelt | -0.0504 | -0.0813 | 0.1298 | 0.2354 | 0.5843 | 0.0556 | 0.2038 |
| *dough_yield* | wholemeal_wheat | 0.0332 | 0.0741 | 0.3955 | 0.4357 | 0.7512 | 0.0813 | 0.2650 |
| *dough_yield* | tta_laboratory | 0.0564 | 0.1431 | 0.9246 | 0.1910 | 0.5311 | 0.0009 | 0.0075 |
| *elevation* | bread_density | 0.0220 | 0.1449 | 0.4049 | 0.6433 | 0.8526 | 0.0022 | 0.0159 |
| *elevation* | dough_age | -0.0351 | -0.0792 | 0.2629 | 0.4089 | 0.7217 | 0.0621 | 0.2212 |
| *elevation* | dough_yield | -0.1040 | -0.1969 | 0.7382 | 0.0143 | 0.0947 | 0.0000 | 0.0001 |
| *elevation* | pH_laboratory | -0.1163 | -0.1412 | 0.7610 | 0.0066 | 0.0480 | 0.0010 | 0.0079 |

| | | | | | | | | |
|---|---|---|---|---|---|---|---|---|
| *elevation* | backslop_freq | -0.0207 | -0.0183 | 0.4969 | 0.6274 | 0.8501 | 0.6678 | 0.8479 |
| *elevation* | barley | -0.0514 | -0.1171 | 0.0033 | 0.2261 | 0.5721 | 0.0057 | 0.0364 |
| *elevation* | bread_ph | 0.1097 | 0.1146 | 0.7510 | 0.0207 | 0.1255 | 0.0157 | 0.0815 |
| *elevation* | dough_size_feeding | 0.0812 | 0.1148 | 0.2360 | 0.0554 | 0.2477 | 0.0067 | 0.0412 |
| *elevation* | ph_value_citizen | 0.0576 | 0.1121 | 0.7627 | 0.1801 | 0.5269 | 0.0089 | 0.0518 |
| *elevation* | rye | 0.1045 | 0.0932 | 0.4026 | 0.0136 | 0.0917 | 0.0279 | 0.1240 |
| *elevation* | semi-wholemeal_wheat | 0.1618 | 0.1840 | 0.3252 | 0.0001 | 0.0016 | 0.0000 | 0.0002 |
| *elevation* | spelt | 0.2026 | 0.1436 | 0.3173 | 0.0000 | 0.0000 | 0.0007 | 0.0065 |
| *elevation* | temp_feeding | -0.0134 | 0.0430 | 0.7396 | 0.7521 | 0.8878 | 0.3109 | 0.5865 |
| *elevation* | temp_storage | -0.0078 | 0.0663 | 0.6125 | 0.8542 | 0.9247 | 0.1183 | 0.3489 |
| *elevation* | time_feeding | -0.0762 | -0.0600 | 0.4229 | 0.0734 | 0.3105 | 0.1585 | 0.3963 |
| *elevation* | wheat | -0.2388 | -0.2317 | 0.4463 | 0.0000 | 0.0000 | 0.0000 | 0.0000 |
| *elevation* | wholemeal_rye | 0.0397 | 0.0488 | 0.3835 | 0.3500 | 0.6662 | 0.2499 | 0.5373 |
| *elevation* | wholemeal_spelt | 0.0242 | 0.0201 | 0.1225 | 0.5690 | 0.8365 | 0.6352 | 0.8274 |
| *elevation* | wholemeal_wheat | 0.0176 | 0.0330 | 0.3116 | 0.6787 | 0.8637 | 0.4364 | 0.6925 |
| *elevation* | tta_laboratory | 0.0138 | 0.0721 | 0.7215 | 0.7491 | 0.8872 | 0.0938 | 0.2953 |
| *pH_laboratory* | bread_density | 0.0485 | 0.0432 | 0.5254 | 0.3126 | 0.6519 | 0.3692 | 0.6356 |
| *pH_laboratory* | dough_age | 0.0439 | -0.0795 | 0.4767 | 0.3068 | 0.6426 | 0.0642 | 0.2257 |
| *pH_laboratory* | dough_yield | 0.0395 | 0.0192 | 0.9833 | 0.3582 | 0.6739 | 0.6550 | 0.8388 |
| *pH_laboratory* | backslop_freq | -0.0826 | -0.0999 | 0.6594 | 0.0550 | 0.2477 | 0.0201 | 0.1013 |
| *pH_laboratory* | barley | -0.0076 | -0.0264 | 0.0482 | 0.8593 | 0.9285 | 0.5388 | 0.7645 |
| *pH_laboratory* | bread_ph | -0.0712 | -0.0565 | 0.9897 | 0.1395 | 0.4802 | 0.2413 | 0.5279 |
| *pH_laboratory* | dough_size_feeding | -0.0266 | 0.0501 | 0.2405 | 0.5351 | 0.8188 | 0.2433 | 0.5284 |
| *pH_laboratory* | ph_value_citizen | 0.0694 | 0.0182 | 0.9948 | 0.1102 | 0.4179 | 0.6764 | 0.8520 |
| *pH_laboratory* | rye | -0.0200 | -0.0530 | 0.4474 | 0.6418 | 0.8526 | 0.2168 | 0.4878 |
| *pH_laboratory* | semi-wholemeal_wheat | -0.0954 | -0.0853 | 0.2912 | 0.0261 | 0.1510 | 0.0467 | 0.1827 |

| | | | | | | | | |
|---|---|---|---|---|---|---|---|---|
| *pH_laboratory* | spelt | 0.0577 | 0.0104 | 0.2551 | 0.1789 | 0.5269 | 0.8089 | 0.9132 |
| *pH_laboratory* | temp_feeding | -0.0457 | -0.0814 | 0.9722 | 0.2875 | 0.6348 | 0.0579 | 0.2095 |
| *pH_laboratory* | temp_storage | -0.0762 | -0.1100 | 0.8063 | 0.0759 | 0.3148 | 0.0102 | 0.0591 |
| *pH_laboratory* | time_feeding | -0.0039 | -0.0967 | 0.6058 | 0.9285 | 0.9643 | 0.0246 | 0.1141 |
| *pH_laboratory* | wheat | -0.0554 | -0.0691 | 0.7215 | 0.1970 | 0.5328 | 0.1073 | 0.3298 |
| *pH_laboratory* | wholemeal_rye | 0.0574 | 0.0762 | 0.4797 | 0.1813 | 0.5269 | 0.0756 | 0.2535 |
| *pH_laboratory* | wholemeal_spelt | 0.0509 | 0.0793 | 0.1445 | 0.2356 | 0.5843 | 0.0645 | 0.2257 |
| *pH_laboratory* | wholemeal_whe at | -0.0416 | -0.0423 | 0.3906 | 0.3324 | 0.6586 | 0.3251 | 0.5989 |
| *pH_laboratory* | tta_laboratory | -0.1745 | -0.1448 | 0.9321 | 0.0000 | 0.0007 | 0.0007 | 0.0067 |
| *backslop_freq* | bread_density | -0.0691 | -0.2610 | 0.3081 | 0.1463 | 0.4802 | 0.0000 | 0.0000 |
| *backslop_freq* | dough_age | 0.0759 | 0.0383 | 0.2983 | 0.0745 | 0.3116 | 0.3688 | 0.6356 |
| *backslop_freq* | dough_yield | -0.0498 | -0.0745 | 0.6483 | 0.2432 | 0.5916 | 0.0804 | 0.2646 |
| *backslop_freq* | pH_laboratory | -0.0826 | -0.0999 | 0.6594 | 0.0550 | 0.2477 | 0.0201 | 0.1013 |
| *backslop_freq* | barley | -0.0289 | -0.0572 | 0.0104 | 0.4971 | 0.7969 | 0.1791 | 0.4258 |
| *backslop_freq* | bread_ph | -0.0318 | 0.0003 | 0.6664 | 0.5047 | 0.7969 | 0.9955 | 0.9976 |
| *backslop_freq* | dough_size_fee ding | 0.2458 | 0.0971 | 0.3383 | 0.0000 | 0.0000 | 0.0222 | 0.1068 |
| *backslop_freq* | ph_value_citizen | 0.0418 | 0.0635 | 0.6633 | 0.3324 | 0.6586 | 0.1400 | 0.3771 |
| *backslop_freq* | rye | -0.1377 | -0.1170 | 0.2032 | 0.0012 | 0.0112 | 0.0058 | 0.0367 |
| *backslop_freq* | semi-wholemeal_whe at | 0.0414 | 0.0758 | 0.2264 | 0.3308 | 0.6586 | 0.0747 | 0.2523 |
| *backslop_freq* | spelt | -0.0895 | -0.0657 | 0.1015 | 0.0351 | 0.1940 | 0.1225 | 0.3561 |
| *backslop_freq* | temp_feeding | 0.0152 | -0.0166 | 0.6497 | 0.7216 | 0.8844 | 0.6966 | 0.8554 |
| *backslop_freq* | temp_storage | 0.5154 | 0.1991 | 0.7639 | 0.0000 | 0.0000 | 0.0000 | 0.0000 |
| *backslop_freq* | time_feeding | -0.0402 | -0.0501 | 0.3796 | 0.3472 | 0.6661 | 0.2407 | 0.5279 |
| *backslop_freq* | wheat | 0.1424 | 0.0957 | 0.5566 | 0.0008 | 0.0081 | 0.0243 | 0.1141 |
| *backslop_freq* | wholemeal_rye | -0.0941 | -0.0634 | 0.2525 | 0.0268 | 0.1519 | 0.1358 | 0.3769 |
| *backslop_freq* | wholemeal_spelt | 0.0095 | 0.0298 | 0.1007 | 0.8230 | 0.9161 | 0.4845 | 0.7333 |
| *backslop_freq* | wholemeal_whe at | 0.0495 | 0.0745 | 0.2975 | 0.2448 | 0.5931 | 0.0796 | 0.2646 |

| | | | | | | | | |
|---|---|---|---|---|---|---|---|---|
| *backslop_freq* | tta_laboratory | -0.0955 | -0.0701 | 0.5955 | 0.0267 | 0.1519 | 0.1046 | 0.3240 |
| *barley* | bread_density | -0.0182 | -0.0730 | 0.0180 | 0.7020 | 0.8720 | 0.1238 | 0.3561 |
| *barley* | dough_age | -0.0188 | -0.0867 | 0.0005 | 0.6583 | 0.8587 | 0.0410 | 0.1668 |
| *barley* | dough_yield | 0.0372 | 0.0615 | 0.0529 | 0.3817 | 0.6970 | 0.1481 | 0.3872 |
| *barley* | pH_laboratory | -0.0076 | -0.0264 | 0.0482 | 0.8593 | 0.9285 | 0.5388 | 0.7645 |
| *barley* | backslop_freq | -0.0289 | -0.0572 | 0.0104 | 0.4971 | 0.7969 | 0.1791 | 0.4258 |
| *barley* | bread_ph | -0.0758 | -0.0534 | 0.0534 | 0.1109 | 0.4179 | 0.2617 | 0.5410 |
| *barley* | dough_size_fee ding | -0.0068 | 0.0227 | 0.0051 | 0.8730 | 0.9322 | 0.5932 | 0.7929 |
| *barley* | ph_value_citizen | -0.1507 | -0.0961 | 0.0342 | 0.0004 | 0.0047 | 0.0250 | 0.1149 |
| *barley* | rye | -0.0239 | 0.0278 | 0.0003 | 0.5742 | 0.8365 | 0.5121 | 0.7492 |
| *barley* | semi-wholemeal_whe at | -0.0034 | 0.0420 | 0.0109 | 0.9354 | 0.9690 | 0.3228 | 0.5969 |
| *barley* | spelt | -0.0084 | 0.1197 | 0.0039 | 0.8431 | 0.9203 | 0.0047 | 0.0317 |
| *barley* | temp_feeding | 0.0650 | 0.0426 | 0.0614 | 0.1252 | 0.4489 | 0.3158 | 0.5922 |
| *barley* | temp_storage | -0.0221 | -0.0248 | 0.0259 | 0.6026 | 0.8397 | 0.5588 | 0.7823 |
| *barley* | time_feeding | 0.1939 | 0.0492 | 0.1830 | 0.0000 | 0.0001 | 0.2478 | 0.5347 |
| *barley* | wheat | -0.0477 | -0.0205 | 0.0022 | 0.2614 | 0.6076 | 0.6287 | 0.8235 |
| *barley* | wholemeal_rye | -0.0259 | -0.0478 | 0.0000 | 0.5419 | 0.8229 | 0.2599 | 0.5410 |
| *barley* | wholemeal_spelt | -0.0068 | -0.0121 | 0.0000 | 0.8723 | 0.9322 | 0.7762 | 0.9073 |
| *barley* | wholemeal_whe at | -0.0207 | -0.0398 | 0.0000 | 0.6256 | 0.8494 | 0.3483 | 0.6129 |
| *barley* | tta_laboratory | 0.2350 | 0.0278 | 0.1281 | 0.0000 | 0.0000 | 0.5189 | 0.7550 |
| *bread_ph* | bread_density | -0.0023 | 0.0266 | 0.5177 | 0.9614 | 0.9706 | 0.5787 | 0.7897 |
| *bread_ph* | dough_age | 0.0097 | -0.0192 | 0.3755 | 0.8389 | 0.9203 | 0.6875 | 0.8554 |
| *bread_ph* | dough_yield | -0.0983 | -0.1069 | 0.9729 | 0.0388 | 0.2069 | 0.0246 | 0.1141 |
| *bread_ph* | pH_laboratory | -0.0712 | -0.0565 | 0.9897 | 0.1395 | 0.4802 | 0.2413 | 0.5279 |
| *bread_ph* | backslop_freq | -0.0318 | 0.0003 | 0.6664 | 0.5047 | 0.7969 | 0.9955 | 0.9976 |
| *bread_ph* | barley | -0.0758 | -0.0534 | 0.0534 | 0.1109 | 0.4179 | 0.2617 | 0.5410 |
| *bread_ph* | dough_size_fee ding | -0.0211 | 0.0489 | 0.2134 | 0.6582 | 0.8587 | 0.3042 | 0.5789 |

| | | | | | | | | |
|---|---|---|---|---|---|---|---|---|
| *bread_ph* | ph_value_citizen | 0.2152 | 0.2655 | 0.9898 | 0.0000 | 0.0001 | 0.0000 | 0.0000 |
| *bread_ph* | rye | -0.0119 | -0.0268 | 0.4460 | 0.8025 | 0.9062 | 0.5730 | 0.7897 |
| *bread_ph* | semi-wholemeal_wheat | -0.0106 | 0.0132 | 0.2679 | 0.8231 | 0.9161 | 0.7816 | 0.9080 |
| *bread_ph* | spelt | -0.0560 | -0.0053 | 0.2396 | 0.2392 | 0.5879 | 0.9106 | 0.9730 |
| *bread_ph* | temp_feeding | -0.0523 | -0.1461 | 0.9685 | 0.2716 | 0.6177 | 0.0020 | 0.0152 |
| *bread_ph* | temp_storage | -0.0584 | -0.0284 | 0.8023 | 0.2196 | 0.5628 | 0.5512 | 0.7771 |
| *bread_ph* | time_feeding | -0.0239 | -0.0409 | 0.6411 | 0.6171 | 0.8397 | 0.3921 | 0.6500 |
| *bread_ph* | wheat | -0.0887 | -0.1046 | 0.7152 | 0.0617 | 0.2701 | 0.0276 | 0.1240 |
| *bread_ph* | wholemeal_rye | 0.0248 | 0.0755 | 0.4779 | 0.6018 | 0.8397 | 0.1121 | 0.3394 |
| *bread_ph* | wholemeal_spelt | 0.0294 | 0.0102 | 0.1450 | 0.5362 | 0.8188 | 0.8306 | 0.9256 |
| *bread_ph* | wholemeal_wheat | 0.0226 | 0.0136 | 0.3911 | 0.6348 | 0.8526 | 0.7751 | 0.9073 |
| *bread_ph* | tta_laboratory | -0.0060 | 0.0015 | 0.9350 | 0.9012 | 0.9526 | 0.9745 | 0.9934 |
| *dough_size_feeding* | bread_density | -0.0194 | -0.0203 | 0.1040 | 0.6833 | 0.8657 | 0.6689 | 0.8479 |
| *dough_size_feeding* | dough_age | 0.0364 | -0.0128 | 0.1234 | 0.3915 | 0.7047 | 0.7631 | 0.8992 |
| *dough_size_feeding* | dough_yield | -0.0100 | -0.0656 | 0.2370 | 0.8146 | 0.9099 | 0.1225 | 0.3561 |
| *dough_size_feeding* | pH_laboratory | -0.0266 | 0.0501 | 0.2405 | 0.5351 | 0.8188 | 0.2433 | 0.5284 |
| *dough_size_feeding* | backslop_freq | 0.2458 | 0.0971 | 0.3383 | 0.0000 | 0.0000 | 0.0222 | 0.1068 |
| *dough_size_feeding* | barley | -0.0068 | 0.0227 | 0.0051 | 0.8730 | 0.9322 | 0.5932 | 0.7929 |
| *dough_size_feeding* | bread_ph | -0.0211 | 0.0489 | 0.2134 | 0.6582 | 0.8587 | 0.3042 | 0.5789 |
| *dough_size_feeding* | ph_value_citizen | -0.0179 | 0.0721 | 0.2364 | 0.6764 | 0.8637 | 0.0931 | 0.2947 |
| *dough_size_feeding* | rye | -0.0322 | -0.0642 | 0.0813 | 0.4482 | 0.7524 | 0.1304 | 0.3689 |
| *dough_size_feeding* | semi-wholemeal_wheat | -0.0071 | 0.0103 | 0.0653 | 0.8675 | 0.9322 | 0.8087 | 0.9132 |
| *dough_size_feeding* | spelt | 0.0415 | -0.0180 | 0.0997 | 0.3278 | 0.6586 | 0.6723 | 0.8491 |
| *dough_size_feeding* | temp_feeding | 0.0275 | -0.0342 | 0.2428 | 0.5167 | 0.8057 | 0.4200 | 0.6802 |

| | | | | | | | | |
|---|---|---|---|---|---|---|---|---|
| *dough_size_feeding* | temp_storage | 0.1935 | 0.0494 | 0.3069 | 0.0000 | 0.0001 | 0.2446 | 0.5295 |
| *dough_size_feeding* | time_feeding | -0.0354 | -0.1166 | 0.1206 | 0.4060 | 0.7217 | 0.0061 | 0.0378 |
| *dough_size_feeding* | wheat | 0.0489 | 0.0006 | 0.2091 | 0.2497 | 0.6005 | 0.9879 | 0.9976 |
| *dough_size_feeding* | wholemeal_rye | -0.0201 | -0.0417 | 0.0980 | 0.6356 | 0.8526 | 0.3263 | 0.5994 |
| *dough_size_feeding* | wholemeal_spelt | 0.0015 | -0.0118 | 0.0356 | 0.9714 | 0.9790 | 0.7815 | 0.9080 |
| *dough_size_feeding* | wholemeal_wheat | -0.0366 | 0.0548 | 0.0635 | 0.3888 | 0.7021 | 0.1963 | 0.4531 |
| *dough_size_feeding* | tta_laboratory | -0.0233 | -0.0348 | 0.2185 | 0.5894 | 0.8397 | 0.4188 | 0.6800 |
| *ph_value_citizen* | bread_density | -0.0049 | -0.0888 | 0.5202 | 0.9181 | 0.9578 | 0.0621 | 0.2212 |
| *ph_value_citizen* | dough_age | -0.0419 | 0.0137 | 0.3621 | 0.3294 | 0.6586 | 0.7503 | 0.8902 |
| *ph_value_citizen* | dough_yield | -0.0323 | -0.1101 | 0.9793 | 0.4526 | 0.7524 | 0.0103 | 0.0591 |
| *ph_value_citizen* | pH_laboratory | 0.0694 | 0.0182 | 0.9948 | 0.1102 | 0.4179 | 0.6764 | 0.8520 |
| *ph_value_citizen* | backslop_freq | 0.0418 | 0.0635 | 0.6633 | 0.3324 | 0.6586 | 0.1400 | 0.3771 |
| *ph_value_citizen* | barley | -0.1507 | -0.0961 | 0.0342 | 0.0004 | 0.0047 | 0.0250 | 0.1149 |
| *ph_value_citizen* | bread_ph | 0.2152 | 0.2655 | 0.9898 | 0.0000 | 0.0001 | 0.0000 | 0.0000 |
| *ph_value_citizen* | dough_size_feeding | -0.0179 | 0.0721 | 0.2364 | 0.6764 | 0.8637 | 0.0931 | 0.2947 |
| *ph_value_citizen* | rye | -0.1535 | -0.1693 | 0.4322 | 0.0003 | 0.0038 | 0.0001 | 0.0009 |
| *ph_value_citizen* | semi-wholemeal_wheat | 0.0164 | 0.0281 | 0.2992 | 0.7036 | 0.8720 | 0.5126 | 0.7492 |
| *ph_value_citizen* | spelt | -0.0406 | -0.0256 | 0.2449 | 0.3449 | 0.6661 | 0.5514 | 0.7771 |
| *ph_value_citizen* | temp_feeding | 0.0279 | -0.0026 | 0.9714 | 0.5157 | 0.8057 | 0.9523 | 0.9870 |
| *ph_value_citizen* | temp_storage | -0.0044 | 0.0069 | 0.8042 | 0.9182 | 0.9578 | 0.8732 | 0.9534 |
| *ph_value_citizen* | time_feeding | -0.2002 | -0.2445 | 0.6385 | 0.0000 | 0.0001 | 0.0000 | 0.0000 |
| *ph_value_citizen* | wheat | 0.1057 | 0.1155 | 0.7260 | 0.0137 | 0.0917 | 0.0070 | 0.0424 |
| *ph_value_citizen* | wholemeal_rye | -0.0241 | -0.0375 | 0.4722 | 0.5749 | 0.8365 | 0.3829 | 0.6485 |
| *ph_value_citizen* | wholemeal_spelt | 0.0192 | 0.0314 | 0.1434 | 0.6546 | 0.8573 | 0.4646 | 0.7157 |
| *ph_value_citizen* | wholemeal_wheat | 0.0245 | 0.0105 | 0.4006 | 0.5693 | 0.8365 | 0.8076 | 0.9132 |
| *ph_value_citizen* | tta_laboratory | -0.3153 | -0.2419 | 0.9221 | 0.0000 | 0.0000 | 0.0000 | 0.0000 |

| | | | | | | | | |
|---|---|---|---|---|---|---|---|---|
| *rye* | bread_density | 0.0284 | 0.1110 | 0.2604 | 0.5495 | 0.8251 | 0.0190 | 0.0964 |
| *rye* | dough_age | -0.0164 | 0.0045 | 0.1541 | 0.6989 | 0.8720 | 0.9160 | 0.9730 |
| *rye* | dough_yield | 0.1453 | 0.1883 | 0.4659 | 0.0006 | 0.0064 | 0.0000 | 0.0001 |
| *rye* | pH_laboratory | -0.0200 | -0.0530 | 0.4474 | 0.6418 | 0.8526 | 0.2168 | 0.4878 |
| *rye* | backslop_freq | -0.1377 | -0.1170 | 0.2032 | 0.0012 | 0.0112 | 0.0058 | 0.0367 |
| *rye* | barley | -0.0239 | 0.0278 | 0.0003 | 0.5742 | 0.8365 | 0.5121 | 0.7492 |
| *rye* | bread_ph | -0.0119 | -0.0268 | 0.4460 | 0.8025 | 0.9062 | 0.5730 | 0.7897 |
| *rye* | dough_size_feeding | -0.0322 | -0.0642 | 0.0813 | 0.4482 | 0.7524 | 0.1304 | 0.3689 |
| *rye* | ph_value_citizen | -0.1535 | -0.1693 | 0.4322 | 0.0003 | 0.0038 | 0.0001 | 0.0009 |
| *rye* | semi-wholemeal_wheat | -0.0482 | -0.0285 | 0.0917 | 0.2558 | 0.6021 | 0.5014 | 0.7397 |
| *rye* | spelt | -0.0616 | 0.0199 | 0.0589 | 0.1463 | 0.4802 | 0.6387 | 0.8274 |
| *rye* | temp_feeding | -0.0741 | -0.0475 | 0.4230 | 0.0807 | 0.3323 | 0.2634 | 0.5410 |
| *rye* | temp_storage | -0.0226 | 0.0629 | 0.3516 | 0.5944 | 0.8397 | 0.1384 | 0.3771 |
| *rye* | time_feeding | 0.1077 | 0.1831 | 0.3478 | 0.0113 | 0.0780 | 0.0000 | 0.0002 |
| *rye* | wheat | -0.3320 | -0.2938 | 0.1226 | 0.0000 | 0.0000 | 0.0000 | 0.0000 |
| *rye* | wholemeal_rye | -0.0813 | -0.1055 | 0.1491 | 0.0552 | 0.2477 | 0.0127 | 0.0695 |
| *rye* | wholemeal_spelt | -0.0598 | -0.0551 | 0.0102 | 0.1584 | 0.5036 | 0.1944 | 0.4520 |
| *rye* | wholemeal_wheat | -0.1673 | -0.1136 | 0.0405 | 0.0001 | 0.0010 | 0.0073 | 0.0438 |
| *rye* | tta_laboratory | 0.1384 | 0.1556 | 0.4629 | 0.0012 | 0.0119 | 0.0003 | 0.0030 |
| *semi-wholemeal_wheat* | bread_density | -0.0659 | -0.1419 | 0.0883 | 0.1646 | 0.5159 | 0.0027 | 0.0194 |
| *semi-wholemeal_wheat* | dough_age | 0.0136 | -0.0630 | 0.1223 | 0.7492 | 0.8872 | 0.1377 | 0.3771 |
| *semi-wholemeal_wheat* | dough_yield | -0.0567 | -0.0444 | 0.2820 | 0.1823 | 0.5269 | 0.2965 | 0.5695 |
| *semi-wholemeal_wheat* | pH_laboratory | -0.0954 | -0.0853 | 0.2912 | 0.0261 | 0.1510 | 0.0467 | 0.1827 |

| | | | | | | | | |
|---|---|---|---|---|---|---|---|---|
| *semi-wholemeal_wheat* | backslop_freq | 0.0414 | 0.0758 | 0.2264 | 0.3308 | 0.6586 | 0.0747 | 0.2523 |
| *semi-wholemeal_wheat* | barley | -0.0034 | 0.0420 | 0.0109 | 0.9354 | 0.9690 | 0.3228 | 0.5969 |
| *semi-wholemeal_wheat* | bread_ph | -0.0106 | 0.0132 | 0.2679 | 0.8231 | 0.9161 | 0.7816 | 0.9080 |
| *semi-wholemeal_wheat* | dough_size_feeding | -0.0071 | 0.0103 | 0.0653 | 0.8675 | 0.9322 | 0.8087 | 0.9132 |
| *semi-wholemeal_wheat* | ph_value_citizen | 0.0164 | 0.0281 | 0.2992 | 0.7036 | 0.8720 | 0.5126 | 0.7492 |
| *semi-wholemeal_wheat* | rye | -0.0482 | -0.0285 | 0.0917 | 0.2558 | 0.6021 | 0.5014 | 0.7397 |
| *semi-wholemeal_wheat* | spelt | -0.0280 | 0.0289 | 0.0479 | 0.5092 | 0.8020 | 0.4954 | 0.7383 |
| *semi-wholemeal_wheat* | temp_feeding | 0.0542 | 0.0576 | 0.2996 | 0.2014 | 0.5399 | 0.1744 | 0.4193 |
| *semi-wholemeal_wheat* | temp_storage | 0.0254 | 0.0622 | 0.2533 | 0.5495 | 0.8251 | 0.1424 | 0.3794 |
| *semi-wholemeal_wheat* | time_feeding | -0.0254 | -0.0161 | 0.1595 | 0.5504 | 0.8251 | 0.7049 | 0.8577 |
| *semi-wholemeal_wheat* | wheat | -0.2035 | -0.2100 | 0.0810 | 0.0000 | 0.0000 | 0.0000 | 0.0000 |
| *semi-wholemeal_wheat* | wholemeal_rye | -0.0629 | -0.0412 | 0.0872 | 0.1384 | 0.4789 | 0.3312 | 0.6010 |
| *semi-wholemeal_wheat* | wholemeal_spelt | -0.0439 | -0.0460 | 0.0000 | 0.3009 | 0.6404 | 0.2787 | 0.5522 |
| *semi-wholemeal_wheat* | wholemeal_wheat | 0.1408 | 0.1730 | 0.2404 | 0.0009 | 0.0089 | 0.0000 | 0.0006 |
| *semi-wholemeal_wheat* | tta_laboratory | 0.0322 | 0.0643 | 0.2887 | 0.4554 | 0.7550 | 0.1352 | 0.3769 |
| *spelt* | bread_density | 0.0349 | 0.1040 | 0.1612 | 0.4626 | 0.7609 | 0.0281 | 0.1240 |
| *spelt* | dough_age | -0.0544 | -0.1531 | 0.0443 | 0.2001 | 0.5386 | 0.0003 | 0.0031 |

| | | | | | | | | |
|---|---|---|---|---|---|---|---|---|
| *spelt* | dough_yield | -0.0879 | -0.0912 | 0.2313 | 0.0384 | 0.2069 | 0.0316 | 0.1356 |
| *spelt* | pH_laboratory | 0.0577 | 0.0104 | 0.2551 | 0.1789 | 0.5269 | 0.8089 | 0.9132 |
| *spelt* | backslop_freq | -0.0895 | -0.0657 | 0.1015 | 0.0351 | 0.1940 | 0.1225 | 0.3561 |
| *spelt* | barley | -0.0084 | 0.1197 | 0.0039 | 0.8431 | 0.9203 | 0.0047 | 0.0317 |
| *spelt* | bread_ph | -0.0560 | -0.0053 | 0.2396 | 0.2392 | 0.5879 | 0.9106 | 0.9730 |
| *spelt* | dough_size_feeding | 0.0415 | -0.0180 | 0.0997 | 0.3278 | 0.6586 | 0.6723 | 0.8491 |
| *spelt* | ph_value_citizen | -0.0406 | -0.0256 | 0.2449 | 0.3449 | 0.6661 | 0.5514 | 0.7771 |
| *spelt* | rye | -0.0616 | 0.0199 | 0.0589 | 0.1463 | 0.4802 | 0.6387 | 0.8274 |
| *spelt* | semi-wholemeal_wheat | -0.0280 | 0.0289 | 0.0479 | 0.5092 | 0.8020 | 0.4954 | 0.7383 |
| *spelt* | temp_feeding | -0.0360 | -0.0449 | 0.2354 | 0.3968 | 0.7101 | 0.2898 | 0.5634 |
| *spelt* | temp_storage | -0.0675 | 0.0412 | 0.1635 | 0.1114 | 0.4179 | 0.3320 | 0.6010 |
| *spelt* | time_feeding | 0.0557 | 0.0106 | 0.1953 | 0.1912 | 0.5311 | 0.8042 | 0.9132 |
| *spelt* | wheat | -0.2059 | -0.1767 | 0.0443 | 0.0000 | 0.0000 | 0.0000 | 0.0004 |
| *spelt* | wholemeal_rye | -0.0982 | -0.0966 | 0.0346 | 0.0204 | 0.1255 | 0.0226 | 0.1079 |
| *spelt* | wholemeal_spelt | 0.0843 | 0.0561 | 0.1159 | 0.0467 | 0.2337 | 0.1859 | 0.4358 |
| *spelt* | wholemeal_wheat | -0.0550 | -0.0281 | 0.0498 | 0.1947 | 0.5311 | 0.5080 | 0.7461 |
| *spelt* | tta_laboratory | 0.0289 | 0.0370 | 0.2470 | 0.5019 | 0.7969 | 0.3903 | 0.6500 |
| *temp_feeding* | bread_density | 0.0117 | -0.0266 | 0.5155 | 0.8054 | 0.9062 | 0.5754 | 0.7897 |
| *temp_feeding* | dough_age | 0.0198 | 0.0162 | 0.3674 | 0.6418 | 0.8526 | 0.7023 | 0.8577 |
| *temp_feeding* | dough_yield | -0.1016 | -0.0629 | 0.9551 | 0.0167 | 0.1070 | 0.1389 | 0.3771 |
| *temp_feeding* | pH_laboratory | -0.0457 | -0.0814 | 0.9722 | 0.2875 | 0.6348 | 0.0579 | 0.2095 |
| *temp_feeding* | backslop_freq | 0.0152 | -0.0166 | 0.6497 | 0.7216 | 0.8844 | 0.6966 | 0.8554 |
| *temp_feeding* | barley | 0.0650 | 0.0426 | 0.0614 | 0.1252 | 0.4489 | 0.3158 | 0.5922 |
| *temp_feeding* | bread_ph | -0.0523 | -0.1461 | 0.9685 | 0.2716 | 0.6177 | 0.0020 | 0.0152 |
| *temp_feeding* | dough_size_feeding | 0.0275 | -0.0342 | 0.2428 | 0.5167 | 0.8057 | 0.4200 | 0.6802 |
| *temp_feeding* | ph_value_citizen | 0.0279 | -0.0026 | 0.9714 | 0.5157 | 0.8057 | 0.9523 | 0.9870 |
| *temp_feeding* | rye | -0.0741 | -0.0475 | 0.4230 | 0.0807 | 0.3323 | 0.2634 | 0.5410 |

| | | | | | | | | |
|---|---|---|---|---|---|---|---|---|
| *temp_feeding* | semi-wholemeal_wheat | 0.0542 | 0.0576 | 0.2996 | 0.2014 | 0.5399 | 0.1744 | 0.4193 |
| *temp_feeding* | spelt | -0.0360 | -0.0449 | 0.2354 | 0.3968 | 0.7101 | 0.2898 | 0.5634 |
| *temp_feeding* | temp_storage | 0.0590 | 0.0221 | 0.7957 | 0.1644 | 0.5159 | 0.6030 | 0.8003 |
| *temp_feeding* | time_feeding | -0.3444 | -0.3289 | 0.5340 | 0.0000 | 0.0000 | 0.0000 | 0.0000 |
| *temp_feeding* | wheat | 0.0552 | 0.0410 | 0.7161 | 0.1932 | 0.5311 | 0.3342 | 0.6032 |
| *temp_feeding* | wholemeal_rye | 0.0453 | 0.0290 | 0.4708 | 0.2863 | 0.6348 | 0.4942 | 0.7383 |
| *temp_feeding* | wholemeal_spelt | -0.0499 | -0.0005 | 0.1258 | 0.2398 | 0.5879 | 0.9909 | 0.9976 |
| *temp_feeding* | wholemeal_wheat | 0.0022 | -0.0402 | 0.3859 | 0.9588 | 0.9706 | 0.3441 | 0.6107 |
| *temp_feeding* | tta_laboratory | 0.1179 | 0.1050 | 0.9211 | 0.0060 | 0.0448 | 0.0146 | 0.0772 |
| *temp_storage* | bread_density | 0.0654 | 0.0941 | 0.4589 | 0.1679 | 0.5237 | 0.0470 | 0.1828 |
| *temp_storage* | dough_age | 0.0701 | 0.0368 | 0.3398 | 0.0989 | 0.3869 | 0.3871 | 0.6498 |
| *temp_storage* | dough_yield | -0.0133 | -0.0767 | 0.7949 | 0.7549 | 0.8883 | 0.0708 | 0.2425 |
| *temp_storage* | pH_laboratory | -0.0762 | -0.1100 | 0.8063 | 0.0759 | 0.3148 | 0.0102 | 0.0591 |
| *temp_storage* | backslop_freq | 0.5154 | 0.1991 | 0.7639 | 0.0000 | 0.0000 | 0.0000 | 0.0000 |
| *temp_storage* | barley | -0.0221 | -0.0248 | 0.0259 | 0.6026 | 0.8397 | 0.5588 | 0.7823 |
| *temp_storage* | bread_ph | -0.0584 | -0.0284 | 0.8023 | 0.2196 | 0.5628 | 0.5512 | 0.7771 |
| *temp_storage* | dough_size_feeding | 0.1935 | 0.0494 | 0.3069 | 0.0000 | 0.0001 | 0.2446 | 0.5295 |
| *temp_storage* | ph_value_citizen | -0.0044 | 0.0069 | 0.8042 | 0.9182 | 0.9578 | 0.8732 | 0.9534 |
| *temp_storage* | rye | -0.0226 | 0.0629 | 0.3516 | 0.5944 | 0.8397 | 0.1384 | 0.3771 |
| *temp_storage* | semi-wholemeal_wheat | 0.0254 | 0.0622 | 0.2533 | 0.5495 | 0.8251 | 0.1424 | 0.3794 |
| *temp_storage* | spelt | -0.0675 | 0.0412 | 0.1635 | 0.1114 | 0.4179 | 0.3320 | 0.6010 |
| *temp_storage* | temp_feeding | 0.0590 | 0.0221 | 0.7957 | 0.1644 | 0.5159 | 0.6030 | 0.8003 |
| *temp_storage* | time_feeding | -0.0031 | -0.0021 | 0.4905 | 0.9428 | 0.9690 | 0.9604 | 0.9881 |
| *temp_storage* | wheat | 0.0829 | -0.0379 | 0.6209 | 0.0505 | 0.2428 | 0.3714 | 0.6376 |
| *temp_storage* | wholemeal_rye | -0.0400 | 0.0706 | 0.3627 | 0.3462 | 0.6661 | 0.0959 | 0.3007 |
| *temp_storage* | wholemeal_spelt | -0.0651 | -0.0872 | 0.0758 | 0.1251 | 0.4489 | 0.0396 | 0.1654 |

| | | | | | | | | |
|---|---|---|---|---|---|---|---|---|
| *temp_storage* | wholemeal_wheat | -0.0313 | -0.1047 | 0.3030 | 0.4608 | 0.7600 | 0.0134 | 0.0728 |
| *temp_storage* | tta_laboratory | -0.0223 | 0.0749 | 0.7530 | 0.6046 | 0.8397 | 0.0816 | 0.2650 |
| *time_feeding* | bread_density | -0.0336 | 0.0861 | 0.2895 | 0.4804 | 0.7795 | 0.0703 | 0.2425 |
| *time_feeding* | dough_age | -0.0002 | 0.0030 | 0.2267 | 0.9965 | 0.9965 | 0.9447 | 0.9837 |
| *time_feeding* | dough_yield | 0.0960 | 0.2029 | 0.6113 | 0.0242 | 0.1427 | 0.0000 | 0.0000 |
| *time_feeding* | pH_laboratory | -0.0039 | -0.0967 | 0.6058 | 0.9285 | 0.9643 | 0.0246 | 0.1141 |
| *time_feeding* | backslop_freq | -0.0402 | -0.0501 | 0.3796 | 0.3472 | 0.6661 | 0.2407 | 0.5279 |
| *time_feeding* | barley | 0.1939 | 0.0492 | 0.1830 | 0.0000 | 0.0001 | 0.2478 | 0.5347 |
| *time_feeding* | bread_ph | -0.0239 | -0.0409 | 0.6411 | 0.6171 | 0.8397 | 0.3921 | 0.6500 |
| *time_feeding* | dough_size_feeding | -0.0354 | -0.1166 | 0.1206 | 0.4060 | 0.7217 | 0.0061 | 0.0378 |
| *time_feeding* | ph_value_citizen | -0.2002 | -0.2445 | 0.6385 | 0.0000 | 0.0001 | 0.0000 | 0.0000 |
| *time_feeding* | rye | 0.1077 | 0.1831 | 0.3478 | 0.0113 | 0.0780 | 0.0000 | 0.0002 |
| *time_feeding* | semi-wholemeal_wheat | -0.0254 | -0.0161 | 0.1595 | 0.5504 | 0.8251 | 0.7049 | 0.8577 |
| *time_feeding* | spelt | 0.0557 | 0.0106 | 0.1953 | 0.1912 | 0.5311 | 0.8042 | 0.9132 |
| *time_feeding* | temp_feeding | -0.3444 | -0.3289 | 0.5340 | 0.0000 | 0.0000 | 0.0000 | 0.0000 |
| *time_feeding* | temp_storage | -0.0031 | -0.0021 | 0.4905 | 0.9428 | 0.9690 | 0.9604 | 0.9881 |
| *time_feeding* | wheat | -0.0863 | -0.1671 | 0.3961 | 0.0426 | 0.2164 | 0.0001 | 0.0010 |
| *time_feeding* | wholemeal_rye | 0.0107 | 0.0606 | 0.2960 | 0.8020 | 0.9062 | 0.1546 | 0.3943 |
| *time_feeding* | wholemeal_spelt | 0.0408 | 0.0362 | 0.1179 | 0.3382 | 0.6658 | 0.3958 | 0.6527 |
| *time_feeding* | wholemeal_wheat | -0.0193 | 0.0516 | 0.2277 | 0.6501 | 0.8568 | 0.2262 | 0.5035 |
| *time_feeding* | tta_laboratory | 0.2309 | 0.1451 | 0.6317 | 0.0000 | 0.0000 | 0.0007 | 0.0067 |
| *wheat* | bread_density | -0.0694 | -0.3064 | 0.3443 | 0.1435 | 0.4802 | 0.0000 | 0.0000 |
| *wheat* | dough_age | 0.0022 | -0.0106 | 0.2720 | 0.9579 | 0.9706 | 0.8032 | 0.9132 |
| *wheat* | dough_yield | -0.1986 | -0.2476 | 0.6911 | 0.0000 | 0.0001 | 0.0000 | 0.0000 |
| *wheat* | pH_laboratory | -0.0554 | -0.0691 | 0.7215 | 0.1970 | 0.5328 | 0.1073 | 0.3298 |
| *wheat* | backslop_freq | 0.1424 | 0.0957 | 0.5566 | 0.0008 | 0.0081 | 0.0243 | 0.1141 |
| *wheat* | barley | -0.0477 | -0.0205 | 0.0022 | 0.2614 | 0.6076 | 0.6287 | 0.8235 |

| | | | | | | | | |
|---|---|---|---|---|---|---|---|---|
| *wheat* | bread_ph | -0.0887 | -0.1046 | 0.7152 | 0.0617 | 0.2701 | 0.0276 | 0.1240 |
| *wheat* | dough_size_fee ding | 0.0489 | 0.0006 | 0.2091 | 0.2497 | 0.6005 | 0.9879 | 0.9976 |
| *wheat* | ph_value_citizen | 0.1057 | 0.1155 | 0.7260 | 0.0137 | 0.0917 | 0.0070 | 0.0424 |
| *wheat* | rye | -0.3320 | -0.2938 | 0.1226 | 0.0000 | 0.0000 | 0.0000 | 0.0000 |
| *wheat* | semi-wholemeal_whe at | -0.2035 | -0.2100 | 0.0810 | 0.0000 | 0.0000 | 0.0000 | 0.0000 |
| *wheat* | spelt | -0.2059 | -0.1767 | 0.0443 | 0.0000 | 0.0000 | 0.0000 | 0.0004 |
| *wheat* | temp_feeding | 0.0552 | 0.0410 | 0.7161 | 0.1932 | 0.5311 | 0.3342 | 0.6032 |
| *wheat* | temp_storage | 0.0829 | -0.0379 | 0.6209 | 0.0505 | 0.2428 | 0.3714 | 0.6376 |
| *wheat* | time_feeding | -0.0863 | -0.1671 | 0.3961 | 0.0426 | 0.2164 | 0.0001 | 0.0010 |
| *wheat* | wholemeal_rye | -0.4041 | -0.3969 | 0.0997 | 0.0000 | 0.0000 | 0.0000 | 0.0000 |
| *wheat* | wholemeal_spelt | -0.1184 | -0.1121 | 0.0215 | 0.0051 | 0.0391 | 0.0081 | 0.0476 |
| *wheat* | wholemeal_whe at | -0.2407 | -0.2216 | 0.1353 | 0.0000 | 0.0000 | 0.0000 | 0.0000 |
| *wheat* | tta_laboratory | -0.5197 | -0.5854 | 0.5514 | 0.0000 | 0.0000 | 0.0000 | 0.0000 |
| *wholemeal_rye* | bread_density | 0.1042 | 0.2153 | 0.3289 | 0.0279 | 0.1567 | 0.0000 | 0.0001 |
| *wholemeal_rye* | dough_age | 0.0214 | 0.0943 | 0.1944 | 0.6151 | 0.8397 | 0.0262 | 0.1188 |
| *wholemeal_rye* | dough_yield | 0.0713 | 0.1441 | 0.4767 | 0.0933 | 0.3697 | 0.0007 | 0.0065 |
| *wholemeal_rye* | pH_laboratory | 0.0574 | 0.0762 | 0.4797 | 0.1813 | 0.5269 | 0.0756 | 0.2535 |
| *wholemeal_rye* | backslop_freq | -0.0941 | -0.0634 | 0.2525 | 0.0268 | 0.1519 | 0.1358 | 0.3769 |
| *wholemeal_rye* | barley | -0.0259 | -0.0478 | 0.0000 | 0.5419 | 0.8229 | 0.2599 | 0.5410 |
| *wholemeal_rye* | bread_ph | 0.0248 | 0.0755 | 0.4779 | 0.6018 | 0.8397 | 0.1121 | 0.3394 |
| *wholemeal_rye* | dough_size_fee ding | -0.0201 | -0.0417 | 0.0980 | 0.6356 | 0.8526 | 0.3263 | 0.5994 |
| *wholemeal_rye* | ph_value_citizen | -0.0241 | -0.0375 | 0.4722 | 0.5749 | 0.8365 | 0.3829 | 0.6485 |
| *wholemeal_rye* | rye | -0.0813 | -0.1055 | 0.1491 | 0.0552 | 0.2477 | 0.0127 | 0.0695 |
| *wholemeal_rye* | semi-wholemeal_whe at | -0.0629 | -0.0412 | 0.0872 | 0.1384 | 0.4789 | 0.3312 | 0.6010 |
| *wholemeal_rye* | spelt | -0.0982 | -0.0966 | 0.0346 | 0.0204 | 0.1255 | 0.0226 | 0.1079 |
| *wholemeal_rye* | temp_feeding | 0.0453 | 0.0290 | 0.4708 | 0.2863 | 0.6348 | 0.4942 | 0.7383 |

| | | | | | | | | |
|---|---|---|---|---|---|---|---|---|
| *wholemeal_rye* | temp_storage | -0.0400 | 0.0706 | 0.3627 | 0.3462 | 0.6661 | 0.0959 | 0.3007 |
| *wholemeal_rye* | time_feeding | 0.0107 | 0.0606 | 0.2960 | 0.8020 | 0.9062 | 0.1546 | 0.3943 |
| *wholemeal_rye* | wheat | -0.4041 | -0.3969 | 0.0997 | 0.0000 | 0.0000 | 0.0000 | 0.0000 |
| *wholemeal_rye* | wholemeal_spelt | -0.0119 | -0.0173 | 0.0562 | 0.7785 | 0.9016 | 0.6845 | 0.8554 |
| *wholemeal_rye* | wholemeal_whe at | -0.0637 | -0.0568 | 0.1360 | 0.1330 | 0.4628 | 0.1803 | 0.4271 |
| *wholemeal_rye* | tta_laboratory | 0.3352 | 0.3856 | 0.5503 | 0.0000 | 0.0000 | 0.0000 | 0.0000 |
| *wholemeal_spelt* | bread_density | -0.0154 | 0.0653 | 0.0616 | 0.7453 | 0.8872 | 0.1684 | 0.4116 |
| *wholemeal_spelt* | dough_age | -0.0132 | -0.0128 | 0.0403 | 0.7564 | 0.8883 | 0.7632 | 0.8992 |
| *wholemeal_spelt* | dough_yield | -0.0504 | -0.0813 | 0.1298 | 0.2354 | 0.5843 | 0.0556 | 0.2038 |
| *wholemeal_spelt* | pH_laboratory | 0.0509 | 0.0793 | 0.1445 | 0.2356 | 0.5843 | 0.0645 | 0.2257 |
| *wholemeal_spelt* | backslop_freq | 0.0095 | 0.0298 | 0.1007 | 0.8230 | 0.9161 | 0.4845 | 0.7333 |
| *wholemeal_spelt* | barley | -0.0068 | -0.0121 | 0.0000 | 0.8723 | 0.9322 | 0.7762 | 0.9073 |
| *wholemeal_spelt* | bread_ph | 0.0294 | 0.0102 | 0.1450 | 0.5362 | 0.8188 | 0.8306 | 0.9256 |
| *wholemeal_spelt* | dough_size_fee ding | 0.0015 | -0.0118 | 0.0356 | 0.9714 | 0.9790 | 0.7815 | 0.9080 |
| *wholemeal_spelt* | ph_value_citizen | 0.0192 | 0.0314 | 0.1434 | 0.6546 | 0.8573 | 0.4646 | 0.7157 |
| *wholemeal_spelt* | rye | -0.0598 | -0.0551 | 0.0102 | 0.1584 | 0.5036 | 0.1944 | 0.4520 |
| *wholemeal_spelt* | semi-wholemeal_whe at | -0.0439 | -0.0460 | 0.0000 | 0.3009 | 0.6404 | 0.2787 | 0.5522 |
| *wholemeal_spelt* | spelt | 0.0843 | 0.0561 | 0.1159 | 0.0467 | 0.2337 | 0.1859 | 0.4358 |
| *wholemeal_spelt* | temp_feeding | -0.0499 | -0.0005 | 0.1258 | 0.2398 | 0.5879 | 0.9909 | 0.9976 |
| *wholemeal_spelt* | temp_storage | -0.0651 | -0.0872 | 0.0758 | 0.1251 | 0.4489 | 0.0396 | 0.1654 |
| *wholemeal_spelt* | time_feeding | 0.0408 | 0.0362 | 0.1179 | 0.3382 | 0.6658 | 0.3958 | 0.6527 |
| *wholemeal_spelt* | wheat | -0.1184 | -0.1121 | 0.0215 | 0.0051 | 0.0391 | 0.0081 | 0.0476 |
| *wholemeal_spelt* | wholemeal_rye | -0.0119 | -0.0173 | 0.0562 | 0.7785 | 0.9016 | 0.6845 | 0.8554 |
| *wholemeal_spelt* | wholemeal_whe at | -0.0009 | -0.0002 | 0.0548 | 0.9837 | 0.9868 | 0.9966 | 0.9976 |
| *wholemeal_spelt* | tta_laboratory | 0.1281 | 0.1395 | 0.1781 | 0.0028 | 0.0232 | 0.0011 | 0.0089 |
| *wholemeal_whe at* | bread_density | -0.0156 | -0.0067 | 0.1915 | 0.7427 | 0.8872 | 0.8881 | 0.9602 |

| | | | | | | | | |
|---|---|---|---|---|---|---|---|---|
| *wholemeal_wheat* | dough_age | 0.0835 | 0.0247 | 0.2188 | 0.0492 | 0.2404 | 0.5613 | 0.7823 |
| *wholemeal_wheat* | dough_yield | 0.0332 | 0.0741 | 0.3955 | 0.4357 | 0.7512 | 0.0813 | 0.2650 |
| *wholemeal_wheat* | pH_laboratory | -0.0416 | -0.0423 | 0.3906 | 0.3324 | 0.6586 | 0.3251 | 0.5989 |
| *wholemeal_wheat* | backslop_freq | 0.0495 | 0.0745 | 0.2975 | 0.2448 | 0.5931 | 0.0796 | 0.2646 |
| *wholemeal_wheat* | barley | -0.0207 | -0.0398 | 0.0000 | 0.6256 | 0.8494 | 0.3483 | 0.6129 |
| *wholemeal_wheat* | bread_ph | 0.0226 | 0.0136 | 0.3911 | 0.6348 | 0.8526 | 0.7751 | 0.9073 |
| *wholemeal_wheat* | dough_size_feeding | -0.0366 | 0.0548 | 0.0635 | 0.3888 | 0.7021 | 0.1963 | 0.4531 |
| *wholemeal_wheat* | ph_value_citizen | 0.0245 | 0.0105 | 0.4006 | 0.5693 | 0.8365 | 0.8076 | 0.9132 |
| *wholemeal_wheat* | rye | -0.1673 | -0.1136 | 0.0405 | 0.0001 | 0.0010 | 0.0073 | 0.0438 |
| *wholemeal_wheat* | semi-wholemeal_wheat | 0.1408 | 0.1730 | 0.2404 | 0.0009 | 0.0089 | 0.0000 | 0.0006 |
| *wholemeal_wheat* | spelt | -0.0550 | -0.0281 | 0.0498 | 0.1947 | 0.5311 | 0.5080 | 0.7461 |
| *wholemeal_wheat* | temp_feeding | 0.0022 | -0.0402 | 0.3859 | 0.9588 | 0.9706 | 0.3441 | 0.6107 |
| *wholemeal_wheat* | temp_storage | -0.0313 | -0.1047 | 0.3030 | 0.4608 | 0.7600 | 0.0134 | 0.0728 |
| *wholemeal_wheat* | time_feeding | -0.0193 | 0.0516 | 0.2277 | 0.6501 | 0.8568 | 0.2262 | 0.5035 |
| *wholemeal_wheat* | wheat | -0.2407 | -0.2216 | 0.1353 | 0.0000 | 0.0000 | 0.0000 | 0.0000 |
| *wholemeal_wheat* | wholemeal_rye | -0.0637 | -0.0568 | 0.1360 | 0.1330 | 0.4628 | 0.1803 | 0.4271 |
| *wholemeal_wheat* | wholemeal_spelt | -0.0009 | -0.0002 | 0.0548 | 0.9837 | 0.9868 | 0.9966 | 0.9976 |
| *wholemeal_wheat* | tta_laboratory | 0.1727 | 0.1917 | 0.4247 | 0.0001 | 0.0008 | 0.0000 | 0.0001 |
| *tta_laboratory* | bread_density | 0.0166 | 0.2313 | 0.5007 | 0.7305 | 0.8872 | 0.0000 | 0.0000 |
| *tta_laboratory* | dough_age | 0.0448 | 0.1080 | 0.4586 | 0.2984 | 0.6404 | 0.0120 | 0.0671 |
| *tta_laboratory* | dough_yield | 0.0564 | 0.1431 | 0.9246 | 0.1910 | 0.5311 | 0.0009 | 0.0075 |
| *tta_laboratory* | pH_laboratory | -0.1745 | -0.1448 | 0.9321 | 0.0000 | 0.0007 | 0.0007 | 0.0067 |
| *tta_laboratory* | backslop_freq | -0.0955 | -0.0701 | 0.5955 | 0.0267 | 0.1519 | 0.1046 | 0.3240 |
| *tta_laboratory* | barley | 0.2350 | 0.0278 | 0.1281 | 0.0000 | 0.0000 | 0.5189 | 0.7550 |

| | | | | | | | | |
|---|---|---|---|---|---|---|---|---|
| *tta_laboratory* | bread_ph | -0.0060 | 0.0015 | 0.9350 | 0.9012 | 0.9526 | 0.9745 | 0.9934 |
| *tta_laboratory* | dough_size_feeding | -0.0233 | -0.0348 | 0.2185 | 0.5894 | 0.8397 | 0.4188 | 0.6800 |
| *tta_laboratory* | ph_value_citizen | -0.3153 | -0.2419 | 0.9221 | 0.0000 | 0.0000 | 0.0000 | 0.0000 |
| *tta_laboratory* | rye | 0.1384 | 0.1556 | 0.4629 | 0.0012 | 0.0119 | 0.0003 | 0.0030 |
| *tta_laboratory* | semi-wholemeal_wheat | 0.0322 | 0.0643 | 0.2887 | 0.4554 | 0.7550 | 0.1352 | 0.3769 |
| *tta_laboratory* | spelt | 0.0289 | 0.0370 | 0.2470 | 0.5019 | 0.7969 | 0.3903 | 0.6500 |
| *tta_laboratory* | temp_feeding | 0.1179 | 0.1050 | 0.9211 | 0.0060 | 0.0448 | 0.0146 | 0.0772 |
| *tta_laboratory* | temp_storage | -0.0223 | 0.0749 | 0.7530 | 0.6046 | 0.8397 | 0.0816 | 0.2650 |
| *tta_laboratory* | time_feeding | 0.2309 | 0.1451 | 0.6317 | 0.0000 | 0.0000 | 0.0007 | 0.0067 |
| *tta_laboratory* | wheat | -0.5197 | -0.5854 | 0.5514 | 0.0000 | 0.0000 | 0.0000 | 0.0000 |
| *tta_laboratory* | wholemeal_rye | 0.3352 | 0.3856 | 0.5503 | 0.0000 | 0.0000 | 0.0000 | 0.0000 |
| *tta_laboratory* | wholemeal_spelt | 0.1281 | 0.1395 | 0.1781 | 0.0028 | 0.0232 | 0.0011 | 0.0089 |
| *tta_laboratory* | wholemeal_wheat | 0.1727 | 0.1917 | 0.4247 | 0.0001 | 0.0008 | 0.0000 | 0.0001 |

**Supplementary Table 8.** Chi-square enrichment test results for sourdough ingredients across countries (related to Figure 7A). *p*-values were corrected using the Benjamini–Hochberg false discovery rate (BH-FDR). Indicator codes: PE = positive enrichment, NE = negative enrichment, M = multiple indicators, U = unique indicator.

| ***Feature*** | ***Category*** | ***Cluster*** | ***Chi2 Statistic*** | ***Cramér's V*** | ***Cluster Count with Category*** | ***Cluster Total*** | ***Category Total*** | ***Raw p-value*** | ***Corrected p-value*** | ***Enrichment Score*** | ***Interpretation*** | ***Indicator*** |
|---|---|---|---|---|---|---|---|---|---|---|---|---|
| *Flour Additions* | yes | Italy | 21.5315 | 0.1398 | 16 | 81 | 74 | 0.0000 | 0.0000 | 2.9416 | PE | M |
| *Flour Change* | yes | France | 7.8217 | 0.0842 | 18 | 23 | 521 | 0.0052 | 0.0213 | 1.6553 | PE | M |
| *Flour Change* | yes | Germany | 17.3575 | 0.1255 | 50 | 158 | 521 | 0.0000 | 0.0002 | 0.6694 | NE | M |
| *Flour Milling Grade* | endosperm | Finland | 50.9589 | 0.2136 | 120 | 142 | 630 | 0.0000 | 0.0000 | 1.4983 | PE | M |
| *Flour Milling Grade* | endosperm | Switzerland | 18.6251 | 0.1291 | 187 | 393 | 630 | 0.0000 | 0.0001 | 0.8436 | NE | M |
| *Flour Milling Grade* | endosperm | Italy | 11.8811 | 0.1031 | 61 | 81 | 630 | 0.0006 | 0.0031 | 1.3352 | PE | M |
| *Flour Milling Grade* | endosperm | Belgium | 10.5508 | 0.0972 | 47 | 113 | 630 | 0.0012 | 0.0055 | 0.7374 | NE | M |
| *Flour Milling Grade* | endosperm | Denmark | 6.2889 | 0.0750 | 4 | 17 | 630 | 0.0121 | 0.0428 | 0.4172 | NE | M |

| | | | | | | | | | | | | |
|---|---|---|---|---|---|---|---|---|---|---|---|---|
| *Flour Milling Grade* | mix | Germany | 7.2910 | 0.0808 | 9 | 158 | 141 | 0.0069 | 0.0252 | 0.4513 | NE | M |
| *Flour Milling Grade* | mix | Denmark | 21.8652 | 0.1399 | 9 | 17 | 141 | 0.0000 | 0.0000 | 4.1940 | PE | M |
| *Flour Milling Grade* | mix | Sweden | 10.5949 | 0.0974 | 9 | 25 | 141 | 0.0011 | 0.0055 | 2.8519 | PE | M |
| *Flour Milling Grade* | mix | United Kingdom | 4.8977 | 0.0662 | 5 | 14 | 141 | 0.0269 | 0.0905 | 2.8293 | PE | M |
| *Flour Milling Grade* | other | Belgium | 43.4672 | 0.1973 | 14 | 113 | 29 | 0.0000 | 0.0000 | 4.7720 | PE | M |
| *Flour Milling Grade* | wholemeal | Finland | 37.6523 | 0.1836 | 9 | 142 | 317 | 0.0000 | 0.0000 | 0.2233 | NE | M |
| *Flour Milling Grade* | wholemeal | Switzerland | 16.2077 | 0.1205 | 141 | 393 | 317 | 0.0001 | 0.0004 | 1.2642 | PE | M |
| *Flour Milling Grade* | wholemeal | Germany | 15.4812 | 0.1177 | 66 | 158 | 317 | 0.0001 | 0.0005 | 1.4719 | PE | M |
| *Flour Milling Grade* | wholemeal | Italy | 5.8948 | 0.0726 | 13 | 81 | 317 | 0.0152 | 0.0527 | 0.5655 | NE | M |
| *Flour Organic* | yes | Finland | 16.4732 | 0.1225 | 64 | 142 | 669 | 0.0000 | 0.0003 | 0.7397 | NE | M |
| *Flour Organic* | yes | Italy | 10.7441 | 0.0989 | 35 | 81 | 669 | 0.0010 | 0.0054 | 0.7092 | NE | M |
| *Flour Organic* | yes | Romania | 10.5630 | 0.0981 | 17 | 46 | 669 | 0.0012 | 0.0055 | 0.6066 | NE | M |
| *Flour Organic* | yes | Germany | 19.7720 | 0.1342 | 122 | 158 | 669 | 0.0000 | 0.0001 | 1.2673 | PE | M |
| *Flour Organic* | yes | Austria | 7.3368 | 0.0817 | 23 | 26 | 669 | 0.0068 | 0.0250 | 1.4519 | PE | M |
| *Flour Type (Grain)* | other | Germany | 7.7723 | 0.0848 | 9 | 156 | 141 | 0.0053 | 0.0214 | 0.4423 | NE | M |
| *Flour Type (Grain)* | other | Denmark | 20.7979 | 0.1387 | 9 | 17 | 141 | 0.0000 | 0.0000 | 4.0588 | PE | M |
| *Flour Type (Grain)* | other | Sweden | 9.9093 | 0.0957 | 9 | 25 | 141 | 0.0016 | 0.0076 | 2.7600 | PE | M |
| *Flour Type (Grain)* | other | United Kingdom | 4.5618 | 0.0650 | 5 | 14 | 141 | 0.0327 | 0.1083 | 2.7381 | PE | M |
| *Flour Type (Grain)* | rye | Germany | 49.2316 | 0.2134 | 79 | 156 | 294 | 0.0000 | 0.0000 | 1.8620 | PE | M |
| *Flour Type (Grain)* | rye | Austria | 35.8923 | 0.1822 | 21 | 26 | 294 | 0.0000 | 0.0000 | 2.9698 | PE | M |
| *Flour Type (Grain)* | rye | Finland | 7.6379 | 0.0841 | 24 | 140 | 294 | 0.0057 | 0.0223 | 0.6303 | NE | M |
| *Flour Type (Grain)* | rye | Romania | 6.3317 | 0.0765 | 4 | 43 | 294 | 0.0119 | 0.0425 | 0.3420 | NE | M |
| *Flour Type (Grain)* | spelt | Switzerland | 18.7321 | 0.1316 | 33 | 383 | 51 | 0.0000 | 0.0001 | 1.8263 | PE | U |
| *Flour Type (Grain)* | wheat | Italy | 35.3790 | 0.1809 | 70 | 80 | 595 | 0.0000 | 0.0000 | 1.5897 | PE | M |
| *Flour Type (Grain)* | wheat | Finland | 21.4634 | 0.1409 | 103 | 140 | 595 | 0.0000 | 0.0000 | 1.3367 | PE | M |

| | | | | | | | | | | | | |
|---|---|---|---|---|---|---|---|---|---|---|---|---|
| *Flour Type (Grain)* | wheat | Germany | 19.4771 | 0.1342 | 60 | 156 | 595 | 0.0000 | 0.0001 | 0.6988 | NE | M |
| *Flour Type (Grain)* | wheat | Austria | 12.3632 | 0.1069 | 5 | 26 | 595 | 0.0004 | 0.0025 | 0.3494 | NE | M |
| *Flour Type (Grain)* | wheat | Switzerland | 8.8784 | 0.0906 | 187 | 383 | 595 | 0.0029 | 0.0128 | 0.8871 | NE | M |
| *Flour Type (Grain)* | wheat | Romania | 7.6346 | 0.0840 | 33 | 43 | 595 | 0.0057 | 0.0223 | 1.3943 | PE | M |
| *Water Source* | bottled water | Switzerland | 7.3664 | 0.0818 | 8 | 391 | 49 | 0.0066 | 0.0250 | 0.4601 | NE | M |
| *Water Source* | bottled water | France | 43.8491 | 0.1995 | 8 | 23 | 49 | 0.0000 | 0.0000 | 7.8225 | PE | M |
| *Water Source* | bottled water | Romania | 15.8865 | 0.1201 | 8 | 46 | 49 | 0.0001 | 0.0004 | 3.9113 | PE | M |
| *Water Source* | bottled water | Italy | 14.9246 | 0.1164 | 11 | 81 | 49 | 0.0001 | 0.0007 | 3.0542 | PE | M |
| *Water Source* | spring water | Romania | 47.6393 | 0.2079 | 5 | 46 | 9 | 0.0000 | 0.0000 | 13.3092 | PE | U |
| *Water Source* | tap water | Romania | 43.6804 | 0.1991 | 22 | 46 | 925 | 0.0000 | 0.0000 | 0.5698 | NE | M |
| *Water Source* | tap water | France | 20.0686 | 0.1349 | 11 | 23 | 925 | 0.0000 | 0.0001 | 0.5698 | NE | M |
| *Water Source* | tap water | Switzerland | 7.8132 | 0.0842 | 345 | 391 | 925 | 0.0052 | 0.0213 | 1.0512 | PE | M |
| *Water Source* | tap water | Austria | 3.9483 | 0.0599 | 26 | 26 | 925 | 0.0469 | 0.1488 | 1.1914 | PE | M |
| *Water Source* | tap water | Finland | 14.0503 | 0.1129 | 135 | 142 | 925 | 0.0002 | 0.0010 | 1.1326 | PE | M |
| *Water Source* | tap water | Belgium | 8.2023 | 0.0863 | 75 | 102 | 925 | 0.0042 | 0.0182 | 0.8760 | NE | M |
| *Water Source* | tap water (descaled) | Germany | 9.7766 | 0.0942 | 22 | 158 | 83 | 0.0018 | 0.0080 | 1.8487 | PE | U |
| *Water Source* | tap water (filtered) | Romania | 41.2981 | 0.1936 | 9 | 46 | 32 | 0.0000 | 0.0000 | 6.7378 | PE | M |
| *Water Source* | tap water (filtered) | Belgium | 11.7526 | 0.1033 | 9 | 102 | 32 | 0.0006 | 0.0033 | 3.0386 | PE | M |
| *Water Source* | tap water (filtered) | France | 5.2865 | 0.0693 | 3 | 23 | 32 | 0.0215 | 0.0734 | 4.4918 | PE | M |
| *Yeast Usage Prior* | yes | Finland | 138.6970 | 0.3548 | 26 | 142 | 696 | 0.0000 | 0.0000 | 0.2899 | NE | M |
| *Yeast Usage Prior* | yes | Romania | 26.7130 | 0.1557 | 12 | 46 | 696 | 0.0000 | 0.0000 | 0.4130 | NE | M |
| *Yeast Usage Prior* | yes | Germany | 38.2563 | 0.1863 | 135 | 158 | 696 | 0.0000 | 0.0000 | 1.3528 | PE | M |
| *Yeast Usage Prior* | yes | Switzerland | 32.3141 | 0.1712 | 291 | 391 | 696 | 0.0000 | 0.0000 | 1.1784 | PE | M |
| *Yeast Usage Prior* | yes | Austria | 4.3669 | 0.0629 | 22 | 26 | 696 | 0.0366 | 0.1179 | 1.3397 | PE | M |

**Supplementary Table 9.** Chi-square enrichment test results for sourdough baking motivations and perceived health benefits across countries (related to Figure 7B). *p*-values were corrected using the Benjamini–Hochberg false discovery rate (BH-FDR). Indicator codes: PE = positive enrichment, NE = negative enrichment, M = multiple indicators, U = unique indicator.

| *Feature* | *Category* | *Cluster* | *Chi2 Statistic* | *Cramér's V* | *Cluster Count with Category* | *Cluster Total* | *Category Total* | *Raw p-value* | *Corrected p-value* | *Enrichment Score* | *Interpretation* | *Indicator* |
|---|---|---|---|---|---|---|---|---|---|---|---|---|
| *Baking Motivation: Enjoy Baking* | yes | Belgium | 20.0601 | 0.1340 | 59 | 113 | 791 | 0.0000 | 0.0005 | 0.7373 | NE | M |
| *Baking Motivation: Enjoy Baking* | yes | Germany | 7.6165 | 0.0826 | 127 | 158 | 791 | 0.0058 | 0.0606 | 1.1351 | PE | M |
| *Baking Motivation: Enjoy Baking* | yes | Finland | 5.5681 | 0.0706 | 113 | 142 | 791 | 0.0183 | 0.1338 | 1.1237 | PE | M |
| *Baking Motivation: Health* | yes | Italy | 31.0100 | 0.1666 | 17 | 81 | 574 | 0.0000 | 0.0000 | 0.4084 | NE | M |
| *Baking Motivation: Health* | yes | Romania | 12.7701 | 0.1069 | 36 | 46 | 574 | 0.0004 | 0.0130 | 1.5230 | PE | M |
| *Baking Motivation: Health* | yes | Germany | 6.0409 | 0.0735 | 96 | 158 | 574 | 0.0140 | 0.1114 | 1.1824 | PE | M |
| *Baking Motivation: Intolerance/Allergy* | yes | Finland | 4.0948 | 0.0605 | 5 | 142 | 15 | 0.0430 | 0.2252 | 2.6221 | PE | M |
| *Baking Motivation: Lower Cost* | yes | Switzerland | 12.5328 | 0.1059 | 36 | 393 | 160 | 0.0004 | 0.0130 | 0.6395 | NE | M |
| *Baking Motivation: Lower Cost* | yes | Hungary | 9.8135 | 0.0937 | 6 | 12 | 160 | 0.0017 | 0.0385 | 3.4906 | PE | M |
| *Baking Motivation: Lower Cost* | yes | Slovenia | 8.3928 | 0.0867 | 6 | 13 | 160 | 0.0038 | 0.0503 | 3.2221 | PE | M |
| *Baking Motivation: Lower Cost* | yes | Sweden | 8.0671 | 0.0850 | 9 | 25 | 160 | 0.0045 | 0.0547 | 2.5133 | PE | M |
| *Baking Motivation: Lower Cost* | yes | Finland | 6.7859 | 0.0779 | 31 | 142 | 160 | 0.0092 | 0.0876 | 1.5241 | PE | M |
| *Baking Motivation: Other Reasons* | yes | Italy | 7.6495 | 0.0828 | 6 | 81 | 26 | 0.0057 | 0.0606 | 3.1823 | PE | M |
| *Baking Motivation: Sustainability* | yes | Switzerland | 9.3261 | 0.0914 | 14 | 393 | 20 | 0.0023 | 0.0431 | 1.9896 | PE | M |
| *Baking Motivation: Taste* | yes | Romania | 9.0583 | 0.0901 | 28 | 46 | 888 | 0.0026 | 0.0450 | 0.7657 | NE | M |
| *Baking Motivation: Taste* | yes | Italy | 6.4603 | 0.0761 | 55 | 81 | 888 | 0.0110 | 0.0935 | 0.8541 | NE | M |
| *Baking Motivation: Taste* | yes | Czechia | 6.3181 | 0.0752 | 3 | 8 | 888 | 0.0120 | 0.0982 | 0.4717 | NE | M |
| *Baking Motivation: Taste* | yes | Finland | 5.5745 | 0.0706 | 124 | 142 | 888 | 0.0182 | 0.1338 | 1.0984 | PE | M |
| *Baking Motivation: Taste* | yes | Belgium | 5.2624 | 0.0686 | 80 | 113 | 888 | 0.0218 | 0.1499 | 0.8905 | NE | M |
| *Baking Motivation: Tradition* | yes | Czechia | 7.0315 | 0.0793 | 3 | 8 | 80 | 0.0080 | 0.0792 | 5.2359 | PE | M |

| | | | | | | | | | | | | |
|---|---|---|---|---|---|---|---|---|---|---|---|---|
| *Baking Motivation: Tradition* | yes | Finland | 4.8617 | 0.0660 | 17 | 142 | 80 | 0.0275 | 0.1705 | 1.6716 | PE | M |
| *Baking Motivation: Tradition* | yes | Sweden | 4.5176 | 0.0636 | 5 | 25 | 80 | 0.0335 | 0.2013 | 2.7925 | PE | M |
| *Perceived Benefit: Better Digestibility* | yes | Romania | 8.6579 | 0.0880 | 32 | 46 | 528 | 0.0033 | 0.0470 | 1.4717 | PE | M |
| *Perceived Benefit: Better Digestibility* | yes | Denmark | 4.9304 | 0.0664 | 3 | 17 | 528 | 0.0264 | 0.1705 | 0.3733 | NE | M |
| *Perceived Benefit: Better Digestibility* | yes | Italy | 4.1245 | 0.0608 | 29 | 81 | 528 | 0.0423 | 0.2252 | 0.7574 | NE | M |
| *Perceived Benefit: Gut Health* | yes | Germany | 13.7568 | 0.1110 | 5 | 158 | 140 | 0.0002 | 0.0111 | 0.2525 | NE | M |
| *Perceived Benefit: Gut Health* | yes | Romania | 9.3801 | 0.0916 | 13 | 46 | 140 | 0.0022 | 0.0431 | 2.2548 | PE | M |
| *Perceived Benefit: Gut Health* | yes | United Kingdom | 4.9729 | 0.0667 | 5 | 14 | 140 | 0.0257 | 0.1705 | 2.8495 | PE | M |
| *Perceived Benefit: Gut Health* | yes | Sweden | 4.2302 | 0.0615 | 7 | 25 | 140 | 0.0397 | 0.2164 | 2.2340 | PE | M |
| *Perceived Benefit: Healthier Ingredients* | yes | Switzerland | 11.0500 | 0.0995 | 25 | 393 | 119 | 0.0009 | 0.0237 | 0.5971 | NE | M |
| *Perceived Benefit: Healthier Ingredients* | yes | Germany | 8.8130 | 0.0888 | 28 | 158 | 119 | 0.0030 | 0.0456 | 1.6634 | PE | M |
| *Perceived Benefit: Healthier Ingredients* | yes | France | 7.6486 | 0.0827 | 7 | 23 | 119 | 0.0057 | 0.0606 | 2.8568 | PE | M |
| *Perceived Benefit: Increased Satiety* | yes | France | 4.4023 | 0.0628 | 3 | 23 | 36 | 0.0359 | 0.2083 | 4.0471 | PE | M |
| *Perceived Benefit: Intolerance/Allergy* | yes | Switzerland | 4.3247 | 0.0622 | 36 | 393 | 77 | 0.0376 | 0.2130 | 1.3288 | PE | M |
| *Perceived Benefit: Mental Health* | yes | Denmark | 4.9009 | 0.0662 | 3 | 17 | 46 | 0.0268 | 0.1705 | 4.2852 | PE | M |
| *Perceived Benefit: Nutrient Availability* | yes | Sweden | 8.9169 | 0.0893 | 9 | 25 | 153 | 0.0028 | 0.0456 | 2.6282 | PE | M |
| *Perceived Benefit: Nutrient Availability* | yes | Austria | 8.1248 | 0.0853 | 9 | 26 | 153 | 0.0044 | 0.0547 | 2.5271 | PE | M |
| *Perceived Benefit: Nutrient Availability* | yes | Italy | 6.4952 | 0.0763 | 3 | 81 | 153 | 0.0108 | 0.0935 | 0.2704 | NE | M |
| *Perceived Benefit: Nutrient Availability* | yes | Switzerland | 5.9016 | 0.0727 | 40 | 393 | 153 | 0.0151 | 0.1171 | 0.7431 | NE | M |
| *Perceived Benefit: Stable Glucose Levels* | yes | Romania | 21.9975 | 0.1403 | 9 | 46 | 50 | 0.0000 | 0.0003 | 4.3709 | PE | M |
| *Perceived Health Benefits* | yes | Italy | 12.4676 | 0.1083 | 55 | 79 | 895 | 0.0004 | 0.0130 | 0.8269 | NE | M |
| *Perceived Health Benefits* | yes | Germany | 6.5616 | 0.0786 | 116 | 151 | 895 | 0.0104 | 0.0935 | 0.9124 | NE | M |
| *Perceived Health Benefits* | yes | Romania | 3.8853 | 0.0605 | 44 | 46 | 895 | 0.0487 | 0.2501 | 1.1361 | PE | M |
| *Perceived Health Benefits* | yes | Austria | 3.8594 | 0.0603 | 26 | 26 | 895 | 0.0495 | 0.2516 | 1.1877 | PE | M |
| *Self-Reported Sourdough Skill (Years)* | <0.5 | Germany | 11.1784 | 0.1021 | 3 | 153 | 104 | 0.0008 | 0.0237 | 0.2023 | NE | M |

| | | | | | | | | | | | | |
|---|---|---|---|---|---|---|---|---|---|---|---|---|
| *Self-Reported Sourdough Skill (Years)* | <0.5 | Switzerland | 5.2984 | 0.0703 | 48 | 380 | 104 | 0.0213 | 0.1499 | 1.3032 | PE | M |
| *Self-Reported Sourdough Skill (Years)* | >5 | Germany | 21.3039 | 0.1409 | 66 | 153 | 294 | 0.0000 | 0.0003 | 1.5744 | PE | M |
| *Self-Reported Sourdough Skill (Years)* | >5 | Sweden | 9.1046 | 0.0921 | 14 | 25 | 294 | 0.0025 | 0.0450 | 2.0438 | PE | M |
| *Self-Reported Sourdough Skill (Years)* | >5 | Switzerland | 7.8843 | 0.0857 | 84 | 380 | 294 | 0.0050 | 0.0592 | 0.8068 | NE | M |
| *Self-Reported Sourdough Skill (Years)* | >5 | Finland | 5.3098 | 0.0703 | 27 | 142 | 294 | 0.0212 | 0.1499 | 0.6939 | NE | M |
| *Self-Reported Sourdough Skill (Years)* | >5 | Italy | 3.9015 | 0.0603 | 30 | 80 | 294 | 0.0482 | 0.2501 | 1.3686 | PE | M |
| *Self-Reported Sourdough Skill (Years)* | 0.5 to 2 | Norway | 10.5138 | 0.0990 | 3 | 3 | 167 | 0.0012 | 0.0301 | 6.4251 | PE | M |
| *Self-Reported Sourdough Skill (Years)* | 0.5 to 2 | Germany | 10.2803 | 0.0979 | 10 | 153 | 167 | 0.0013 | 0.0326 | 0.4199 | NE | M |
| *Self-Reported Sourdough Skill (Years)* | 0.5 to 2 | Netherlands | 4.5965 | 0.0655 | 8 | 24 | 167 | 0.0320 | 0.1966 | 2.1417 | PE | M |
| *Self-Reported Sourdough Skill (Years)* | 2 to 5 | Finland | 22.3861 | 0.1444 | 83 | 142 | 429 | 0.0000 | 0.0003 | 1.4619 | PE | M |
| *Self-Reported Sourdough Skill (Years)* | 2 to 5 | Sweden | 7.2004 | 0.0819 | 3 | 25 | 429 | 0.0073 | 0.0748 | 0.3001 | NE | M |
| *Self-Reported Sourdough Skill (Years)* | 2 to 5 | Romania | 4.4952 | 0.0647 | 11 | 46 | 429 | 0.0340 | 0.2017 | 0.5981 | NE | M |
| *Self-Reported Sourdough Skill (Years)* | professional | Czechia | 13.3896 | 0.1117 | 3 | 5 | 79 | 0.0003 | 0.0123 | 8.1494 | PE | M |
| *Self-Reported Sourdough Skill (Years)* | professional | Switzerland | 6.6145 | 0.0785 | 39 | 380 | 79 | 0.0101 | 0.0935 | 1.3940 | PE | M |

**Supplementary Table 10.** Chi-square enrichment test results for sourdough storage and home/bakery environments across countries. p-values were corrected using the Benjamini–Hochberg false discovery rate (BH-FDR). Indicator codes: PE = positive enrichment, NE = negative enrichment, M = multiple indicators, U = unique indicator.

| ***Feature*** | ***Category*** | ***Cluster*** | ***Chi2 Statistic*** | ***Cramér's V*** | ***Cluster Count with Category*** | ***Cluster Total*** | ***Category Total*** | ***Raw p-value*** | ***Corrected p-value*** | ***Enrichment Score*** | ***Interpretation*** | ***Indicator*** |
|---|---|---|---|---|---|---|---|---|---|---|---|---|
| *Dough Storage Container* | glass jar | United Kingdom | 5.2668 | 0.0691 | 8 | 14 | 919 | 0.0217 | 0.1261 | 0.6852 | NE | M |
| *Dough Storage Container* | glass jar | Finland | 4.8135 | 0.0661 | 128 | 142 | 919 | 0.0282 | 0.1551 | 1.0809 | PE | M |
| *Dough Storage Container* | other | United Kingdom | 19.3020 | 0.1323 | 3 | 14 | 21 | 0.0000 | 0.0004 | 11.2449 | PE | M |
| *Dough Storage Container* | plastic container | Finland | 6.9390 | 0.0794 | 10 | 142 | 162 | 0.0084 | 0.0756 | 0.4790 | NE | U |
| *Dough Storage Cover* | closed lid | Finland | 20.4713 | 0.1363 | 102 | 142 | 593 | 0.0000 | 0.0003 | 1.3349 | PE | M |
| *Dough Storage Cover* | closed lid | Switzerland | 15.2300 | 0.1176 | 179 | 391 | 593 | 0.0001 | 0.0020 | 0.8508 | NE | M |

| | | | | | | | | | | | | |
|---|---|---|---|---|---|---|---|---|---|---|---|---|
| *Dough Storage Cover* | closed lid | Germany | 12.4660 | 0.1064 | 106 | 158 | 593 | 0.0004 | 0.0068 | 1.2467 | PE | M |
| *Dough Storage Cover* | fabric | Italy | 22.3307 | 0.1424 | 9 | 81 | 28 | 0.0000 | 0.0001 | 4.3730 | PE | M |
| *Dough Storage Cover* | fabric | Netherlands | 6.1456 | 0.0747 | 3 | 24 | 28 | 0.0132 | 0.0974 | 4.9196 | PE | M |
| *Dough Storage Cover* | loose lid | Finland | 13.4132 | 0.1103 | 33 | 142 | 413 | 0.0002 | 0.0044 | 0.6201 | NE | M |
| *Dough Storage Cover* | loose lid | Switzerland | 11.4041 | 0.1017 | 173 | 391 | 413 | 0.0007 | 0.0117 | 1.1806 | PE | M |
| *Dough Storage Cover* | loose lid | Denmark | 6.7071 | 0.0780 | 12 | 17 | 413 | 0.0096 | 0.0845 | 1.8835 | PE | M |
| *Dough Storage Cover* | loose lid | Italy | 4.4607 | 0.0636 | 21 | 81 | 413 | 0.0347 | 0.1727 | 0.6918 | NE | M |
| *Dough Storage Cover* | other | Italy | 4.2603 | 0.0622 | 5 | 81 | 25 | 0.0390 | 0.1885 | 2.7210 | PE | M |
| *Dough Storage Cover* | plastic wrap | Switzerland | 7.1833 | 0.0807 | 24 | 391 | 43 | 0.0074 | 0.0672 | 1.5731 | PE | M |
| *Dough Storage Cover* | plastic wrap | Austria | 6.4896 | 0.0767 | 4 | 26 | 43 | 0.0109 | 0.0922 | 3.9428 | PE | M |
| *Dough Storage Cover* | plastic wrap | Italy | 3.9628 | 0.0600 | 7 | 81 | 43 | 0.0465 | 0.2029 | 2.2148 | PE | M |
| *Feeding Storage Location* | dough corner in bakery | Germany | 18.0757 | 0.1281 | 5 | 158 | 6 | 0.0000 | 0.0006 | 5.8122 | PE | U |
| *Feeding Storage Location* | fridge/cool place | Italy | 14.1107 | 0.1132 | 18 | 81 | 107 | 0.0002 | 0.0031 | 2.2887 | PE | M |
| *Feeding Storage Location* | fridge/cool place | Germany | 5.1817 | 0.0686 | 7 | 158 | 107 | 0.0228 | 0.1282 | 0.4563 | NE | M |
| *Feeding Storage Location* | kitchen | Germany | 31.5897 | 0.1693 | 66 | 158 | 685 | 0.0000 | 0.0000 | 0.6720 | NE | M |
| *Feeding Storage Location* | kitchen | Belgium | 7.8789 | 0.0846 | 77 | 102 | 685 | 0.0050 | 0.0493 | 1.2145 | PE | M |
| *Feeding Storage Location* | kitchen | Hungary | 5.8486 | 0.0729 | 12 | 12 | 685 | 0.0156 | 0.1038 | 1.6088 | PE | M |
| *Feeding Storage Location* | other | Italy | 5.9005 | 0.0732 | 6 | 81 | 29 | 0.0151 | 0.1022 | 2.8148 | PE | M |
| *Feeding Storage Location* | other | Finland | 4.4678 | 0.0637 | 8 | 142 | 29 | 0.0345 | 0.1727 | 2.1408 | PE | M |
| *Feeding Storage Location* | proofing chamber | Germany | 99.3663 | 0.3003 | 53 | 158 | 117 | 0.0000 | 0.0000 | 3.1595 | PE | M |
| *Feeding Storage Location* | proofing chamber | Finland | 9.5285 | 0.0930 | 4 | 142 | 117 | 0.0020 | 0.0270 | 0.2653 | NE | M |
| *Feeding Storage Location* | proofing chamber | Spain | 4.6758 | 0.0651 | 3 | 7 | 117 | 0.0306 | 0.1635 | 4.0366 | PE | M |
| *Feeding Storage Location* | proofing chamber | Belgium | 4.5609 | 0.0643 | 4 | 102 | 117 | 0.0327 | 0.1662 | 0.3694 | NE | M |
| *Feeding Storage Location* | warm place | Finland | 4.3619 | 0.0629 | 29 | 142 | 158 | 0.0368 | 0.1812 | 1.4244 | PE | U |

| | | | | | | | | | | | | |
|---|---|---|---|---|---|---|---|---|---|---|---|---|
| *Fermented Acidic Beverages* | yes | United Kingdom | 8.5751 | 0.0876 | 6 | 14 | 146 | 0.0034 | 0.0373 | 3.2789 | PE | M |
| *Fermented Acidic Beverages* | yes | Belgium | 5.1778 | 0.0681 | 23 | 113 | 146 | 0.0229 | 0.1282 | 1.5572 | PE | M |
| *Fermented Acidic Beverages* | yes | Finland | 4.6133 | 0.0643 | 10 | 142 | 146 | 0.0317 | 0.1635 | 0.5388 | NE | M |
| *Fermented Alcoholic Beverages* | yes | Slovenia | 5.4213 | 0.0697 | 3 | 13 | 57 | 0.0199 | 0.1182 | 4.5223 | PE | M |
| *Fermented Alcoholic Beverages* | yes | Sweden | 4.1801 | 0.0612 | 4 | 25 | 57 | 0.0409 | 0.1920 | 3.1354 | PE | M |
| *Fermented Dairy Products* | yes | Germany | 20.1523 | 0.1343 | 54 | 158 | 229 | 0.0000 | 0.0003 | 1.6671 | PE | M |
| *Fermented Dairy Products* | yes | Finland | 19.0396 | 0.1306 | 9 | 142 | 229 | 0.0000 | 0.0004 | 0.3092 | NE | M |
| *Fermented Dairy Products* | yes | France | 16.5063 | 0.1216 | 13 | 23 | 229 | 0.0000 | 0.0011 | 2.7570 | PE | M |
| *Fermented Dairy Products* | yes | Austria | 9.1967 | 0.0907 | 12 | 26 | 229 | 0.0024 | 0.0292 | 2.2513 | PE | M |
| *Fermented Vegetables/Fruits* | yes | Finland | 14.8221 | 0.1152 | 23 | 142 | 340 | 0.0001 | 0.0022 | 0.5321 | NE | M |
| *Fermented Vegetables/Fruits* | yes | Hungary | 9.3482 | 0.0915 | 9 | 12 | 340 | 0.0022 | 0.0282 | 2.4640 | PE | M |
| *Fermented Vegetables/Fruits* | yes | United Kingdom | 6.1376 | 0.0741 | 9 | 14 | 340 | 0.0132 | 0.0974 | 2.1120 | PE | M |
| *Fermented Vegetables/Fruits* | yes | Austria | 5.8030 | 0.0721 | 14 | 26 | 340 | 0.0160 | 0.1038 | 1.7690 | PE | M |
| *Fermented Vegetables/Fruits* | yes | Norway | 3.9748 | 0.0597 | 3 | 3 | 340 | 0.0462 | 0.2029 | 3.2853 | PE | M |
| *Other Fermentation Types* | yes | Italy | 6.1799 | 0.0744 | 8 | 81 | 45 | 0.0129 | 0.0974 | 2.4516 | PE | M |
| *Other Fermentation Types* | yes | Switzerland | 4.0718 | 0.0604 | 9 | 393 | 45 | 0.0436 | 0.1985 | 0.5684 | NE | M |
| *Other Fermentations* | yes | Finland | 35.7845 | 0.1819 | 35 | 142 | 523 | 0.0000 | 0.0000 | 0.5095 | NE | M |
| *Other Fermentations* | yes | Hungary | 7.4356 | 0.0829 | 11 | 12 | 523 | 0.0064 | 0.0595 | 1.8947 | PE | M |
| *Other Fermentations* | yes | Sweden | 4.7894 | 0.0666 | 18 | 25 | 523 | 0.0286 | 0.1551 | 1.4882 | PE | M |
| *Other Fermentations* | yes | Germany | 4.0156 | 0.0609 | 86 | 153 | 523 | 0.0451 | 0.2002 | 1.1618 | PE | M |
| *Ownership of Animals (No Hair)* | yes | Sweden | 7.9960 | 0.0846 | 3 | 25 | 23 | 0.0047 | 0.0472 | 5.8278 | PE | M |
| *Ownership of Cat* | yes | Belgium | 16.0826 | 0.1200 | 41 | 113 | 237 | 0.0001 | 0.0013 | 1.7101 | PE | M |
| *Ownership of Cat* | yes | Germany | 7.4804 | 0.0818 | 20 | 158 | 237 | 0.0062 | 0.0591 | 0.5966 | NE | M |
| *Ownership of Cat* | yes | France | 5.6684 | 0.0712 | 10 | 23 | 237 | 0.0173 | 0.1051 | 2.0492 | PE | M |
| *Ownership of Dog* | yes | Germany | 5.9372 | 0.0729 | 15 | 158 | 184 | 0.0148 | 0.1020 | 0.5763 | NE | M |

| | | | | | | | | | | | | |
|---|---|---|---|---|---|---|---|---|---|---|---|---|
| *Ownership of Dog* | yes | Spain | 5.7548 | 0.0718 | 4 | 7 | 184 | 0.0164 | 0.1039 | 3.4689 | PE | M |
| *Pet Ownership* | yes | Belgium | 19.3977 | 0.1327 | 57 | 102 | 391 | 0.0000 | 0.0004 | 1.5736 | PE | M |
| *Pet Ownership* | yes | Germany | 10.0073 | 0.0953 | 38 | 158 | 391 | 0.0016 | 0.0226 | 0.6772 | NE | M |
| *Pet Ownership* | yes | Spain | 5.7032 | 0.0720 | 6 | 7 | 391 | 0.0169 | 0.1051 | 2.4136 | PE | M |
| *Pet Ownership* | yes | Netherlan ds | 4.6069 | 0.0647 | 14 | 24 | 391 | 0.0318 | 0.1635 | 1.6426 | PE | M |
| *Presence of Kids* | yes | Belgium | 8.6006 | 0.0890 | 50 | 101 | 388 | 0.0034 | 0.0373 | 1.3869 | PE | M |
| *Presence of Kids* | yes | Germany | 5.9466 | 0.0740 | 42 | 157 | 388 | 0.0147 | 0.1020 | 0.7495 | NE | M |
| *Presence of Kids* | yes | Czechia | 4.0608 | 0.0611 | 5 | 6 | 388 | 0.0439 | 0.1985 | 2.3346 | PE | M |
| *Presence of Plants* | yes | Switzerlan d | 26.4896 | 0.1560 | 157 | 387 | 557 | 0.0000 | 0.0000 | 0.7924 | NE | M |
| *Presence of Plants* | yes | Sweden | 9.7183 | 0.0945 | 21 | 25 | 557 | 0.0018 | 0.0250 | 1.6408 | PE | M |
| *Presence of Plants* | yes | Belgium | 8.3105 | 0.0874 | 66 | 101 | 557 | 0.0039 | 0.0413 | 1.2764 | PE | M |
| *Presence of Plants* | yes | Germany | 6.1794 | 0.0754 | 65 | 156 | 557 | 0.0129 | 0.0974 | 0.8139 | NE | M |
| *Presence of Plants* | yes | Finland | 4.2444 | 0.0625 | 83 | 139 | 557 | 0.0394 | 0.1885 | 1.1664 | PE | M |
| *Storage After Feeding* | fridge | Finland | 40.4009 | 0.1933 | 84 | 142 | 860 | 0.0000 | 0.0000 | 0.7436 | NE | M |
| *Storage After Feeding* | fridge | Switzerlan d | 18.6984 | 0.1315 | 331 | 381 | 860 | 0.0000 | 0.0004 | 1.0920 | PE | M |
| *Storage After Feeding* | fridge | Germany | 6.4952 | 0.0775 | 134 | 153 | 860 | 0.0108 | 0.0922 | 1.1009 | PE | M |
| *Storage After Feeding* | keep at rt | Finland | 34.9822 | 0.1799 | 53 | 142 | 204 | 0.0000 | 0.0000 | 1.9778 | PE | M |
| *Storage After Feeding* | keep at rt | Switzerlan d | 13.2838 | 0.1109 | 49 | 381 | 204 | 0.0003 | 0.0046 | 0.6815 | NE | M |
| *Storage After Feeding* | keep at rt | Germany | 6.4325 | 0.0771 | 17 | 153 | 204 | 0.0112 | 0.0936 | 0.5888 | NE | M |
| *Storage After Feeding* | keep at rt | Czechia | 6.1369 | 0.0753 | 4 | 6 | 204 | 0.0132 | 0.0974 | 3.5327 | PE | M |

**Supplementary list 1.** Deduplicated and curated list of the main questions and interests obtained from the participants to the co-design workshops across Europe.

- What are the health effects of using sourdough?
- Does the microbial composition of a sourdough change over time?
- Why does sourdough sometimes go bad, which microbes can do it?
- Does the method of storing a sourdough influence it?
- Is there a difference between sourdoughs made with additive-free flour and normal flour?
- Can you change or regulate the taste and function of a sourdough?
- Do different sourdoughs have different microbes?
- Does a change in cereal flour type have an effect on the sourdough?
- Is filtered water okay for use in a sourdough?
- What is the optimal feeding frequency?
- How long can you leave your sourdough without feeding?
- What are the influences of the addition of different ingredients?
- How much does a starter vary between different timepoints? When is it stable?
- Which microorganisms are the most useful in my sourdough and can I selectively enrich them in some way?
- Does the air quality/ pollution influence a starter more than the general geolocation?
- Most important influences on the sourdough?
- Which microorganisms are present and is there a link with the conditions?
- Does my sourdough contain rare microorganisms?
- What is the link between a sourdough and the sensorial properties of the bread?
- How acidic is my sourdough? Which compounds are present? Are there unique ones, absences, health benefits related to some,...
- Is a professional sourdough starter different? Is it easier to maintain?
- Is it safe to eat a sourdough starter without baking it?
- Is it possible to use gut-bacteria to inoculate a sourdough starter?
- Is the microbial activity of the sourdough affecting the dough behaviour and the bread?
- Can you use gluten-free flour to bake sourdough bread?
- Does the taste of the bread change depending on the acids that are produced in the sourdough?
